# Art’s Hidden Topology: A window into human perception

**DOI:** 10.1101/2024.10.16.618741

**Authors:** Emil Dmitruk, Beata Bajno, Joanna Dreszer, Bibianna Bałaj, Ewa Ratajczak, Marcin Hajnowski, Romuald A. Janik, Marek Kuś, Shabnam N. Kadir, Jacek Rogala

**Affiliations:** Biocomputation Research Group, University of Hertfordshire, Hatfield, United Kingdom!; Independent artist, Poland; Institute of Psychology Faculty of Philosophy and Social Sciences, Nicolaus Copernicus University in Toruń Gagarina 39, 87-100 Torun, Poland; Institute of Theoretical Physics and Mark Kac Center for Complex Systems Research, Jagiellonian University, Łojasiewicza 11, 30-348 Kraków, Poland; Center for Theoretical Physics, Polish Academy of Sciences, Poland; Centre for Research on Culture, Language, and Mind, University of Warsaw, Warsaw, Poland; Centre for Systemic Risk Analysis, University of Warsaw, Warsaw, Poland

## Abstract

Generations of researchers have sought a link between features of an artistic image and the audience’s experience. However, a direct link between the properties of an image and the responses evoked has still not been established. Given the importance of shape to human perception and artistic creation, it can be assumed that one of the most important aspects of an artistic image is the use of different visual structures. We show that a method from the field of computational topology, persistent homology, can be used to analyse properties of image structures and composition at multiple scales. In order to determine the reliability of this method as a tool for analysing visual artworks, we analysed two different sets of abstract paintings that revealed significant discrepancies in the eye tracking and electroencephalography (EEG) activity of viewers. Our research showed that our newly developed method using persistent homology, not only clearly distinguished between two sets of images, which was not possible with common statistical image properties, but also allowed us to map topological features onto gaze fixation heat maps.

## 1 Introduction

Understanding the complex interaction between art and human perception has long intrigued researchers. During the last 150 years, there have been many attempts to use scientific methods to understand the human aesthetic experience [Fechner, 1876, Bense, 1969, Kawabata and Zeki, 2004, Leder et al., 2004, Silvia, 2006, Vessel et al., 2012]. In visual art, the mainstream of this research path has focused, and continues to focus, on the relationship between properties of images and both the subjective human experience and objective measurable neural responses to viewing the images [Kawabata and Zeki, 2004, Rigau et al., 2008, Umilta’ et al., 2012, Menzel et al., 2018]. However, although it is acknowledged that works of art often reflect the statistical regularities present in natural scenes, a direct correlation between these statistics and the experience evoked by art is weak or at most, moderate [Mather, 2018].

Concurrently, studies in both human and non-human primates (e.g. macaques) have revealed that higher-order brain regions are more responsive to holistic shapes than to basic visual elements such as edges used to define the shape [Śary et al., 1993, Vogels and Orban, 1996,Grill-Spector et al., 2001,Malach et al., 1995]. Specifically, the investigations with humans show that these brain regions respond more strongly to objects defined from either luminance, texture, illusory contours, motion, stereo, or colour than to random stimuli or uniform textures generated from the same primary cues [Appelbaum et al., 2006, Gilaie-Dotan et al., 2002, Grill-Spector et al., 1998, Kastner et al., 2000, Kourtzi and Kanwisher, 2000, Mendola et al., 1999]. Intriguingly, the holistic representation of an object is recognized earlier than its constituent parts [He and Nakayama, 1992,Rensink and Enns, 1998,Hochstein and Ahissar, 2002,Wolfe and Horowitz, 2004, Kok and de Lange, 2014]. This early processing of the object information was also confirmed by numerous neuroimaging studies using both EEG (for a review see [Kaneshiro et al., 2015]) and fMRI (for a review see [Margalit et al., 2016]).

These neural mechanisms, fundamental for recognizing objects and faces [Freud et al., 2019], are operational from as early as six months of age [Emberson et al., 2017]. Notably, their role extends beyond essential survival-related actions to encompass the perception of possible and impossible objects, imagery, and abstract shape representations [Baker and Kellman, 2018]. The ability to perceive and recognise shapes and objects is not only fundamental to our interaction with the world but also crucial for artists in depicting scenes or objects accurately. This skill is particularly important for individuals with advanced drawing and painting abilities. Research by Robles et al. [Robles et al., 2022] demonstrates a significant correlation between the accuracy of shape judgement in skilled individuals and their drawing precision. This suggests that enhanced abilities in realistic drawing are likely linked to a more precise perception of shapes.

Given the importance of shape and object perception in artistic creation, it is reasonable to postulate that these elements are also vital tools for artists in conveying impressions to their audience. This concept is eloquently explored in Arnheim’s seminal work ‘Art and Visual Perception’, where he delves into the role of visual perception in art and how artists use shapes, among other elements, to create specific psychological effects in viewers. He posits that artists are keenly aware of the impact of certain shapes in crafting these effects [Arnheim, 1974].

We believe that one of the most crucial aspects of an artistic image, especially in terms of viewer experience, is indeed the use of various visual structures. The aforementioned neuroscientific studies indicate that the brain perceives shapes in a seemingly qualitative manner before the shapes can be given explicit geometric descriptions. This is why six-month-old babies are nevertheless able to recognise and appreciate objects and faces despite being unable to paint. In mathematics, topology can be considered a quantitative way of studying qualitative properties of shapes that are unchanged under continuous deformation (where bending and stretching are permitted, but cutting and glueing are forbidden). To tackle these questions, we turn to persistent homology [Zomorodian and Carlsson, 2005], an approach that captures the topological properties of an image’s composition on multiple scales (via the computation of homology classes for a hierarchy of combinatorial objects derived from the image).

This mathematical method builds up abstract structures in very much the same way as an artist composing the shapes of a painting from colours, shades and edges. The method explores qualitative questions about spaces: Are there any distinctive shapes, such as connected components, holes, and voids? How many shapes are there? Are there any holes? How many holes are there? It is, therefore, a useful tool that enables us to quantify the qualitative features of an abstract image to which people are drawn. The appeal of focusing on topological properties lies also in their independence from arbitrarily chosen coordinates and metric properties of the objects being perceived. Persistent homology operates on multiple scales and dimensions, by way of enabling not only the detection of less salient, less noticeable structures but also the disclosure of detailed properties of the image not achievable from commonly used statistical image properties. Unlike figurative art, abstract art does not consist of identifiable objects that can be given clear labels; this detailed topological decomposition of the artworks in terms of shapes enables us to analyse how viewers mentally process a composition comprised of abstract shapes.

To determine the reliability of persistent homology as a tool for analysing visual artworks and to investigate the relationship between topological image properties and human experience, we conducted a study in which we held two different art exhibitions for our experiment: one presenting original abstract art by a contemporary visual artist and the second presenting artificially generated pseudo-artistic abstract images. Analysis of audience responses to these exhibitions revealed significant discrepancies in conscious subjective experience (via questionnaires). Further analysis of eye-tracking data and electroencephalography (EEG) activity revealed discrepancies in the physiological processing of art, suggesting different image properties and their differential effect on human experience and brain activity.

To further examine these observations, we compared the topologically derived properties of the digital copies of the images from the two exhibitions and then investigated the relation between topologically derived image properties and the participants’ eye movements. Finally, we contrasted our findings obtained from the use of persistent homology with conventional statistical image properties calculated for the two sets of images. This methodological comparison was aimed at assessing the effectiveness of persistent homology in capturing the complexity of human perception in visual art. Our current study focuses on the results obtained during the laboratory rather than the gallery segment of the experiment.

## 2 Methods

To ascertain whether topologically derived properties can reliably differentiate between artistic and pseudo-artistic images and whether captured differences are related to human eye movement and brain activity, we used two sets of images: abstract artworks (hereinafter referred to in the text as ‘art’ or ‘artistic’) and abstract artificially generated (hereinafter referred to in the text as ‘pseudo-art’ or ‘pseudo-artistic’) images. The pseudo-artistic images were produced by altering a publicly available generative adversarial network (BigGAN).

Within each of the two sessions (separated by a one-week interval), the images were viewed twice: once in an art gallery and once in a laboratory. There were two separate groups of participants: one group was shown only the artistic images, whereas the other group was only shown the pseudo-artistic images. Both groups of participants were under the impression that they were viewing artworks made by a real artist for a solo exhibition. In the following sections, we describe the origin of the artistic images and the method used to produce the pseudo-artistic images. Next, we describe an experimental procedure designed to collect eye-tracking data and EEG. Finally, we describe how the tools from persistent homology were used to examine differences in what participants were looking at or neglecting to look at.

### 2.1 Images and preprocessing

The artistic images used in the laboratory were digital copies of original works on display at the gallery where the first part of the study was conducted. The works were created as a result of an “unrestricted creative process” (as the artist describes it) by Lidia Kot [Kot, 2022], a contemporary Polish visual artist. This process resulted in a collection of 12 colour art prints prepared especially for the artist’s planned solo exhibition. Each artwork had identical dimensions, 50 *cm ×* 100 *cm* (*W × H*).

Examples of the artworks are shown in Fig. 1-A, and digital copies of all original artworks used in the study are available in the supplementary materials (Sup. Fig. Supp.B.1).

**Figure 1:**
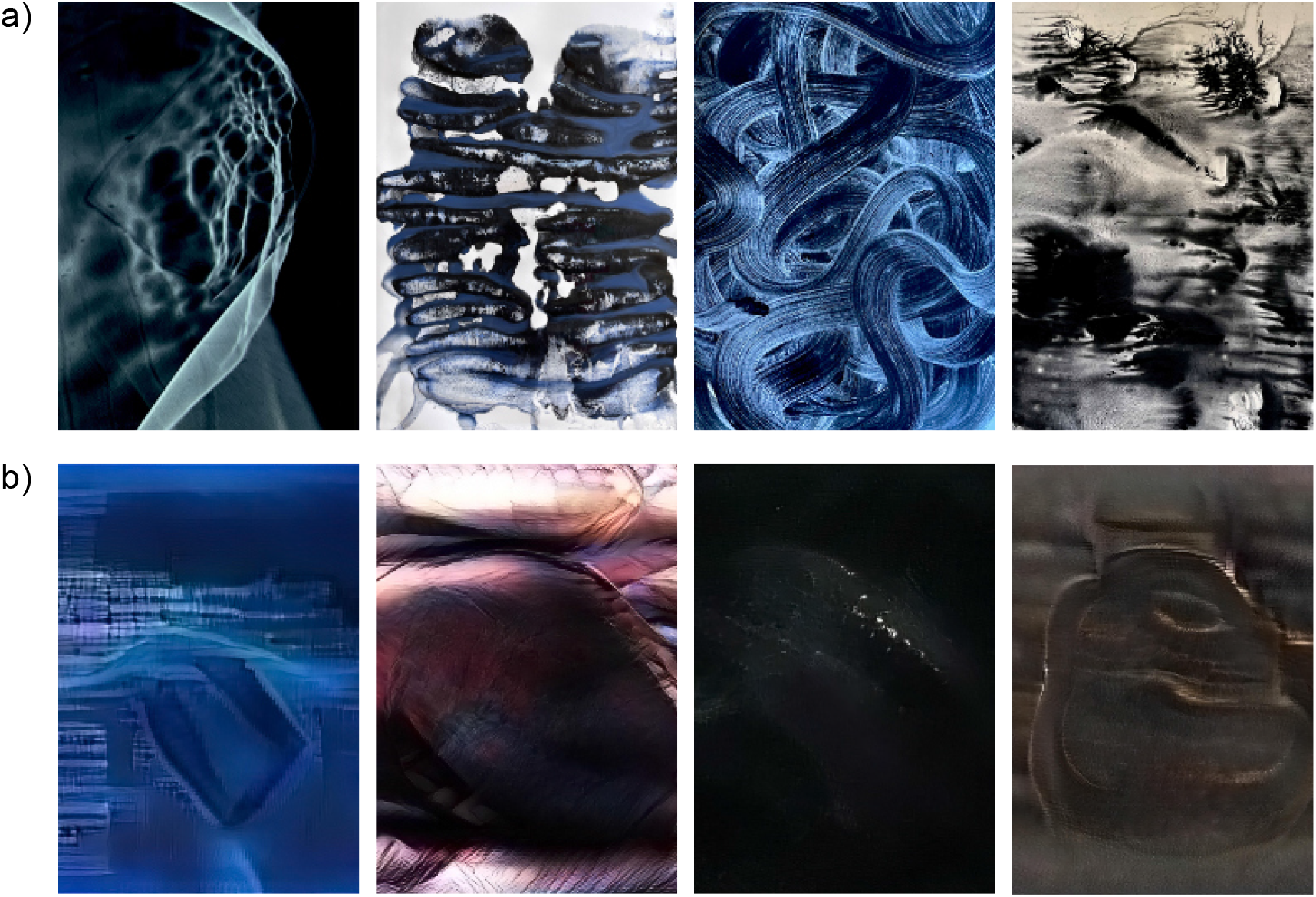
Sample of artistic images [Kot, 2022](a) and pseudo-artistic images (b). The titles, from left to right, are: a) ‘Black holes of memory’ (orig. ‘Czarne dziury pamięci”); ‘Lungs of Blackness’ (orig. ‘Płuca czerni”); ‘Ear of Blackness’ (orig. ‘Ucho czerni’); ‘Guts of Blackness’ (orig. ‘Jelita czerni’). b) ‘Vibrations of time’ (orig. ‘Wibracje czasu’); ‘The Inside’ (orig. ‘Wnętrze’); ‘Cold fire’ (orig. ‘Zimny ognień); ‘Everything is a thing and nothing is everything’.

The pseudo-artistic images were produced using methodology introduced in [Janik, 2023], which involved randomly perturbing certain weights of the generative adversarial neural network BigGAN and using the resulting perturbed neural networks to produce images. The underlying publicly available BigGAN model [Brock et al., 2019] had undergone training on the ImageNet dataset comprising around one million images of the human environment. The BigGAN network is able to produce photorealistic images across 1000 object categories, such as cars, buildings, landscapes, animals, and more. In the present context, it is important to emphasize that the training dataset did not contain any images of art.

In order to ensure that the generated images had an abstract non-figurative character, the weights of all layers in Block 2, as well as those of a layer in the “self-attention” block of the BigGAN network (see Fig. 1 in [Janik, 2023]), were substituted by random numbers. All the remaining weights remained unchanged. In this way, we constructed 500 different neural networks by drawing 500 random vectors of neural network weights, which gave us a sufficient variety of image distortion types or styles. Subsequently, a subset of 480 ImageNet categories was specifically chosen to exclude any that featured animals or human faces to ensure a strictly non-figurative character for the pseudo-artistic images. Then, for each of the 500 network modifications, images were generated using 9 categories randomly selected from this ‘non-figurative’ subset. In this way, we obtained 4500 neural network-generated images of size 256 *×* 256. The images generated by the perturbed neural networks were then enlarged to a 2048 *×* 2048 resolution by first reducing colour noise in Photoshop (Adobe Systems Inc., Adobe Photoshop 2022) and subsequently upscaling using an SRGAN [Ledig et al., 2017] superresolution neural network. Examples of the generated images are shown in Fig. 1-B, and digital copies of all artificially generated images used in the study are available in the supplementary materials (Sup. Fig. Supp.B.2).

For the purpose of this study, the original artworks were downscaled to the resolution 1434*×*2048 (*W ×H*) using Photoshop. All pseudo-artistic images were rescaled to match the resolution of the artistic images. Next, in order to mitigate differences in audience response arising merely from large disparities in brightness or colour, each of the 4500 artificially generated images was compared with each of the artist’s 12 artworks using Matlab’s (The MathWorks, Inc.) *imabsdiff* function. The *imabsdiff* function compares two images of the same size by computing the absolute value of the difference between the pixel intensities of the two images (resulting in a matrix of absolute pixel value differences). Next, this matrix was averaged to obtain a single value for each pair of compared images (by comparing each of 12 images with each of 4500 generated images, resulting in 54000 comparisons). The averaged differences of all comparisons were then placed in ascending order, and the 12 images with the smallest difference were selected as the set of pseudo-artistic images used in the present study. Selected pseudo-artistic images were then printed to the same size as the artistic ones in the same print shop. The titles for the pseudo-artistic exhibition were generated using original curatorial texts and publicly available painting titles from exhibitions held at the Wozownia Gallery. All texts were created using GPT-3 software (beta.openai.com) and initially generated in English before being translated into Polish via Google Translate. From the pool of generated titles, 12 were randomly selected using Matlab (MathWorks Inc.) *randperm* function, while 1 pseudo-curatorial text was selected for the exhibition. The pseudo-curatorial text was proofread by an artist. Finally, titles were randomly assigned to pseudo-artistic images using the Matlab (MathWorks Inc.) *randi* function. The display order of pseudo-artistic images in the gallery was also drawn using the Matlab (MathWorks Inc.) *randi* function. Each set of images had its own dedicated exhibition open to the public. Both exhibitions ran for the same duration and were held at the same venue. The exhibition of the pseudo-artistic images opened directly after the exhibition of the artist Lidia Kot [Kot, 2022] had ended.

In the laboratory, the raw images that were displayed were represented in the JPEG format, with 8 bits for each RGB channel. For the topological analysis with one-parameter persistent homology, all images were converted to greyscale, where every pixel in greyscale was computed as a weighted average of three colours according to the formula [CCIR, 2011, Holy, 2021]: *Y* = 0.299*R* + 0.587*G* + 0.114*B*, where R, G, and B are the pixel intensities in red, green and blue. This conversion follows the CCIR 601 format originally issued in 1987 by the International Radio Consultative Committee (now the ITU Radiocommunication Sector). All images were displayed on a 21-inch computer screen with a 1920 x 1080 dpi resolution.

### 2.2 Experimental procedure

To verify whether artistic and pseudo-artistic images evoke distinct responses under laboratory and ecological conditions (i.e. in the gallery), we recruited 58 participants from Nicolaus Copernicus University in Toruń, Poland. Participants provided informed consent to take part in the study, which was approved by the local ethics committee at the Nicolaus Copernicus University in Toruń (decision # 16/2021/FT). Participants were either second-year art students or persons of similar age and socio-economic status who declared an interest in visual art. Randomly divided into two equal groups, participants were matched according to gender ratio and age. Each participant in the first group was required to visit Lidia Kot’s [Kot, 2022] exhibition and then immediately proceed to the laboratory. Participants in the second group followed the same procedure but were instructed to visit the second exhibition consisting of pseudo-art images. The participants were required to revisit the exhibition and the laboratory a second time, not less than one week after their initial visit.

Both laboratory visits were made immediately after visits to the exhibition. To capture putative differences in brain activity emerging during the perception of artistic vs. artificially generated pseudo-artistic images, eye-tracking data were collected in the gallery, and simultaneous eye-tracking and EEG data were recorded in the laboratory.

In the gallery, participants were asked first to follow a curated order of viewing the images and then to freely engage with the exhibition. During laboratory visits, they were instructed to observe images displayed on a computer screen whilst their eye movements were tracked and EEG recordings were made. The images were presented in the following scheme: first, the title of an image was presented for two seconds, and then, for 7 seconds, the corresponding image was presented. This was followed by a 2-second mask of white noise, then 2 seconds of grey mask, after which another title and image were presented etc. After recording EEG and eye movements, participants were asked to score each image separately on a 6-item flow questionnaire. Each question was scored using an unlabelled slider (with no ticks). The slider moved over a 5-hidden-point Likert scale, which recorded the participants’ responses. After removing data containing excessive artifacts (both in the EEG and eye-tracking signal), data from 50 individuals (26 in Art group) were finally used for analysis.

### 2.3 Eye tracking recording and data analysis

To capture differences in perception of the two different sets of images in the laboratory, we used two basic parameters of eye movements: average fixation durations and average saccades amplitude. Average fixation duration is a measure of attention. Short fixations indicate implicit processing, whereas longer fixations (typically above 70ms of duration) mark the initiation of perceptual information reception and processing by the brain [Glöckner and Herbold, 2011,Schall and Romano Bergstrom, 2014]. The amplitude of saccades is a measure of visual space exploration. The data were preprocessed using a positional algorithm for the automated correction of vertical drift in eye-tracking data [Carr et al., 2022]. For laboratory eye-tracking measurements, the Eyetracker SMI RED 500 was used. This device operates at a sampling frequency of 500 Hz and employs a 5-point calibration procedure for enhanced precision and accuracy.

In the gallery, participants used a wearable eye-tracker built into glasses. To measure attention, we used the average visual intake duration, equivalent to the average fixation duration in the laboratory setting, and the average saccade duration, equivalent to the average saccade amplitude. Both measures were computed for the areas of images extracted from the movie recorded during the visit and for the time spent by participants individually at each image. In the gallery, we used SMI Eye Tracking Glasses 2 Wireless (SMI ETG 2w), dark pupil eye tracking with a binocular sampling rate of 60 Hz, parallax compensation, and accuracy of 0.5°(declared by the manufacturer). After the glasses had been fitted to the study participant and secured, a 1-point calibration procedure was performed. During the study, scene view was recorded with a scene camera (video resolution: 1280x960p and 24FPS), and audio was recorded with an integrated microphone.

All data were curated for outlier treatments using the winsorizing method introduced by Tukey and McLaughlin [Tukey and McLaughlin, 1963]. That is, any value of a variable above or below 2 standard deviations on each side of the variables’ mean was replaced with the value at plus or minus 2 standard deviations itself and tested for normality using the Shapiro test. Since both laboratory and gallery eye tracking data failed to conform to the normal distribution (Shapiro test *p <* 0.001) the comparison was performed using a non-parametric Kruskal-Wallis ANOVA and post-hoc Dunn test with False Discovery Rate (FDR) correction for multiple comparisons. For other non-parametric comparisons, the Mann-Whitney test was used. The significance of differences in statistical analyses is indicated by the *p*-value resulting from the applied tests. Differences are assumed to be significant when the *p*-value is less than 0.05.

### 2.4 EEG recording and data analysis

To analyse putative differences in brain activity we used brain connectivity measures applied to the electroencephalograph (EEG) signal. Brain connectivity can be used to estimate how different brain regions interact during cognitive tasks and how these interactions change in response to interventions. In our study, we hypothesised that the two sets of images could challenge subjects with different levels of difficulty and engagement causing differences in the organisation of brain connectivity patterns. To infer EEG brain connectivity, we use statistical dependencies between EEG signals from different brain regions to estimate how different regions of the brain communicate with each other, hence we used the weighted phase lag index (WPLI) [Vinck et al., 2011, Hardmeier et al., 2014] calculated for all pairs of electrodes.

For EEG measurements, the 64-channel BrainAmp amplifiers manufactured by Brain Products GmbH were used. This device operates at a sampling frequency of 1000 Hz. Signal pre-processing and analyses were performed using MNE-Python [Gramfort et al., 2013]. The preprocessing procedure included downsampling to 500 Hz and filtering in the 0.2-70 Hz range with a finite impulse response (FIR) filter with a Hamming window automatically adjusted to signal length. The power noise of 50 Hz and its higher harmonics (100 Hz and 150 Hz) were removed using a notch filter. Next, the data were cut into 7-second chunks (epochs) matching the duration of the periods when participants viewed the images. To compensate for the high variability of the EEG and to increase statistical power, we combined recordings from both laboratory visits and restricted the number of electrodes included in analyses to the 32 electrodes that matched the International 10-20 system. Electrodes used in analyses included: Fp1, Fp2, F7, F3, Fz, F4, F8, FC5, FC1, FC2, FC6, T7, C3, Cz, C4, T8, TP9, CP5, CP1, CP2, CP6, TP10, P7, P3, Pz, P4, P8, PO9, O1, Oz, O2 and PO10. The same preprocessing procedure was also applied to resting-state data collected from both groups prior to any gallery or laboratory visit, except we replaced the 7-second epochs collected while viewing the images with segments of the same duration extracted from the resting state. The resting state data were used to verify that the groups did not differ in EEG features even before the study began.

All analyses were performed separately for each of the following canonical EEG bands (delta: 0.5-4Hz, theta: 4-8 Hz, alpha: 8-13 Hz, beta: 13-30 Hz, and gamma: 30-70 Hz). Statistical comparisons of group differences were performed using the mass Mann-Whitney U-tests with FDR correction for multiple comparisons.

### 2.5 Topological analysis

We provide a quick informal introduction to topological data analysis in the context that we use in this paper, namely persistent homology from filtrations of cubical complexes derived from 2D images. For mathematical definitions see Supplementary Section A.1; for a more comprehensive account, consult [Hatcher, 2001, Kaczynski et al., 2004, Otter et al., 2017, Ghrist, 2014].

#### 2.5.1 Persistent homology method

Central to the analysis of topological features in datasets, regardless of their representation (e.g., as an image), are the concepts of a filtration and persistence. To illustrate the construction of a 1-parameter filtration for our application, consider an image composed of pixels with varying shades of grey. A filter that permits only shades above a certain intensity to pass through might inadvertently exclude certain geometric structures formed by darker pixels. By varying the filter’s threshold, some structures may either emerge or vanish. In our example, this visual filtering results in a mathematical filtration - whereby we obtain a sequence of cubical complexes (see Subsection 2.5.2 below) ordered by inclusion, that is, a complex obtained from thresholding at a lower level of the filtration is included in the complex obtained at any higher level of the filtration. The span of filtration parameter values over which a particular topological structure exists is termed its persistence. The lowest parameter value for which a structure exists is called its birth, and the highest, its death. Persistence, therefore, is a structure’s lifetime. Structures characterized by extended persistence periods are deemed the most salient, typically offering distinctive insights into the object under examination. We have just described informally a tool used in computational topology [Ghrist, 2007], persistent homology [Zomorodian and Carlsson, 2005].

When this method is applied to two-dimensional objects such as digital images, each pixel is a 2D elementary cube (or square), and two pivotal topological characteristics come to the forefront (as further described in Section 2.5.2). The first is the number of connected components of the parts of an image that have exceeded the filtration threshold, i.e. discrete, disconnected segments within a given structure which are regions of the same or darker colour, hereinafter referred to as the ‘dimension 0’ or ‘dim 0’ cycles (formally, cycle representatives of homology classes). The second is the number of ‘holes’ present in a designated area e.g., the count of regions of a certain colour entirely encircled by areas of a different colour, hereinafter referred to as the ‘dimension 1’ or ‘dim 1’ cycles. In algebraic topology, these integers correspond to the Betti numbers (formally, ranks of homology groups), denoted *β*_0_ and *β*_1_ respectively, as illustrated in Fig. 2.

**Figure 2:**
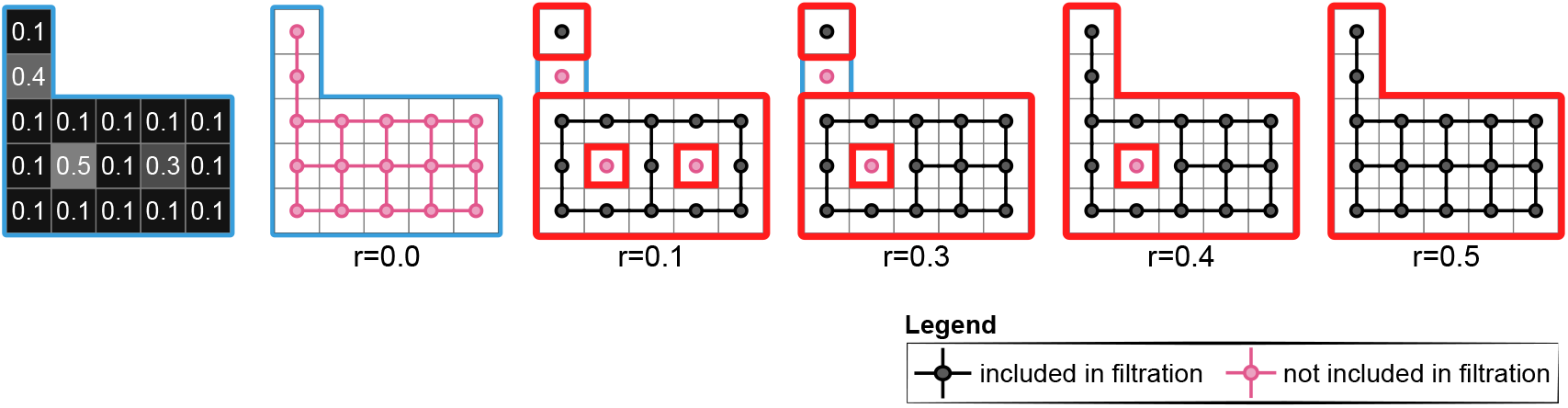
Exploring topology and persistence through filtration. Visualization of a pixel-based structure with varying brightness levels, demonstrating the use of grey intensity (on a scale from 0 to 1) as a filtration parameter. At a grey intensity of 0.1, the structure is divided into two distinct parts, outlined by a red contour) with two ‘holes’ of grey intensities 0.3 and 0.5. This results in Betti numbers *β*_0_ = 2 and *β*_1_ = 2. As the filter transparency (greyness) increases, changes in the visibility of pixels at grey level 0.3, the two distinct shapes are still present, although one of the holes in the bigger part (with intensity r=0.3) disappears, changing the Betti numbers to *β*_0_ = 2 and *β*_1_ = 1. At the intensity r=0.4, the two shapes merged into one part with one hole (*β*_0_ = 1 and *β*_1_ = 1). If we further increase filter intensity to r=0.5 all pixels will become visible yielding Betti numbers *β*_0_ = 1 and *β*_1_ = 0.

Through persistence homology, it is possible to capture the emergence (birth) and disappearance (death) of components and voids relative to filtration values. The evolution of those topological properties is commonly presented in the form of barcodes [Ghrist, 2007]. The barcodes are often transformed into persistence diagrams [Carlsson and Zomorodian, 2009] and persistence landscapes [Bubenik, 2015]. These techniques are elucidated and depicted in Fig. 3, offering graphical insights that effectively highlight variations in topological properties among the objects under scrutiny.

**Figure 3:**
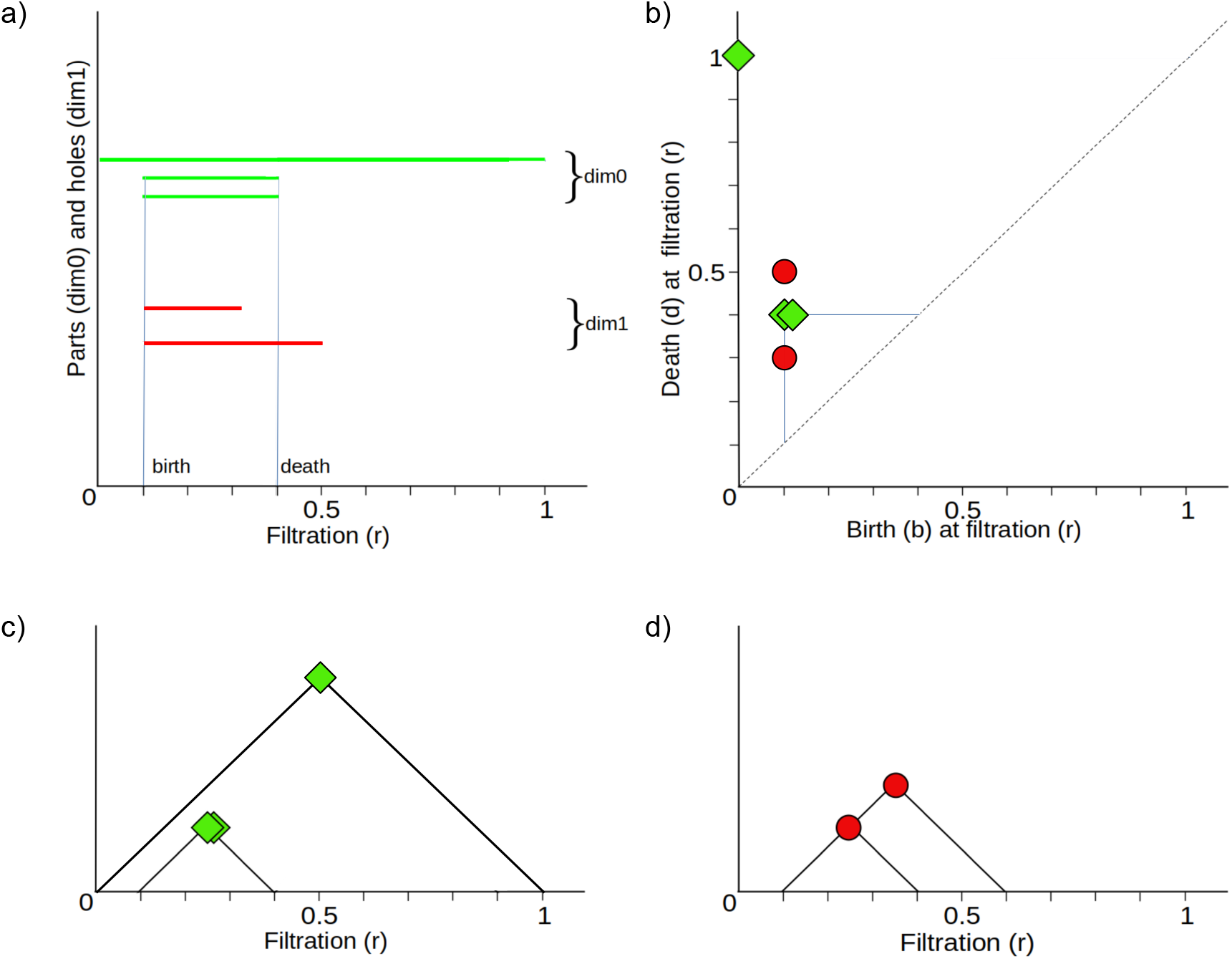
Visualizations of the filtration process. a) ‘barcodes’ - each structure corresponds to a line segment parallel to the axis of the filtration parameter *r*, which begins when the structure appears (i.e. at point *r_b_*, birth) and ends when it disappears (at point *r_d_*, death) marked by two vertical lines for clarity; b) persistence diagram - persistence is also representable via a two-dimensional scatter plot diagram with coordinates (*r_b_, r_d_*) representing births and deaths of the shape’s parts and holes on two orthogonal filtration axes marked by the horizontal blue lines orthogonal to respective axes. Naturally, points on this diagram occupy only the area above the main diagonal, c) and d) persistence landscape - connecting each of the birth/death points to the diagonal with vertical and horizontal lines, as shown for the uppermost point, green point on b, we get a system of ”pyramids” isosceles right triangles. After rotating the persistence diagram by *π/*4, it becomes a persistence landscape drawn separately for dim0 - disconnected parts (Fig. 2c.) and dim1 - holes (Fig. 2d.)

As well as providing valuable graphical representations for qualitative assessment, persistence landscapes (sequences of piecewise linear functions) also enable us to use standard tools from statistics [Bubenik, 2015]. Without a transformation to persistence landscapes, a multiset of barcodes is inadequate for defining even simple notions such as mean and median. In addition to comparing landscapes using L1 distance (see Supplementary Fig. Supp.B.7), we also used simple measures such as the total area beneath the ‘pyramids’ in the persistence landscape (this can also be considered the L1-norm of the persistence landscape). Even though this quick quantification involves some information loss concerning an image’s topological structure, it provided an adequate first glimpse for distinguishing between the two different sets of images in our application.

However, our goal was not only to investigate differences in art pieces - we also investigated whether human perception was guided by the topological properties of the images uncovered via persistent homology. For this purpose, we derived measures that could bind these topological structures with their spatial distribution, namely feature maps. We then investigated their relationship with the gaze-derived heatmaps, as explained in the following sections.

#### 2.5.2 Cubical complexes

An important aspect of the topological investigation is the decision on how to decompose the object of study, i.e. the image, using topological structures. Here, we shall compute persistent homology from a filtration of cubical complexes [Kaczynski et al., 2004, Otter et al., 2017] with the *V − construction* [Bleile et al., 2021].

To obtain a cubical complex, we exploit the fact that the digital representation of an image is a matrix, where the pixels are laid out in a grid, where we take the neighbourhood of a pixel to be limited to the 4 compass directions (some of which are not possible, e.g. the pixels at the periphery of the image). Using this grid structure, a cubical complex is formed as a topological space constructed from a union of vertices and edges, where a vertex is assigned to a pixel, and edges are formed from adjacent pixels which satisfy a condition, e.g. being above (or below) a certain value in the greyness scale as is demonstrated in Fig. 2 and Fig. 4. Each cubical complex thus created (one for each value of the filtration) is dependent on the relationship between neighbouring pixels. Hence, a cubical complex could be, e.g. a group of disjoint pixels located in the image matrix, pixels covering some extent of an image, pixels encircling an area (or hole) containing pixels that are not yet included in the filtration or any combination of the above.

**Figure 4:**
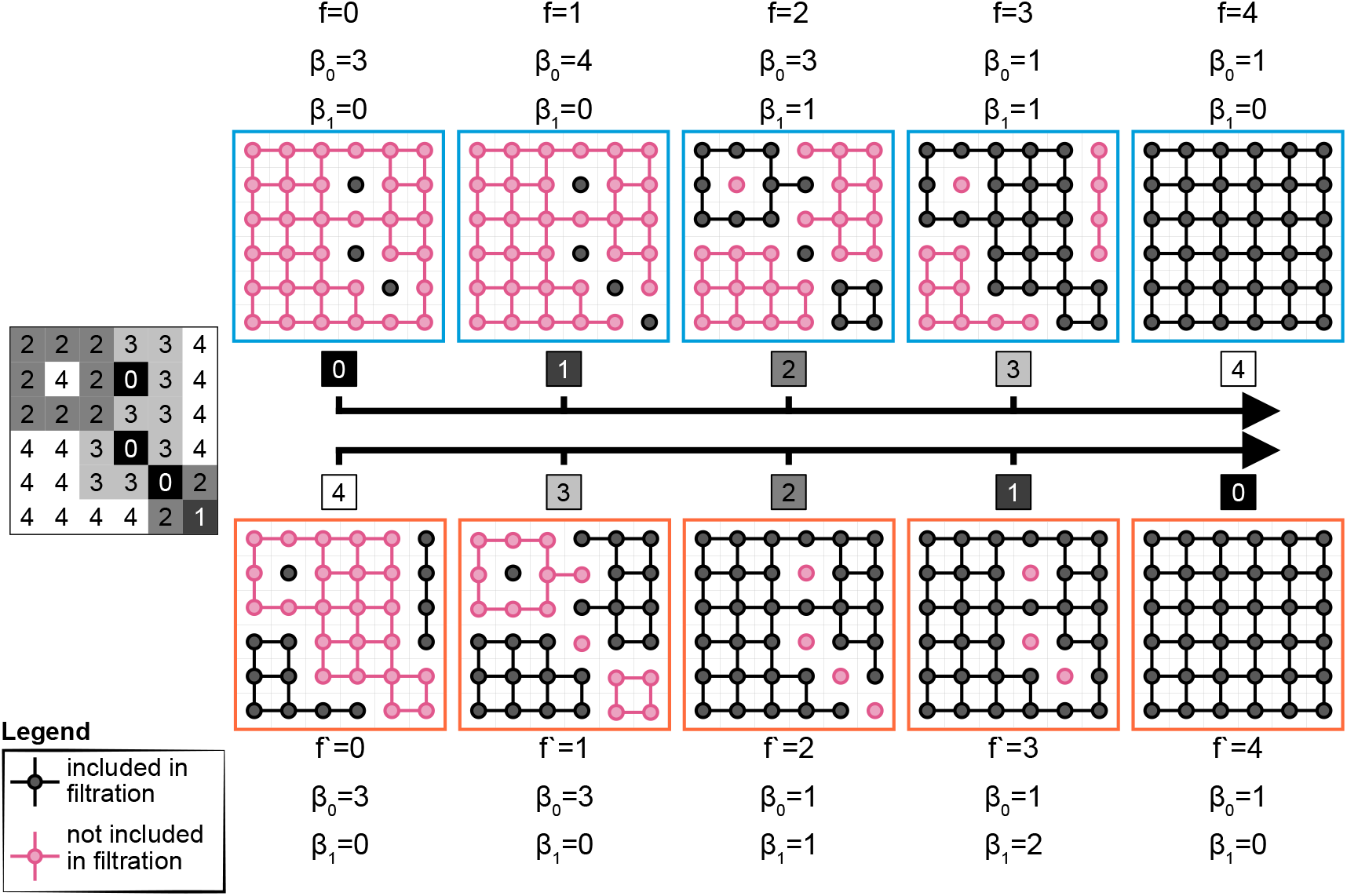
Demonstration of the duality of black-to-white (BW) (first row) and white-to-black filtrations (WB) (second row). The image from which the cubical complexes are derived is to the left of the rows in the middle. Components (and corresponding connections) that comprise a cubical complex at a particular step of the filtration, *f* (or *f ^′^*), are coloured in black, whereas the pink components are not included in the filtration. Pink components also belong to the dual cubical complex coming from the reverse filtration at step *f ^′^* = *M −* 1 *− f*, where *M* + 1 is the total number of filtration steps [0*, M*].

From this, it naturally follows that the topological structures we can investigate are limited to 0-dimensional (connected components) and 1-dimensional (loops) cycles. We record the locations in a filtration where such structures emerge (births) and disappear (deaths). The list of all such births and deaths results in the multi-set of barcodes to which we can associate a persistence landscape (see Fig. 3). More details on obtaining persistence landscapes from cubical complexes derived from image data can be found in the Supplementary Materials A.2. Finally, we also plot Betti curves, as defined in A.2, as a way of showing how the Betti numbers vary throughout the filtration.

Apart from the natural and easily definable transition from image to a nested sequence of cubical complexes, an advantage of this cubical representation of the image is that once the topological content of the image is established, it is possible to study not only the properties of the cycles but also their spatial distribution and spatial extent within an image. This allowed us to compare the images from the exhibition in two ways-by following a standard approach of analysing persistence properties using persistence landscapes and, as introduced in our work, by exploring the spatial distribution of the topological structures using feature maps.

**Duality of persistence homology in cubical complexes on images** Lastly, the persistent homology groups of cubical complexes thus defined exhibit a simple but interesting duality. Topological invariants computed for the filtration starting from black and finishing on white (BW) can be related to invariants computed from the reverse filtration starting from white and finishing on black (WB), namely persistent homology classes of dimension 1 can be (injectively) mapped to dimension 0 classes (connected components) of the reverse filtration. This is similar to results obtained in [Garin et al., 2020, Bleile et al., 2021] and is an example of Alexander duality [Munkres, 1984]. A proof can be easily deduced from looking at Fig. 4; a sketch proof is provided in Supplementary Section A.1.

The advantage of this is that by using information from both filtrations (BW and WB) and only dimension 1 cycles, we capture almost all information about cycles in images-this was utilised in the analysis of feature maps. The only discrepancies are due to cycles which comprise or interact with cycles at the boundary of the image.

Fig. 4 illustrates the duality of filtrations via a simple example. At each filtration step (different steps of the dual filtrations), the underlying grid is partitioned by the connected components (0-cycles of one cubical complex and its complement with respect to the grid, which we shall denote the ‘dual’ complex. Let *f* and *f ^′^* be the steps in the BW and WB filtrations, respectively, where *f ^′^* = *M −* 1 *− f* . For this toy example, the total length of a filtration, *M*, is 4 (for a general greyscale image, *M* = 255). Hence, the black components for *f* = 0 are pink components for *f ^′^* = 3 and vice versa. Hence, as shown in Table 1, for any given pair (*f, f ^′^*) = (*f,* 4 *−* 1 *− f*) the connected components from both filtrations partition the underlying grid; hence the number of partitions of the grid is also the sum of the zeroth Betti numbers.

**Table 1:** Number of partitions for filtration steps for the toy example.

| $f$ | $f'$ | #partitions | $\beta_0^f$ | $\beta_0^{f'}$ |
| --- | --- | --- | --- | --- |
| 0 | 3 | 4 | 3 | 1 |
| 1 | 2 | 5 | 4 | 1 |
| 2 | 1 | 6 | 3 | 3 |
| 3 | 0 | 4 | 1 | 3 |

For the BW filtration at a grey intensity level, *f* = 0 (on a scale from 0 to 4), the cubical complex consists of three distinct parts marked with black dots. This results in the Betti numbers being *β*_0_ = 3 for the three components and *β*_1_ = 0 indicating the lack of holes. As the filter transparency increases to 1 (*f* = 1), one more disconnected component is added, increasing the zeroth Betti number *β*_0_ to 4. In the next filtration step *f* = 2, there are 2 changes: (1) two components are joined (bottom right corner), decreasing *β*_0_ by 1, and (2) another component merges with new elements joining the filtration, forming a hole, thus increasing *β*_1_ to 1. At intensity *f* = 3, all 3 shapes merge into a single connected component with one hole (*β*_0_ = 1 and *β*_1_ = 1). If we further increase filter intensity to *f* = 4, all pixels will be added to the cubical complex, yielding Betti numbers *β*_0_ = 1 and *β*_1_ = 0.

Now, for every filtration step *f* with non-zero *β*_1_, we can find the dual step, *f ^′^* = *M −* 1 *− f*, in the reverse filtration, where there exists dimension 0 component, which we call its dual. For example, as shown in Fig. 4 for *f* = *{*2, 3*}* top row, a cycle exists in the top left corner, and in WB filtration for *f ^′^* = *{*1, 0*}*, bottom row, there is a connected component present in that corner). If we study the reverse filtration, the same rule applies- for the two steps, *f ^′^* = *{*2, 3*}* in which *β*_1_ is not zero, we can find dual components in steps *f* = *{*1, 0*}*. In all cases, the persistence of dual components is equal (unless there is an interaction with a cycle on the periphery). This can also be seen within the barcodes and persistent landscape, as demonstrated with another example where most cycles do not interact with the periphery of the image in Supplementary Fig. Supp.B.6.

Altogether, with this duality, we have shown that the topological properties derived from a single filtration in both dimensions are similar to those from both filtrations but in the same dimension, and we exploit this fact in constructing feature maps.

#### 2.5.3 Topological features maps

In visual art, artistic impressions are created with objects, colours, and textures. Whereas the first two are directly impacted throughout the artistic process, texture is controlled at the earliest stages, when the technique is chosen. Furthermore, texture has a dual nature- for images, the texture is the spatial distribution of colours or changes in contrast on the canvas, whereas, for physical objects, texture refers to the smoothness or roughness of an object’s surface. In painting, texture is a combination of these two aspects, as both are combined (by the artist’s selection of canvas and paint) to create the overall artistic impression.

In this space of objects, colours, and textures, our method is inherently sensitive not only to shapes of various sizes but also to their spatial distribution, which makes it suitable for investigating both objects and textures. The motivation is that, in areas with rich texture, there are many small patches of colour, which translate into a large number of cycles per unit area. On the other hand, smooth areas do not vary as abruptly in terms of colour, so they shall contain fewer distinct cycles per unit area (low cycle density).

All values from a feature map yield a distribution that was specified using its Empirical Cumulative Distribution Function (ECDF). This function is the cumulative sum of the count of windows having a given property value (e.g. a cumulative sum of a histogram showing the counts of windows having a given cycle density). Even though some 2D spatial information is lost, we shall show that the ECDF provides a convenient way of comparing and visualising distributions derived from the feature maps of the images combined with the gaze maps of individuals as will be seen below.

To explore textures and differences in the spatial extent of visual structures between the two sets of images and their effect on perception, we introduce topological feature maps. The pipeline visualising this process is shown in Fig. 5. Topological feature maps are topographic maps conveying the spatial location of properties of various topological features of an image that were obtained by persistent homology calculations. Starting with the image depicting the location of all 1-dimensional cycle representatives (e.g. Fig. 13), one can overlay a square grid mesh consisting of non-overlapping windows of fixed size. Then, for each window, a feature is derived-it could be either cycle density (computed as the total number of cycles in each grid window), maximal 1-dimensional cycle perimeter (the maximal number of pixels in the perimeter of a cycle lying within grid window) or maximal persistence (given by the largest persistence of all the cycles that are found in the grid window). A feature map can therefore be specified by a matrix *M* where, depending on the choice of topological descriptor, *M_i,j_* might be the number of cycles in the square in the *i*th row and *j* the column of the grid (in the case of the cycle density map). Full details can be found in the Supplementary Information sections A.3, A.4, and A.5.

**Figure 5:**
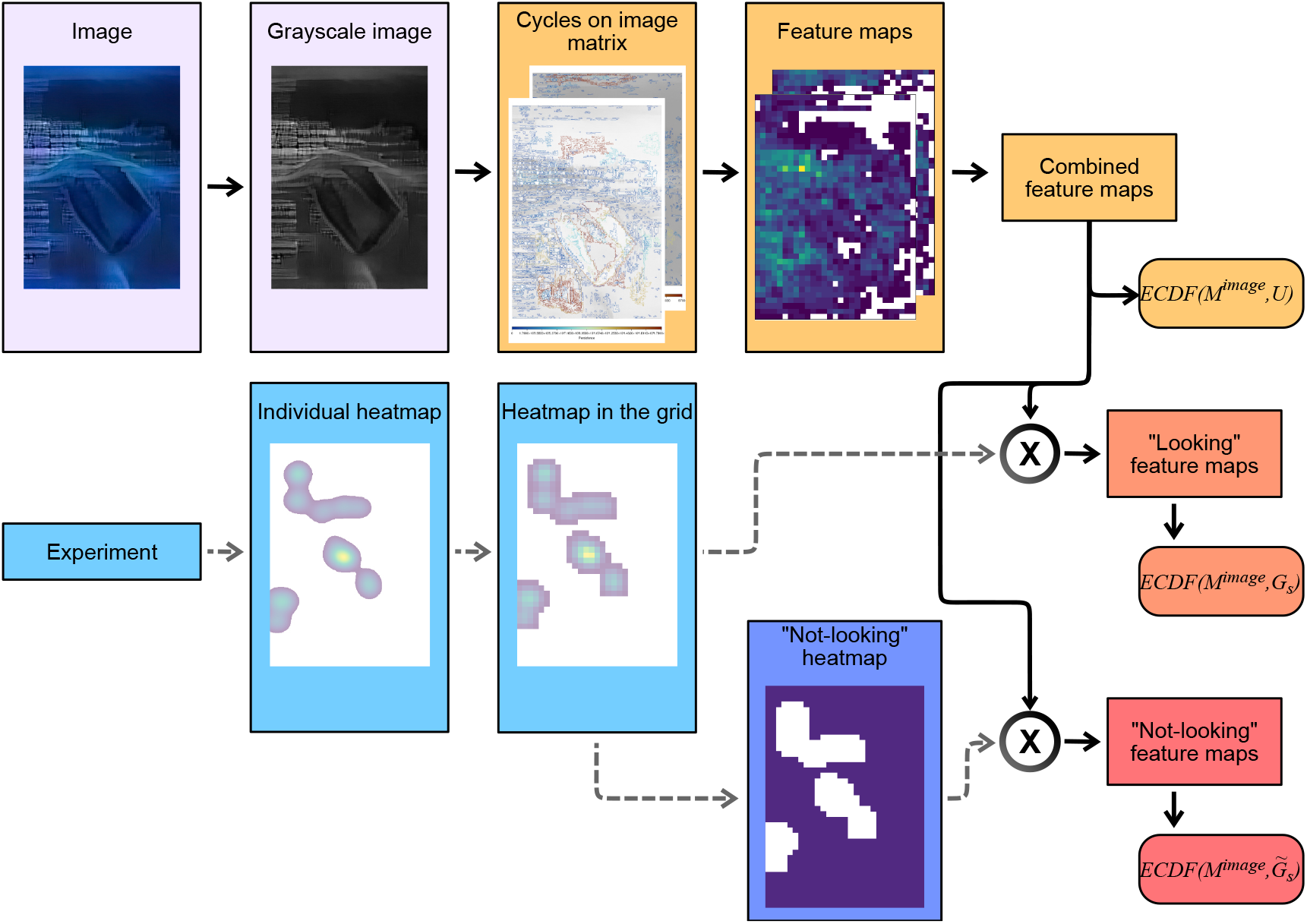
Visualisation of the pipeline used for generating 3 types of ECDF curves. After the image is converted to greyscale, the spatial distribution of cycles is generated from the BW and WB filtrations. From this, 3 types of feature maps are derived (for clarity, only one is shown in the image), and an *ECDF* (*M ^image^, U*) (intrinsic to the image) is obtained. Using the gaze-heatmaps from a subject *s*, *Gs* its complement heatmap *Gs*, *ECDF* (*M ^image^, Gs*) (looking) and *ECDF* (*M ^image^, Gs*) (not looking) are obtained by weighting the image feature maps with the corresponding gaze-heatmap.

These three topologically derived measures, cycle density, maximal persistence, and cycle perimeter, are proxies for texture in the image, local contrast, and shape size, respectively.

As mentioned, feature maps are derived from the locations of 1-dimensional cycle representatives, and this, at first glance, excludes the information about the topology in dimension 0. To overcome this, we exploit the duality feature of cycles and incorporate the information about the cycles in dimension 0 from BW filtration by studying their dual in dimension 1 in WB filtration. As explained in Supplementary Information Section B.3 and shown in Supplementary Fig. Supp.B.7 very little is lost by only using dimension 1 cycles.

Finally, we devised composite feature maps derived from both filtrations (WB and BW) and used these composite feature maps in our study. The feature maps were as follows: (1) cycle density, which was obtained from summing the densities per unit grid area from both filtrations; (2) maximum cycle persistence, which took the maximum persistence of all the cycles from both filtrations present per unit grid area; (3) maximum cycle perimeter, which took the maximum perimeter of all the cycles from both filtrations present in any particular square in the grid.

All images were the same size, and the grid mesh used, 50 pixels by 50 pixels, was the same for every image. Although grid size may seem arbitrary, we verified that the choice makes very little difference by testing multiple versions of the grid size. In each case, a clear separation was achieved (see Supplementary Fig. Supp.B.13 for the demonstration; more results are presented in Supplementary Materials B.6). Cycle representatives for dimension 1 homology classes used in this study were obtained using the ”Ripserer.jl” software [Bauer, 2021,Kaji et al., 2020,C^̌^ufar and Virk, 2021, C^̌^ufar, 2020] and they are shown in the Supplementary Section B.5 for art and pseudo-art and both filtrations; a relation of ECDF and cycles density is shown in the Fig. 14.

Further details, together with the algorithmic framework for this procedure, are presented in the Supplementary Materials A.4.

Altogether, the study of ECDF curves reveals the nature of the paintings’ texture, but as shown in the next section, gaze-weighted ECDF curves were also used as an indicator of what features the participants were interested in while interacting with art.

Our exploration of topologically derived properties is then grounded in their inherent independence from arbitrarily defined coordinates and metric attributes of perceived objects [Carlsson, 2009]. It allows for direct comparisons between different images by examining their intrinsic features. In the context of 2D visual art, persistent homology primarily considers and analyses images based on either a single greyscale channel or three colour channels. The use of a common scale of 256 points for pixel intensity ensures a fair and unbiased comparison across all analysed images. Topological analysis using pixel-based cubical complexes is relatively fault-tolerant to differences in image resolution. Our analyses show that the output of topologically derived features is only affected after a large reduction in the resolution of analysed images. For details, see Fig. Supp.B.8 in supplementary materials. Furthermore, topological properties exhibit a notable resilience to disturbances, such as variations in visual acuity or noise [Smith et al., 2021]. It is also free from the arbitrary biases that researchers typically introduce.

**Analysis of the relationship between feature maps and fixation patterns** Potential differences in the eye movements of the investigated groups could have resulted from the different topological properties of the observed image sets. To verify this hypothesis, we compared information from eye-tracking movements and topologically derived properties of the images.

To this end, fixation sequences lasting over 75ms were extracted from the eye-tracking data of each participant and used to generate fixation heatmaps. These heatmaps, created by applying a Gaussian kernel to qualifying fixation positions, were produced for each participant, image, viewing, and session (a total of 4 viewings, 2 per session). More details on heatmap generation can be found in supplementary materials in A.3.

Subsequently, each of the heatmaps was overlaid onto the feature maps (cycle density, cycle parameter, or maximal persistence) derived for each image, giving a weighted distribution (and subsequently, weighted ECDF) of the parameter.

The idea behind weight, which was the average time a participant looked at the region of the image, was that subjects’ gaze was drawn by the topological features- the more participants spent time looking at some window from the mesh grid, the more important that window was and the higher the weight; this also means that the topological properties of that window are more important.

Two comparisons were made to investigate what kinds of features participants’ eyes were drawn to. First, we compared the topologically derived properties of the areas that subjects looked at in the two datasets. This was done by comparing the distributions of the mean squared ECDF errors (MSE), computed as a mean squared difference of *ECDF* (*M^image^, U*) and *ECDF* (*M^image^, G_s_*) (looking), that is a mean squared difference of the ECDF from feature map of an image and ECDF derived from a feature map of the same image weighted with a heatmap of subject *s* viewing it. The statistical tests were performed by comparing the distribution of MSE for artistic images with the distribution of MSE for pseudo-artistic images. The second comparison was made between areas subjects looked at and areas where they did not look. This was computed as a mean squared difference of *ECDF* (*M^image^, G_s_*) (looking) and the ECDF derived from the complement gaze *ECDF* (*M^image^, G_s_*) (not looking), this way comparing the differences of features that a viewer, s, looked at with the features in the image that the same viewer did not explore.

The comparisons were performed between artistic and pseudo-artistic images. In all comparisons, data was used from all images for all visits and sessions and repeated for three feature maps- maximal cycle persistence, maximal cycle perimeter, and cycle density. More details of this procedure, including a precise definition of these feature maps, can be found in the Supplementary Information A.5.

### 2.6 Statistical analysis

The statistical methods used in eye tracking and EEG analysis are described in the sections on EEG and eye tracking data analysis. The following are the statistical methods used for persistence landscapes comparison and image analysis.

#### 2.6.1 Topological data analyses

**Empirical Cumulative Distribution Function analysis** In the study of the spatial distribution of cycle density, the shapes of the ECDF curves generated from the grid of images were compared using the Kolmogorov-Smirnov (KS) statistic [Nikulin, 2020, Smirnov, 1944]. The choice of the grid size was motivated by achieved total number of windows (smaller grid size)- the larger it was the better the statistical power of the KS test.

Friedman test was used to assess the difference between regions where subjects looked at and did not look at (using combined data from two exhibitions, one test for each of the feature maps), and in the post hoc analysis, the Wilcoxon signed-rank test was used to determine differences within blocks (with one block being data for the Artist exhibition and another for the exhibition with pseudo-artistic images).

#### 2.6.2 Statistical Image Properties (SIP)

Many statistical measures can be applied to the analysis of images such as artworks. As a benchmark for the persistence homology method, we have chosen statistics commonly used for artistic image analyses representing a wide variety of parameters, from spectral to non-linear measures such as complexity or entropy. The final set of SIPs used for comparison with persistent homology analyses included: Fourier slope - the least squares fit of a line to the log-log power spectrum of the images transformed into the frequency domain [Koch et al., 2010], Self-similarity

- self-similarity assessment based on the Pyramid Histogram of Oriented Gradients (PHOG) that is of occurrences of gradient orientation computed on a dense grid of uniformly spaced cells on an image to facilitate edge detection [Amirshahi et al., 2012], Complexity - the mean norm of the Histogram of Oriented Gradients (HOG) across all orientations [Redies et al., 2012], Anisotropy
- the variance of gradient strength in the HOGs across its bin entries [Redies et al., 2012] and the edge density [Redies et al., 2017]. The details of the statistical methods can be found in the cited articles. The analyses were performed using scripts provided by the authors of relevant methods provided in the supplementary information to the cited articles.

The p-value represents the probability of obtaining the results by chance alone. In this study, a value below 0.05 was used to indicate the statistical significance of the results.

## 3 Results

### 3.1 Results of the eye tracking and EEG data analysis - differences between groups watching the two image sets

The results showed statistically significant differences between the groups in terms of eye movements under both laboratory conditions and during gallery visits, as well as in the EEG analyses in the laboratory.

#### 3.1.1 Results of the eye tracking data analysis

Analyses of data collected during gallery visits showed significant differences between groups and visits (Kruskal-Wallis ANOVA statistics=76.1425, *p*=2.06e-16). A subsequent post hoc Dunn test showed significantly longer average visual intake duration in the group viewing pseudo-artistic images for the first (*p*=1.54e-08; Fig. 6-A) and second (*p*=2.69e-09; Fig. 6-B) visit.

**Figure 6:**
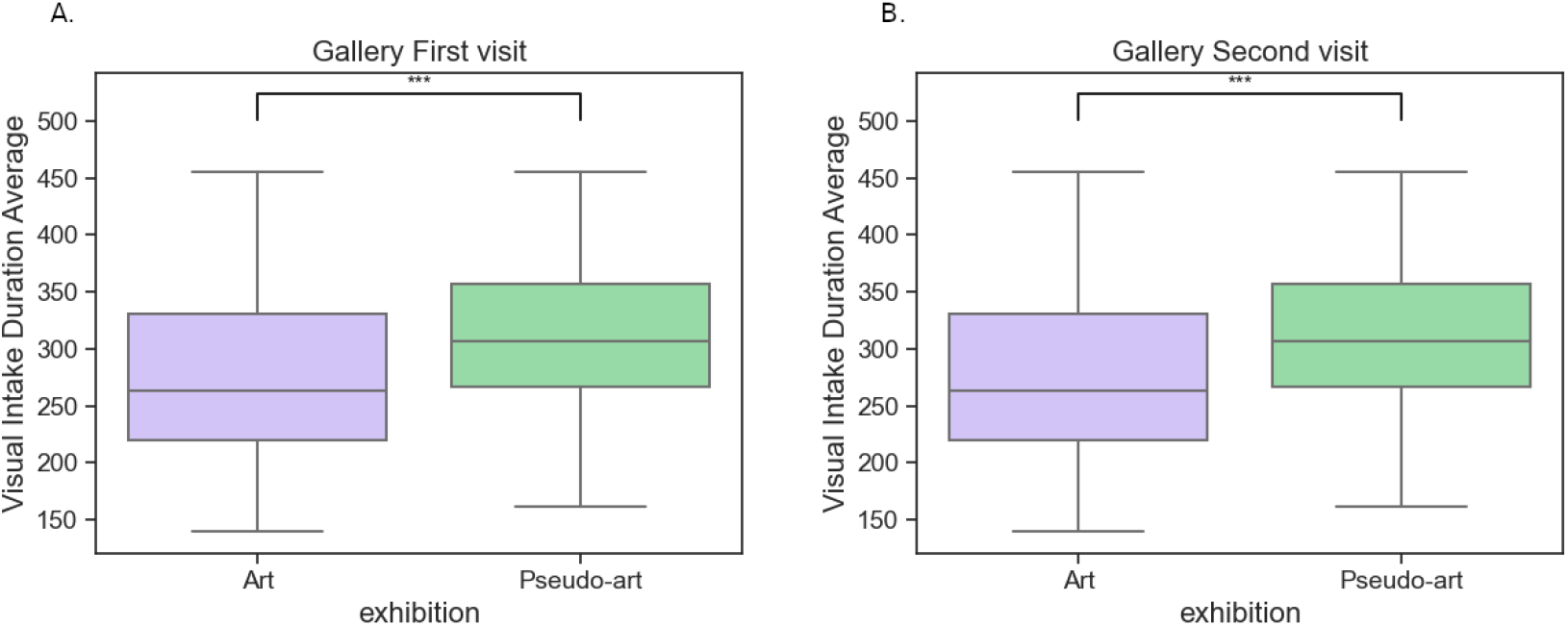
Comparison between art and pseudo-art of average visual intake duration during first (A) and second (B) visit in a gallery. The boxplots show medians for a given group (horizontal line) together with the middle 50% of the data denoted by interquartile range box (the distance between the first and third quartiles) and ranges for the bottom 25% and the top 25% of the data values, excluding outliers (whiskers). *** above horizontal line over 1st visit boxplot denotes *p <* 0.001

Similar analyses of the average saccade durations also yielded significance of the non-parametric ANOVA test (Kruskal-Wallis statistic=165.1596, p=1.41e-35), and the post-hoc Dunn test showed longer average saccades in the case of artistic images for the first (*p*=8.41e-19; Fig. 7-A) and second visit (*p*=2.39e-18; Fig. 7-B).

**Figure 7:**
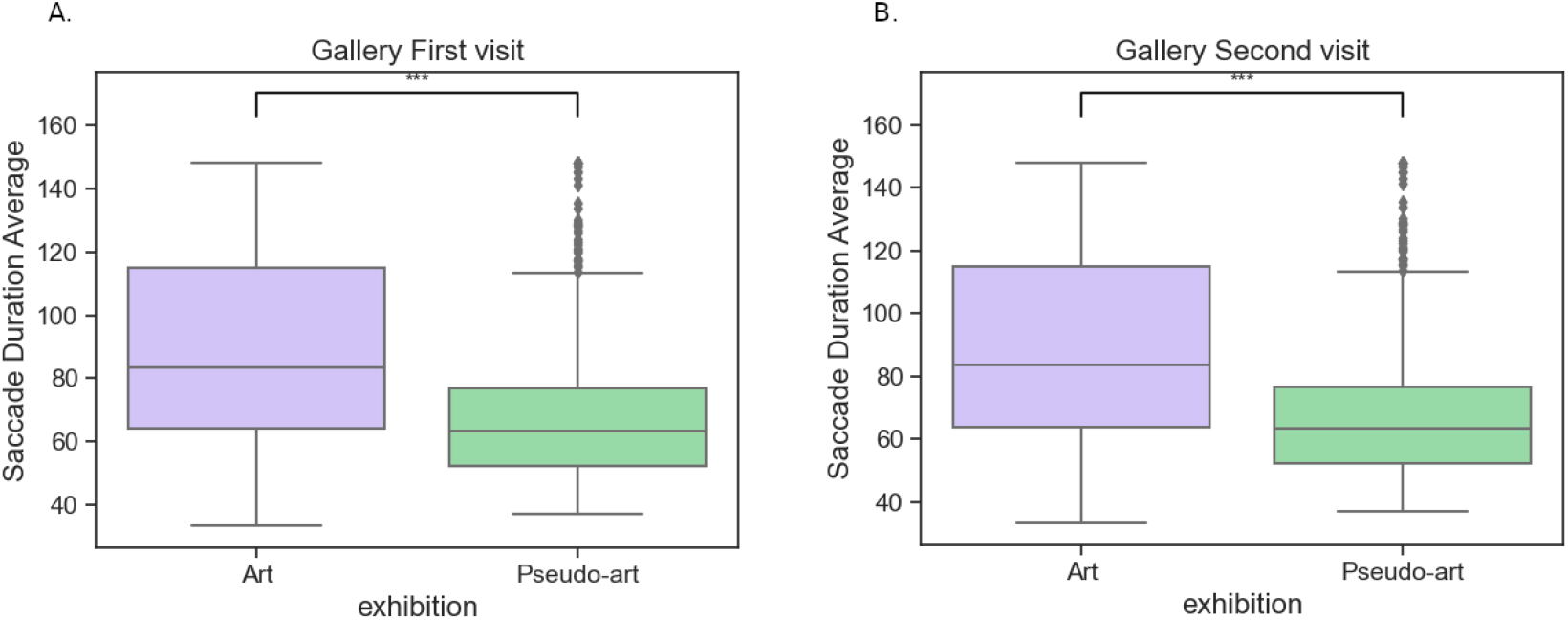
Comparison between art and pseudo-art of average saccade duration during first (A) and second (B) visit in a gallery. The boxplots show medians for a given group (horizontal line) together with the middle 50% of the data denoted by interquartile range box (the distance between the first and third quartiles) and ranges for the bottom 25% and the top 25% of the data values, excluding outliers (whiskers). *** above horizontal line over 1st visit boxplot denotes *p <* 0.001

Interestingly, during laboratory visits, the longer eye fixations were observed for the group viewing the artist works. In detail: Kruskal-Wallis ANOVA analyses for the two laboratory visits (week 1 and week 2) and two groups of participants (viewing either the artistic or pseudo-artistic images) yielded the value of the test statistics = 15.8951 and corresponding *p* = 0.0011. The Dunn test subsequently confirmed the significance of the differences in the first (*p* = 0.043; 8-A)) and second (*p* = 0.0049; 8-B)) visit.

Comparisons of the average saccade duration between the artistic and pseudo-artistic images showed no significant differences (Kruskal-Wallis statistic=6.8328, *p* = 0.0774)

#### 3.1.2 Results of the EEG data analysis

The reception of abstract art is considered a serious intellectual challenge leading to interactions between neural systems shaping sensory-motor responses, evaluation and meaning-making [Chatterjee and Vartanian, 2014, Chatterjee, 2014, Shimamura and Palmer, 2012]. Assuming that the two sets of images may present different challenges to the participants, one may expect different organisation of brain networks involved in the analyses of presented images. To investigate this possibility, we compared the strength of the EEG signal correlations between the two groups.

The initial comparison of the EEG connectivity based on the resting-state signals collected prior to gallery and laboratory visits showed no significant group differences in neither investigated band (Maximum U statistic values and minimum p values for all pairs of electrodes in analysed EEG bands: delta *U* = 810.5*, p* = 1.0, alpha *U* = 789.5*, p* = 1.0, beta *U* = 808.5*, p* = 1.0, gamma *U* = 888.5*, p* = 0.0807 FDR corrected), thus excluding the possibility that possible disparities observed during image viewing stemmed from initial group variations.

The comparison of the group averaged signal correlations while participants were viewing images yielded significant differences in three out of five analysed EEG bands. In the delta and gamma bands connectivity strength was higher for the participants watching artist images, while in the beta band correlations were stronger in the pseudo-artistic group. Fig. 9. No significant differences were found in the theta and alpha bands.

**Figure 8:**
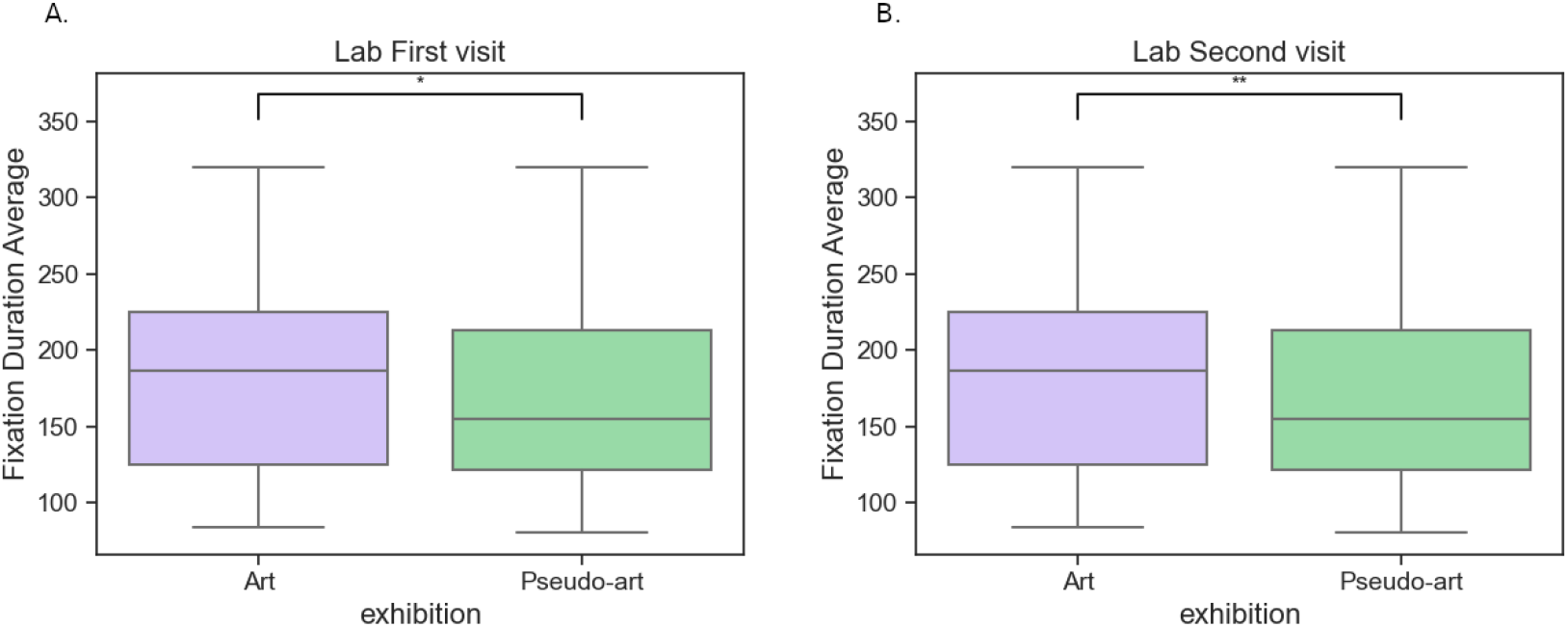
Comparison between art and pseudo-art of average fixation duration during first (A) and second (B) visit in a laboratory. The boxplots show medians for a given group (horizontal line) together with the middle 50% of the data denoted by interquartile range box (the distance between the first and third quartiles) and ranges for the bottom 25% and the top 25% of the data values, excluding outliers (whiskers). * above horizontal line over 1st visit boxplot denotes *p <* 0.05, ** denotes *p <* 0.01

**Figure 9:**
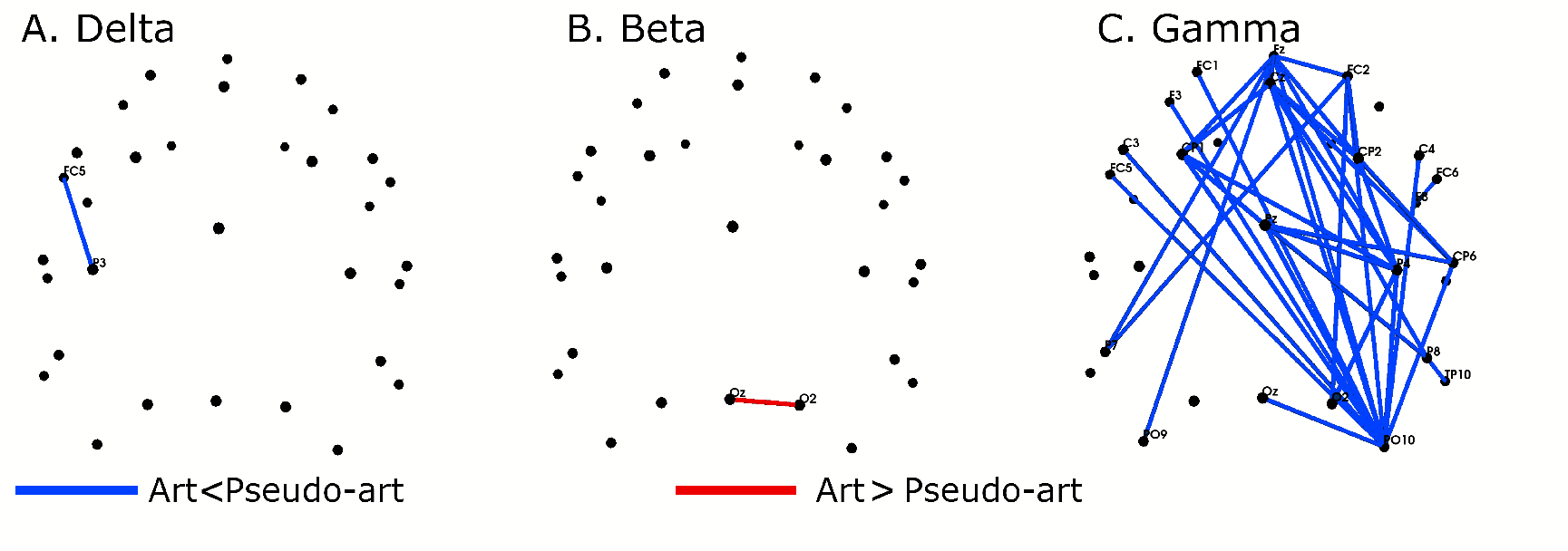
Differences in the strength of EEG connectivity (wPLI) between the artistic and pseudo-artistic groups in delta (A), beta (B) and gamma (C) EEG bands. There were no significant differences in the theta and alpha bands (not shown). Red colour denotes higher connectivity strength in the artistic group, blue in the pseudo-artistic group. All differences are significant at *p <* 0.05 (FDR corrected). Only the names of significant electrode pairs are shown.

Thus, viewing artistic or pseudo-artistic images by two different groups of participants revealed statistically significant differences in eye movements and EEG network organisation. Since our participants were unaware that one set of images was actually generated by the machine learning algorithms, it can be assumed that the differences were due to different processing of the stimuli from the two exhibitions. To investigate whether this might be due to their different topological properties, we examined the persistence homology of images. Then, we compared the spatial distribution of the topological properties of the images with the fixation heat maps.

### 3.2 Topological analysis

#### 3.2.1 Topological analyses of images

Visual inspection of the landscapes of the artist’s images showed that the barcodes spanned almost the entire range of filtration parameters, indicating the presence of some topological cycles with high persistence (a sample is shown in Fig. 10-A, please see Supp.B.3 for all images). For pseudo-artistic images, however, the barcodes were shorter and born in a narrow range, indicating lower persistence as compared to Artist ones, as shown in an example in Fig. 10-B (the visualisation of properties for all pseudo-artistic images is shown in Supplementary Fig. Supp.B.4). Also, the number of cycles for pseudo-artistic images was lower by an order of magnitude than for Artist images in both dimensions (the average of cycles were: *mean^Art^* = 29046 *±* 21542, *mean^Art^* = 26775 *±* 14941, *mean^AG^* = 1725 *±* 1501, *mean^AG^* = 2206 *±* 1462, where *d* subscript is used to denote dimension); this is reflected in the landscapes by the number of layers (Fig. Supp.B.6).

**Figure 10:**
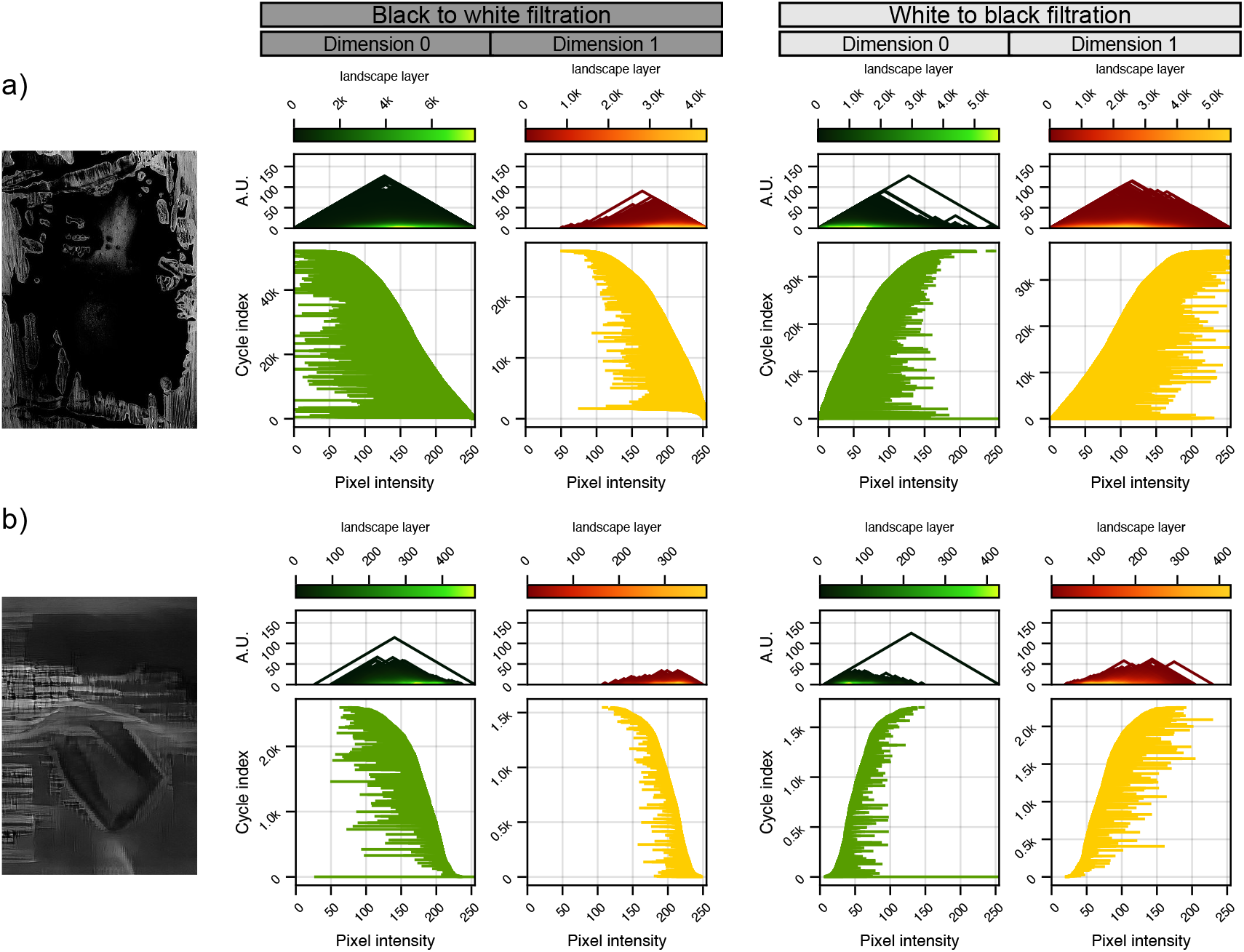
Examples of topological properties of artistic [Kot, 2022](a) and pseudo-artistic images (b). Left column: greyscale-converted original image; middle section: topological results for filtration from black to white; right section: topological results filtration from white to black; within each section-left column: topological characteristic in dimension 0 and right column: topological characteristics in dimension 1. The topological characteristics, the same for both dimensions, are: persistence landscapes(top) and barcodes (bottom). The horizontal axis for both characteristics is pixel intensity filtration steps (in the range [5, 255]). Vertical axes are: for persistence landscapes-arbitrary units; for barcodes - index of a barcode in the birth sorted list of all barcodes for an image. Every landscape (for every dimension) is constructed from a different number of layers- the layers are coloured according to the legend above each landscape.

To statistically assess these apparent differences, we averaged the landscape heights of each layer at each filtration step separately over each set of images. The results are shown in Fig. 11. A visual comparison of the averaged topological properties of the artistic and pseudo-artistic images already shows a stark difference. The statistical non-parametric Wilcoxon test with permutation yielded highly significant differences between the two groups with *p* = 0.0011 and *p* = 0.0014 for dimension 0 and dimension 1 respectively (10000 permutations). Furthermore, as shown in Fig. 12 the Betti curves have dramatically different forms with very little overlap between the two groups for the majority of the filtration steps.

**Figure 11:**
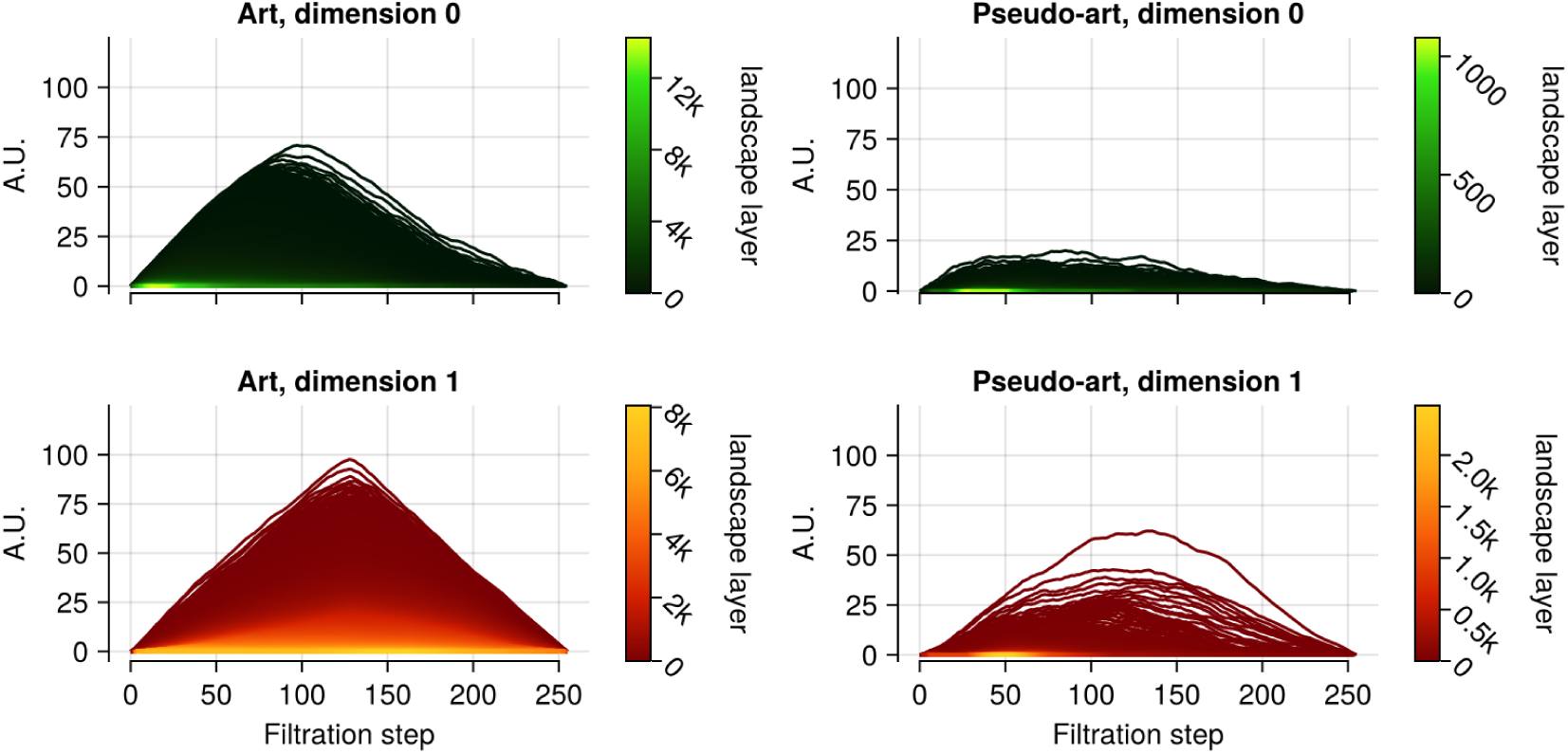
Average group persistence landscapes for BW filtration. The average persistence landscape was computed for each group: artistic (left column) and pseudo-artistic images (right column), with results for dimensions 0 shown in the top row and dimension 1 in the bottom row. Every individual landscape was constructed from cycles of persistence greater than 5 pixel intensity values [Blackwell, 1946]. The average landscapes for artistic and pseudo-artistic images were significantly different for each dimension under the non-parametric Wilcoxon test with permutation, both *p <* 0.001.

**Figure 12:**
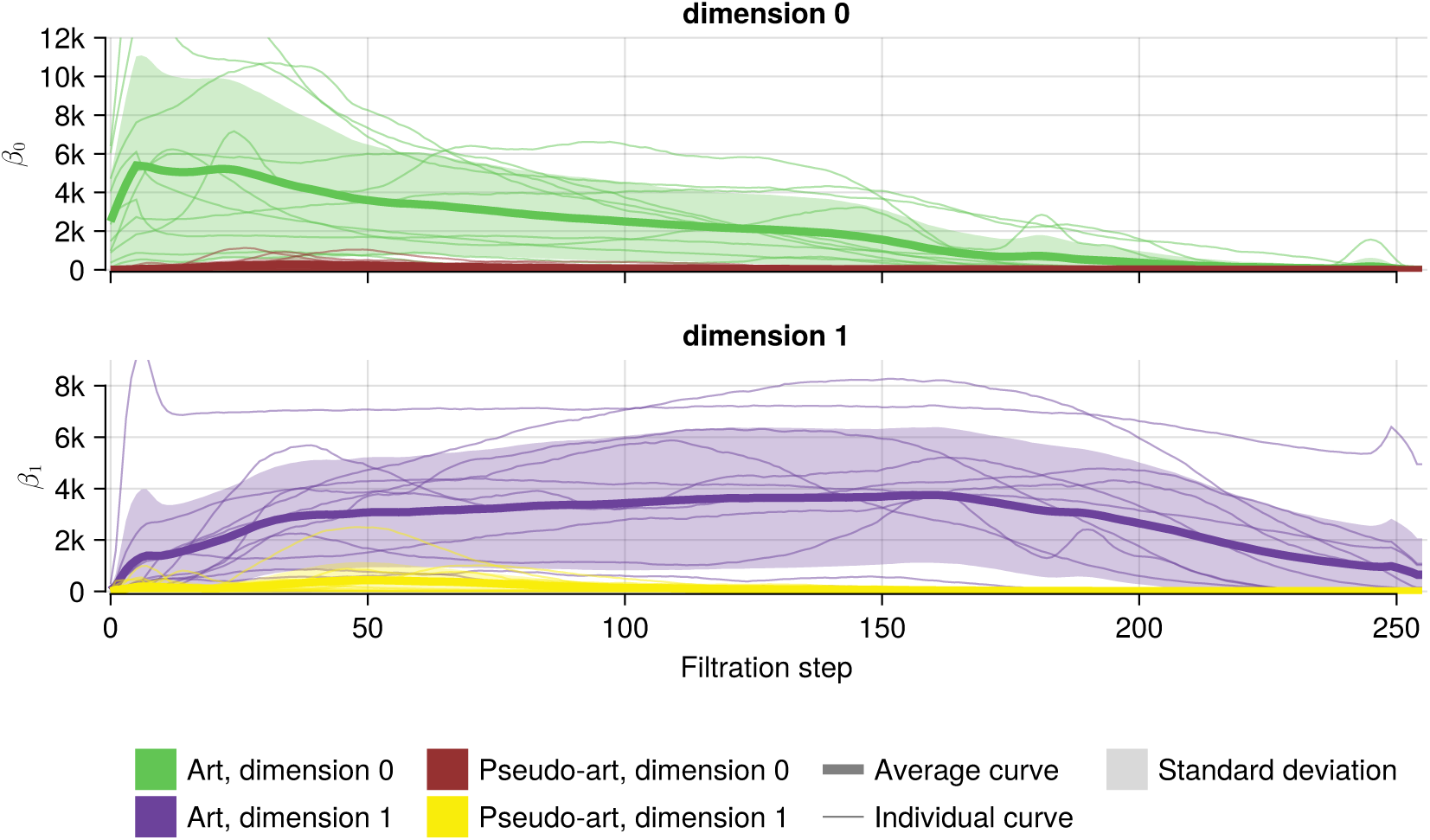
Average Betti curves with 1 standard deviation above and below shaded, together with Betti curves for individual images for the BW filtration. The average Betti curve was computed for each group: artistic and pseudo-artistic images, with results for dimensions 0 shown in the top row and dimension 1 in the bottom row. Supplementary Fig. Supp.B.5 shows Betti curves for the WB filtration.

This allows us to state the following: the study of persistence landscapes from one filtration is sufficient to capture the relation between topological properties across all images, and so far, presented average landscapes from BW filtration are sufficient to describe topological differences between two groups of images. At the same time, it is possible to capture nearly all topological relations in a single image by using only topological structures from dimension 1 and both filtrations. This duality can be seen for all images in Supplementary Fig. Supp.B.3 and Supplementary Fig. Supp.B.4.

#### 3.2.2 Topologically derived image properties and eye movement patterns

The topographies or spatial distributions of the densities, maximal persistence, and cycle perimeter were analysed by computing and comparing empirical cumulative distribution functions (ECDFs). We determined significant distinctions between the density distributions of artistic images and those of pseudo-artistic images.

Fig. 14 shows ECDF curves for cycle density (both filtrations) computed for each of the 24 analyzed images, highlighting the discernible disparities between the two categories resulting in an almost clean separation (more details are presented in Supp.B.14). Changing the window size has little effect on these ECDFs (see Supplementary Fig. Supp.B.13) and certainly does not disrupt the separability of the two sets of images. It exhibits a scale-free property with respect to window size. A clear separation could not be achieved for the ECDFs of maximal persistence (shown in Supplementary Fig. Supp.B.15) or for those of cycle perimeter (shown in Supplementary Fig. Supp.B.16).

So far, we have demonstrated that there were significant differences between the artistic and the pseudo-artistic images: firstly in terms of the persistence barcodes which characterise the stability of the topological features, as evidenced in the radically different average persistence landscapes in Fig. 11. Secondly, there is a lack of intersection among the Betti curves for the two groups as shown in Fig. 12. Thirdly, in terms of topologically-derived topographic descriptors such as the density of cycles as shown in Fig. 14. Next, we sought to find a relationship between the topologically derived properties of the image and eye movements recorded during viewing.

First, we wanted to determine whether participants paid attention to the same topological properties in the artistic and pseudo-artistic images. To this end, we overlaid individual fixation heatmaps onto the topological feature maps generated for each image and computed mean squared errors (MSEs) between the ECDF derived from the same feature map weighted by the fixation heatmaps and ECDFs derived from the image’s feature map (i.e. weighted by a uniform gaze, *ECDF* (*M_k_, U*)). We then compared the resulting MSEs of the ECDF curves (see Methods section for details) of the two groups.

The results shown in Fig. 15 indicate that averaged MSEs calculated for maximal persistence and maximum cycle perimeter were significantly higher for pseudo-artistic images than for artistic ones. In contrast, no significant difference was observed for MSEs corresponding to differences in the cycle density ECDFs (see Table 2 below for the results of the Mann-Whitney U-tests).

**Figure 13:**
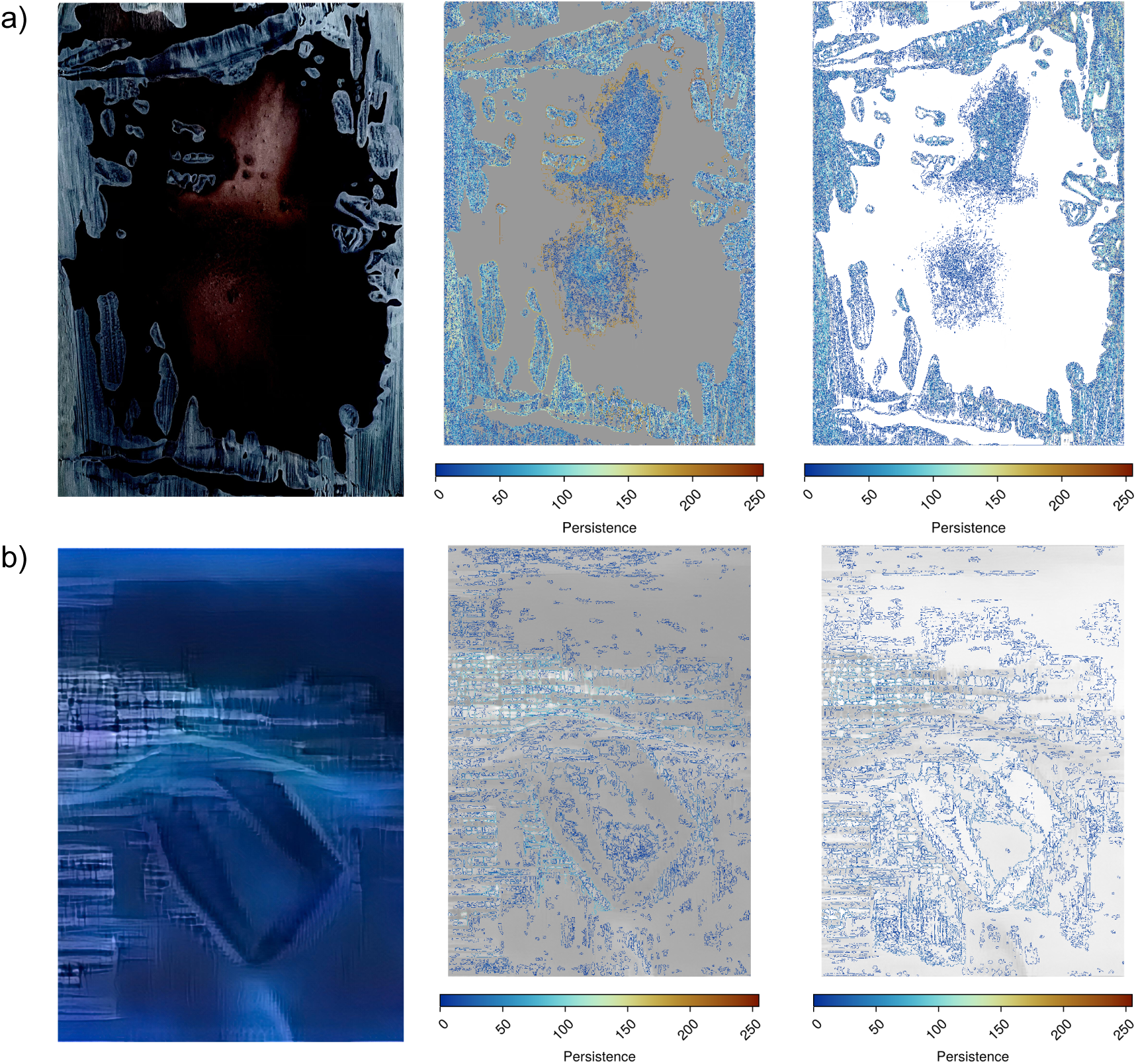
Visualisation of cycles for BW filtration (middle column) and WB filtration (right column) for an artistic [Kot, 2022](a) and pseudo-artistic image (b). Cycles are coloured according to their persistence, as indicated by each legend.

**Figure 14:**
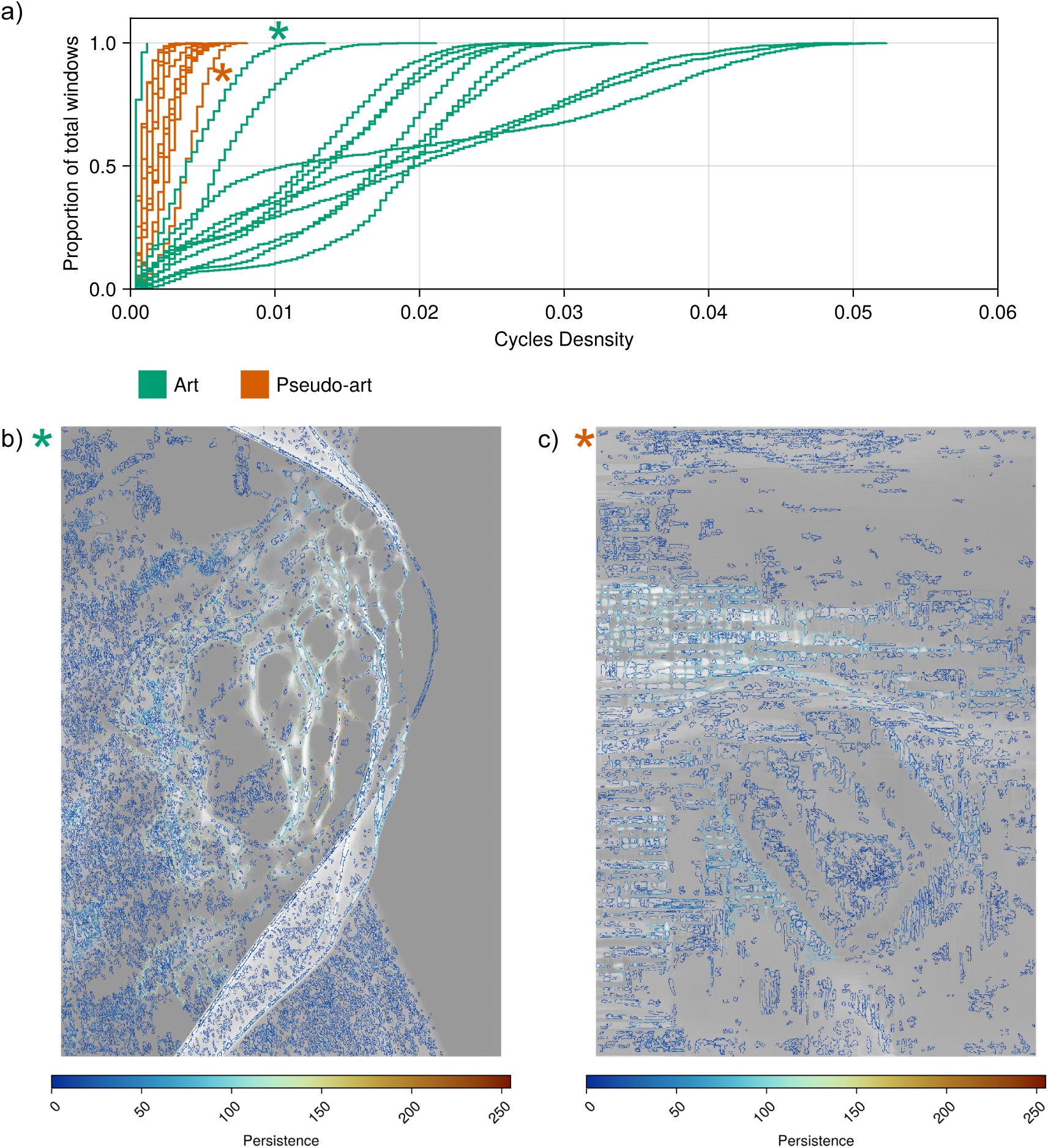
Cycle density comparison for all images (a) and 2 samples of cycles visualisation (b,c). Every curve in (a) represents the ECDF of the cycle density distribution for images for both exhibitions; density used for ECDF is the sum of densities from BW and WB filtrations; artistic (green) and pseudo-artistic (orange). The vertical axis is the proportion of all windows from a grid mesh. All curves were statistically different under the Kolmogorov-Smirnov test. Images for which cycles were visualised are those marked with ”*” in the figure with ECDF plots.

**Figure 15:**
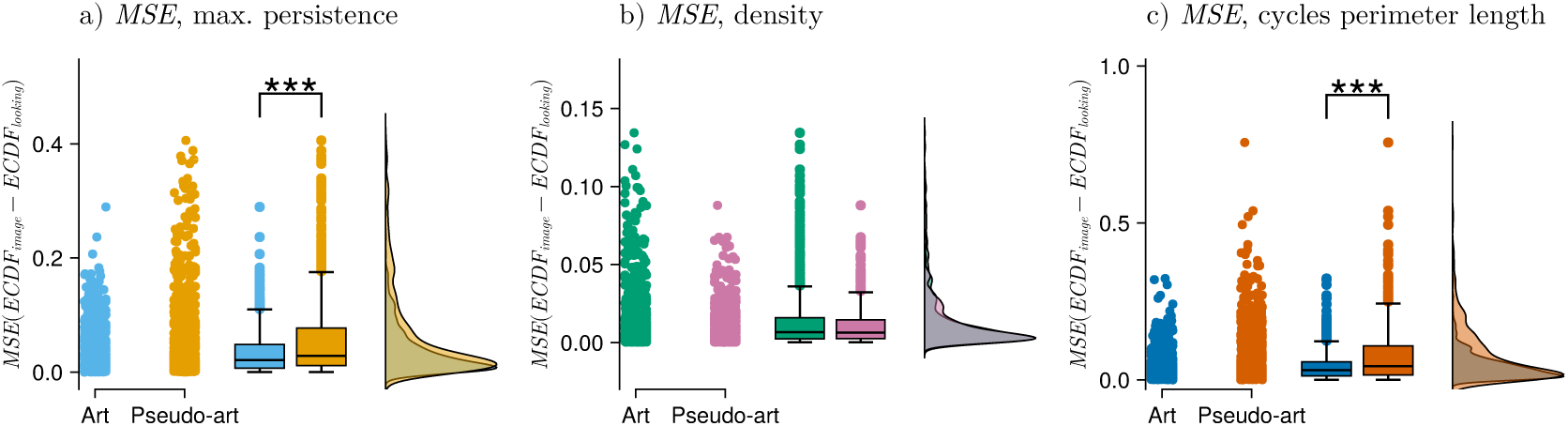
Comparison of how topological features were explored in both exhibitions. MSE between image features distribution ECDF and the features weighted by each participant’s gaze duration data. Results for: a) Persistence; b) Cycle density; c) Cycle perimeter. (Exhibitions are indicated on the axis). The lineup of plots within each subfigure is as in Fig. 8. Statistics performed by Mann-Whitney U test, comparing MSE distribution for the artistic images to MSE distribution for the pseudo-artistic images, *** *p <* 0.001.

**Table 2:**
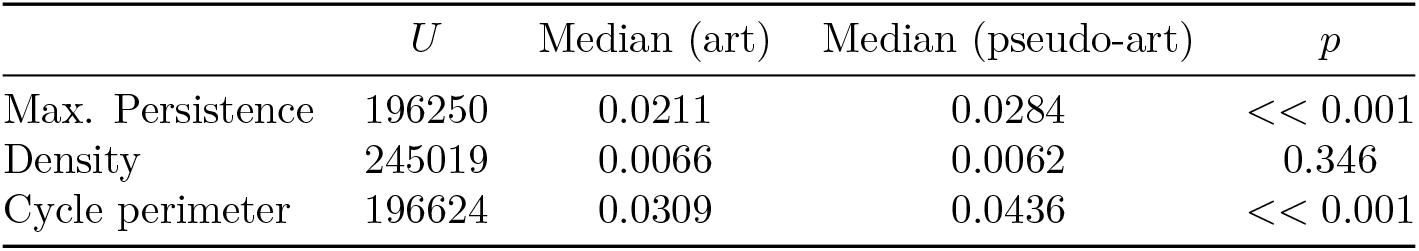
Results of Mann-Whitney U-test. In all cases from Fig. 15 *n_Art_* = 693 and *n_A.G._* = 687.

Next, we sought to identify whether participants are drawn to specific topologically derived properties within a viewed image. To this end, we compared the distributions of persistence, cycle density, and cycle perimeter of the image area covered by each person’s gaze heatmap (‘looking’ *ECDF* (*M_k_, G_s_*) for person *s* looking at image *k*) with the distributions for the same derived properties but of the uniformly weighted areas outside the person’s gaze heatmap (‘not looking’ *ECDF* (*M_k_, G_s_*)). The MSE between these two ECDFs was computed and plotted for each viewing of each image by each participant and plotted in Fig. 16.

**Figure 16:**
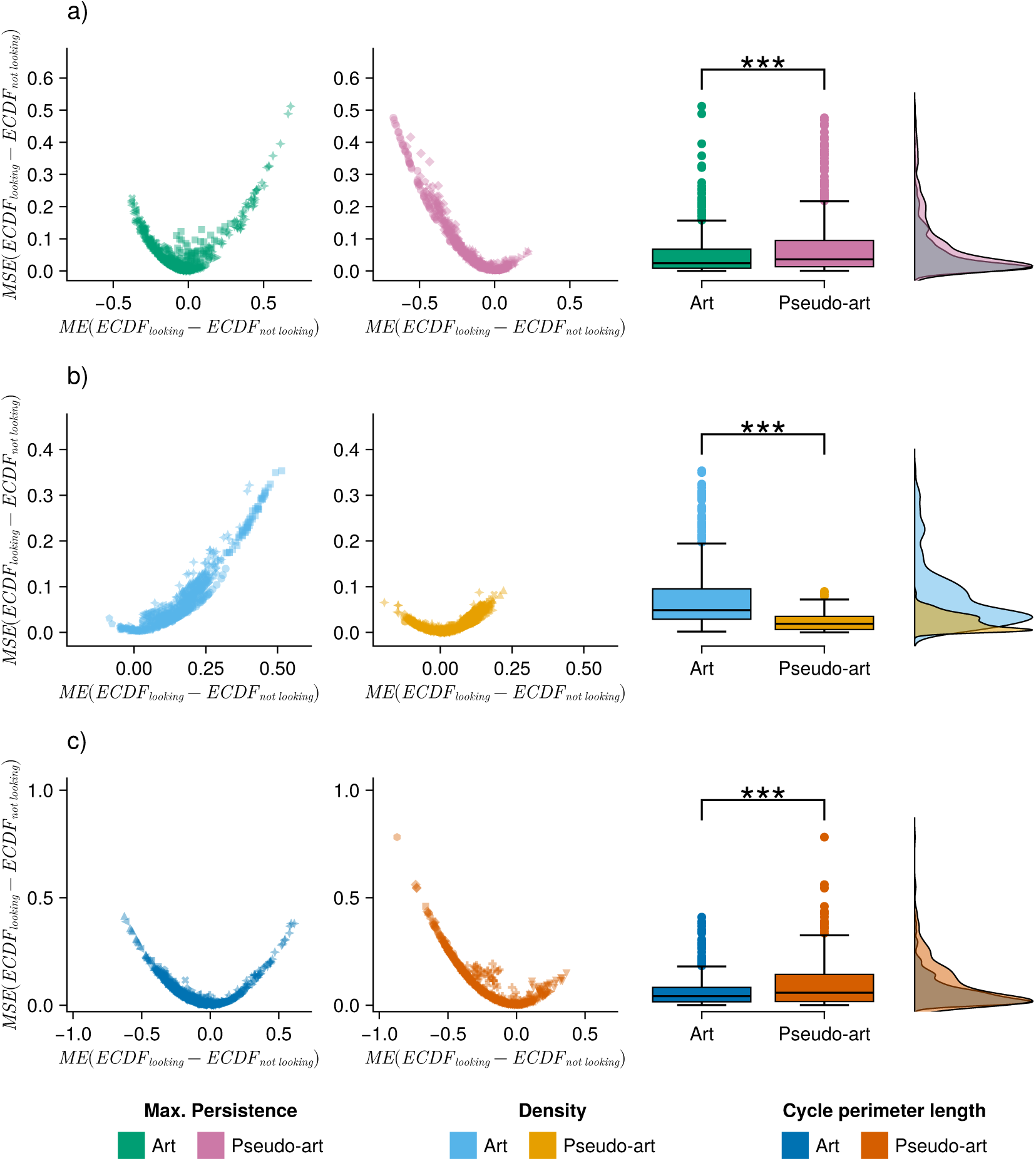
Analysis of topologically derived features that participants were attracted to. Each point on the plots is derived from the ECDFs obtained from a single participant viewing a single image. For every image and every participant who viewed the image, a weighted feature map distribution was obtained from: (1) the regions where the participant looked (weighted by gaze duration) and (2) the regions where the participant did not look (uniform weighting). a) results for persistence feature maps; b) results for cycle density feature maps; c) results for cycle perimeter derived maps (exhibitions are indicated in the legends). In each plot in the first two columns, we show MSE on the vertical axis and ME on the horizontal axis, both computed from the difference *ECDF*_looking_ *− ECDF*_not looking_ for each participant, i.e. it compares the ECDF of regions that a participant looked at with the regions that the same participant did not look at. The third column shows the distribution of MSEs on which significance tests were conducted. Statistics performed with Mann–Whitney U test, comparing MSE distribution of ECDF between the artistic and pseudo-artistic images, *** *p <* 0.001.

The results indicated that pseudo-artistic images had significantly higher MSE for maximal persistence than the artistic images (Mann-Whitney U test, two-sided *p <<* 0.001; Fig. 16). At the same time, a significant difference was observed for MSE of the cycle density (Mann-Whitney U test, two-sided *p <<* 0.001) and for cycle perimeter ECDF (Mann-Whitney U test, two-sided *p <<* 0.001). All details about the tests are presented in Table 3.

**Table 3:** Results of Mann-Whitney U-test for MSE of ECDF for looking and not looking. In all cases from Fig. 16 *n_Art_* = 693 and *n_A.G._* = 687.

| | $U$ | Median (art) | Median (pseudo-art) | $p$ |
| --- | --- | --- | --- | --- |
| Max. Persistence | 200488 | 0.0240 | 0.0362 | $<< 0.001$ |
| Density | 377815 | 0.0486 | 0.0189 | $<< 0.001$ |
| Cycle perimeter | 204175 | 0.0423 | 0.0578 | $<< 0.001$ |

Summarizing the analyses of the topological properties being of special interest for artistically oriented participants, we found that both groups preferentially fixated on the image areas characterized by higher persistence as shown by Fig. Supp.B.17. Here, the blue ECDF for each image, *ECDF* (*M_k_, U*), is intrinsic to the image itself and can be thought of as arising from a ‘gaze’ that is a uniform scan of the entire image.

In the case of cycle density, the preference towards areas of higher density was found only in the group viewing the pseudo-artistic artificially generated images.

### 3.3 Comparison of the persistence homology with current state-of-the-art methods

In order to compare image set discrimination using topological properties and commonly used statistical image properties, we ran a series of comparisons between the two sets of images using total landscape area in dimension 0, total landscape area in dimension 1, Fourier slope, Self-Similarity, Complexity, Anisotropy and edge-density using non-parametric Wilcoxon test with FDR adjustment for multiple comparisons. The results are shown in Table 5.

**Table 4:** Summary of image property preferences from Fig. 16.

| Property | Art | Pseudo-art |
| --- | --- | --- |
| Max. Persistence | No clear preference | **Preference for more persistent cycles** |
| Cycle Density | **Preference for lower density** | No clear preference |
| Max. Cycle Perimeter | No clear preference | **Preference for cycles with greater spatial extent** |

**Table 5:** Comparison of averaged topological and statistical image properties. Results of the non-parametric Wilcoxon test. Significant differences are shown in bold.

|  | Topological image properties |  | Statistical image properties |  |  |  |  |
| --- | --- | --- | --- | --- | --- | --- | --- |
|  | dim0 | dim1 | Fourier slope | Self-similarity | Complexity | Anisotropy | Edge density |
| uncorrected | 0.0011 | 0.0014 | 0.0464 | 0.3123 | 0.0011 | 0.1749 | 0.0061 |
| FDR corrected | **0.0033** | **0.0033** | 0.0650 | 0.3123 | **0.0033** | 0.2041 | **0.0107** |

Although a statistical comparison of the groups showed significant differences for four different measures, including two topological ones, it would be interesting to see how well the different measures perform in separating the groups. To this end, we plotted for each of the measures data points and boxplots for the groups being compared. Results of this comparison are shown in Fig. 17.

**Figure 17:**
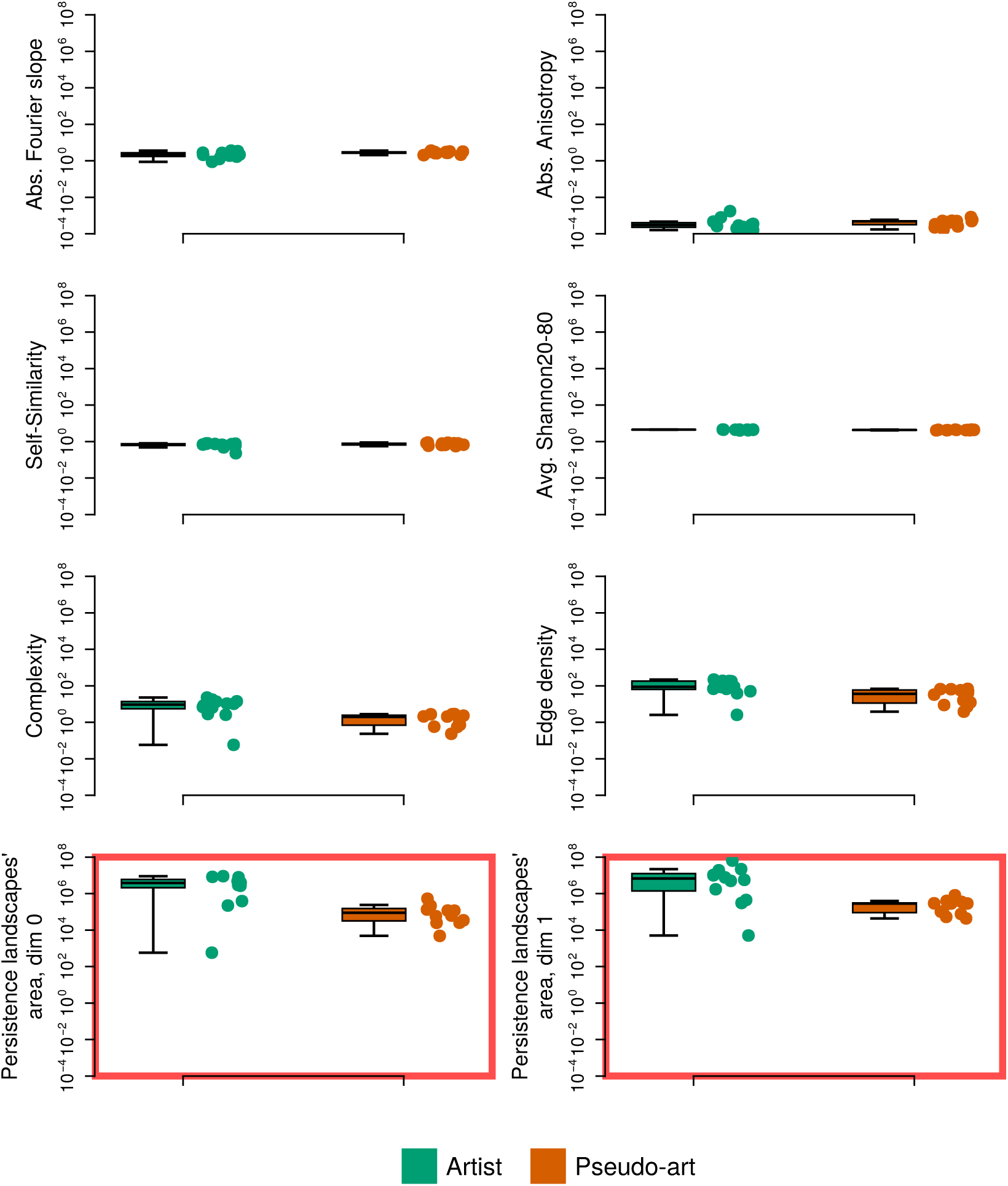
Comparison of topological properties (outlined by red rectangle) and chosen statistical image properties; left panel, from top: Fourier slope, self-similarity, complexity, anisotropy; right panel from the top: average Shannon entropy edge density, the area under landscape for dimension 0, the area under landscape for dimension 1 (area under persistence landscapes computed for BW filtration). For each measure boxplot (boxplot explanation the same as in Fig. 4) and corresponding data cloud are shown. Results for artistic images are in green; pseudo-artistic images are in orange.

In conclusion, although some statistical properties of an image, such as complexity or edge density, show significant differences between artistic and pseudo-artistic images, the best separation is achieved only by topologically derived metrics. It is also worth noting that both statistical metrics (i.e. edge density and complexity) are holistic in nature, confirming the relevance of the choice of topology as a method for analysing visual art.

## 4 Discussion

Our results show that a systematic mathematical exploration of shapes using topological descriptors derived from persistent homology is more effective than using a plethora of commonly used statistical measures (SIPs). Most of the SIPs, apart from complexity, were unable to distinguish between the sets of artistic images and those generated by the artificial neural network (pseudo-artistic images), as shown in Fig. 17. The separation provided by the cycle density cumulative distribution curves in Fig. 14, however, is more informative than that provided by complexity, as the latter only ascribes a single scalar value to each image. Hence, although the measurement of ‘complexity’ was able to separate the two sets of images equally well, it provides merely one macroscopic quantity for each image, which does not provide any scope for further detailed investigation for questions such as ‘What aspects of this image did people prefer to view?’ or ‘What aspects of this image attracted human eyes the most’.

By analogy to thermodynamics, where macroscopic properties such as temperature, pressure, etc., emerge via statistical mechanics from microscopic constituents such as atoms, molecules, and electrons, the SIPs can be considered as macroscopic properties of an image, whereas the pixels can be considered its microscopic constituents. There is also a mesoscale level of understanding in fields such as field mechanics that emerges from examining the collective flow of microscopic particles, e.g. boundary layers, vortex sheets. We posit that persistent homology using cubical complexes is an analogously comprehensive breakdown of visual structure that operates on the mesoscale level that is neither too coarse (macroscopic SIPs) nor too detailed (individual pixel values). Although other hand-crafted measures, such as edge density, can also be used locally, enabling the derivation of a topographical map, such measures do not easily reveal mesoscale structures in a systematic way. Another benefit of a PH approach is that the evolution of these shape structures can be followed via the Betti curves. Fig. 12 shows that this evolution is dramatically different for the two classes of images.

To systematically decompose abstract images, we focused on three properties derived from our topological investigation: persistence, the density of cycles, and the perimeter of 1-dimensional cycles. Cycles are representatives of each topological invariant (homology class) identified by the method. Long-lived or ‘persistent’ cycles are objects that are the most salient in terms of the filtration and are, therefore, the easiest to see, i.e. they are bounded by pixels with a large difference in contrast. Cycle density can be considered a coarse proxy for texture (two distinct textures may have the same cycle density but consist of differently shaped cycles). Cycle perimeter is a measure of spatial extent, as in general, cycles with large spatial extent will have a larger perimeter (in general, it will differ according to fractality present in the image). Cycle perimeter, therefore, reveals mesoscale spatial structures that compose an image. As shown in Fig. 14, cycle density was a property that was able to almost cleanly distinguish between the two sets of images.

Furthermore, a mathematical duality inherent in the construction of ‘opposite’ filtrations of cubical complexes captures the characteristics and shapes of an image that also appeals to the eye. The saliency of some high-contrast structures surrounded by connected components will be missed by using a filtration in only one direction. The advantage of using a pair of opposing filtrations in both directions (from small to large pixel values and in the reverse direction) is that the union of the sets of the dimension 1 cycles of both filtrations gives a comprehensive description of salient features as well as providing a uniform way to measure spatial extent (via cycle perimeter). Using a mixture of dimension 0 and dimension 1 cycles would not allow this. We can thereby systematically interrogate structures in an image on multiple scales: both spatially and in terms of intensity.

Concurrent eye-tracking and EEG experiments enabled us to uncover differences in the ways people viewed and processed abstract art vs abstract images generated by an artificial neural network, both in a relative and an absolute sense. To explain these differences, we compared the distribution of a topologically derived property in the places where each participant looked in an image with the distribution of the same topologically derived features in the totality of the underlying image (using ECDFs). The distributions of the various topological descriptors stemming from the underlying image can be considered proxies for image composition in terms of structures on various scales. Fig. 15 shows highly significant differences when considering maximal persistence and cycle perimeter length. Hence, the contrast and spatial extent of structures within an image, revealed by topological analyses, played important roles depending on whether they were part of an artistic image or a pseudo-artistic image. Thus, the way people viewed images was dependent on the spatial distributions of topological properties of the viewed image itself. This notion that the importance of the particular image feature may depend on the whole image’s properties has so far not been investigated in any studies known to us.

Having established differences in the way people viewed art versus pseudo-art, we further interrogated how people looked at different parts of an image and whether the regions of an image a participant neglected to look at were measurably different to the areas he/she preferred to view. The results from Fig. 16 (summarised in Table 4) showed that when viewing artistic images people had no clear preferences when it came to maximum persistence or cycle perimeter, however, there was a strong preference for regions with lower cycle density. In contrast, when viewing the pseudo-artistic images, there was a strong preference for regions containing more persistent cycles, as well as regions containing cycles with large spatial extent. For the pseudo-art, there was no clear preference for regions in terms of cycle density. These findings add credence to our previous observation that the importance of any particular image feature to the viewer may depend on the other properties of the image that is being viewed.

Particularly interesting results of our study are the earlier mentioned differences in eye movement patterns in the gallery and laboratory. The reversed pattern of the visual intake duration/fixation duration for the laboratory and gallery was present during the first and second visits, therefore it could not be attributed to novelty effect. Some clues to solving this puzzle may lie in the topological properties of the two sets of images. The pseudo-artistic images are characterised by the presence of a higher proportion of low persistent cycles, which, when subject to changing lighting and viewpoint conditions in the gallery, may result in an alteration in the complexes emerging from the filtration used for deriving topological image properties, thus changing the number and shapes of dimension 1 and dimension 0 objects (loops and connected components). This emergence and fading of ‘hidden’ topological shapes caused by the motion of the viewer could give an impression of flickering, thus attracting his or her attention, which is expressed by longer durations of the visual intakes. In the case of artistic images, the topological dimensions were characterised by the existence of many more highly persistent cycles (spanning almost the whole range of pixel intensity); the images were, therefore, more robust to changing lighting conditions and spectator motion.

These observations could not have been made in investigations carried out only in laboratory conditions in the situation where a computer screen is used to project the images to be viewed and analysed. The stable lighting conditions created by the screen and the fact that the light is emitted by the screen deprive the viewer of effects introduced by textures. Texture, such as the brushstrokes that characterize seemingly flat works of art, can affect early visual processing (e.g., structural complexity, symmetry and balance), which are important properties of an artwork, and which affect the perceptual and evaluative processes of the artwork [Pelowski et al., 2017]. Although the study of lighting conditions conducted by Pelowski [Pelowski et al., 2019] did not show any significant effect on viewers’ subjective experience, the results of this particular study cannot be a caveat to our inference. This is because first, the authors studied the effect of different but stable light properties that exclude filtering processes, second, the topological properties of the studied artworks are unknown (they may have high persistence), and third,

- there is no eye-tracking data to compare physiological responses nor comparison with the laboratory experiment.

In our opinion, the observation that changing lighting conditions and viewer movement can affect the visual properties of an image is very important. Most of the original artworks are/were created under different and changing lighting conditions and with the presence of constant movement of the artist observing the effect of his or her work. It cannot be ruled out that modern and past masters were aware of the effect of lighting conditions on viewers’ perception and consciously created their paintings to affect viewers differently depending on whether the painting was viewed in the morning or evening, on a sunny day or in the presence of torchlight. Thus, presenting contemporary and ancient visual art in artificial, stable light with a never-changing spectrum can deprive us of the impressions originally designed by the artists.

In summary, persistence homology, together with our new toolkit for mapping topological features onto fixation heatmaps, offers a comprehensive method for studying human response and perception of visual art. This new approach, along with the toolkit we developed, allows for the following:

1. It provides a holistic, unbiased approach that takes into account shape preferences in human perception.
2. It captures the significance of individual image features in relation to the properties of the whole image.
3. It enables us to grasp the influence of viewer movement and lighting conditions on physiological responses to artistic stimuli.
4. It enables experimenters to fine-tune their visual stimuli more carefully in any experiments as well as leads to the development of more interesting derived measures based on this topological decomposition of the visual stimulus. The alignment and arrangement of cycles can lead to new ways of characterising and systematically generating and modulating textures found in images.

This study has the obvious limitation that we have only used artistic images produced by a single artist. Also, each participant viewed only a single exhibition. Further experiments using a larger number of works from a wider range of artists would be needed to confirm whether these findings are peculiar to the particular abstract artist chosen or hold more generally for abstract art. Nevertheless, the experimental design was single-blind in that considerable steps were taken to ensure that both sets of participants believed they were viewing the work of a real human artist. Despite there being an order of magnitude difference in the total number of cycles and in cycle density, at no point was suspicion raised by the participants that the pseudo-art was in any way ‘artificial’ or not produced by a human artist.

Our current results seem to indicate that the proposed method and the developed tools could be beneficial in future research on visual art.

## Acknowledgments

RJ was supported by a Priority Research Area DigiWorld grant under the Strategic Programme Excellence Initiative at the Jagiellonian University (Kraków, Poland).

