## Supplemental for "Art’s Hidden Topology: A window into human perception"

### A Supplementary Information

#### A.1 Persistent Homology and Betti curves

For each image $I$ we build a cubical complex $C_j(I)$ for each step of the filtration $j \in [0, M]$ ($M = 255$) using the procedure described in Section 2.5.2, where the one-dimensional filtration parameter used is pixel intensity. Hence we have the following chain of inclusions (a family of cubical complexes) to which the persistence algorithm [Carlsson and Zomorodian, 2009] is applied:

$$C_0(I) \subset C_1(I) \subset \ldots \subset C_r(I) \subset C_{r+1}(I) \subset \ldots \subset C_M(I). \tag{A.1}$$

The $i$th Betti curve of $C(I) = \{C_0(I), C_1(I), \ldots, C_M(I)\}$ is defined by:

$$\beta_i(j) = \beta_i(C_j(I)) := \operatorname{rank} H_i(C_j(I), k),\ j \in [0, M], \tag{A.2}$$

where $H_i(C_j(I), k)$ is the $i$th homology group of $C_j(I)$ with coefficients in $k$ ($k$ can be any field; the software implementation that we used, took $k = \mathbb{Z}/2\mathbb{Z}$.):

$$H_i(C_j(I), k) := \frac{\ker \partial_i}{\operatorname{im} \partial_{i+1}}. \tag{A.3}$$

The maps $\partial_i$ are the standard boundary maps. The inclusion maps $\iota_r : C_r(I) \hookrightarrow C_{r+1}(I)$ induce maps on the homology $(\iota_r)_i : H_i(C_r(I), k) \to H_i(C_{r+1}(I), k)$. These induced maps allow us to follow cycle representatives of homology classes as they evolve through the filtration, from one complex to the next. In particular, we note when they are 'born' 'birth time' $b$ (the value of the pixel intensity when it first appears) and when they die 'death time' $d$ (the value of the pixel intensity when it ceases to exist).

**The Persistent Homology** (PH) classes of the cubical complex derived from an image can therefore be summarised by a collection of persistence intervals or 'barcodes'.

**Sketch proof of duality** : Let $\tilde{C}(I)$ be the reverse filtration of $C(I)$ where the pixel intensity order is reversed. The cubical complexes for $\tilde{C}(I)$ at step $f$ are complementary to those of $C(I)$ at step $M - 1 - f$ of the filtration. If the grid of the underlying image is embedded onto an $S^2$ (via 1-point sphere compactification) by Alexander duality (or simple observation in this case) For $[\tau] \in H_0(C_j(I), k)$ there is a corresponding complementary class $[\tilde{\tau}] \in H_1(\tilde{C}_{M-1-j}(I), k)$. The birth time of the corresponding PH representative of $\tau$, $b_\tau$ is related to death time of PH representative corresponding to $\tilde{\tau}$, $d_{\tilde{\tau}}$ via $b_\tau = M - 1 - d_{\tilde{\tau}}$ and vice versa. In our situation, where the image does not lie on a sphere, this relationship holds everywhere except on the boundary.

#### A.2 Persistence landscapes.

For the purpose of comparative study, instead of using standard barcode representation, the results from PH were represented as persistence landscapes [Bubenik and Dłotko, 2017]. For each cycle of a given image and dimension, a piecewise linear function $f_{(b,d)} : \mathbb{R} \to [0, \infty)$ was defined as:

$$f_{(b,d)} = \begin{cases} 0 & \text{if } x \notin [b, d] \\ x - b & \text{if } x \in [b, \frac{b+d}{2}] \\ -x + d & \text{if } x \in (\frac{b+d}{2}, d] \end{cases} \tag{A.4}$$

The intuition behind this formula is to build an upward-pointing, right-angled isosceles triangle with the base positioned between birth and death. Having this conversion of a multiset of $n$ persistence barcodes $\{(b_i, d_i)\}_{i=1}^n$ into functions, it is now possible to construct a persistence landscape as a set of functions $\lambda_k : \mathbb{R} \to \mathbb{R}$ such that $\lambda_k(x)$ is the $k$th largest value of $\{f_{(b,d)}(x)\}_{i=1}^n$ and $\lambda_k(x) = 0$ when there is no $k$th largest value (The index $k$ in $\lambda_k(x)$ denotes the layer of the landscape. This process is demonstrated in Fig. 3 (e.g. barcodes for dimension 1 in Fig. 3-A contributed to a persistent landscape with 2 layers, as shown in 3-D).

When an operation on a few landscapes is performed (e.g. averaging for a group or distance computation between a pair of landscapes), the operation is applied to each layer separately. Therefore, the (mean) average landscape is the landscape that results from taking the mean of each layer separately; hence, the average landscape $\hat{\lambda}$ is defined as a collection of functions $\hat{\lambda}_k(x)$ for each of its layers indexed by $k$:

$$\hat{\lambda}_k\left(x\right) = \frac{1}{n}\sum_{i=1}^{n}\lambda_k^i\left(x\right) \tag{A.5}$$

where $n$ is total number of landscapes to be averaged and $\lambda_k^i$ is as defined as the $k$th layer for the $i$th landscape.

The $L^1$ distance between two landscapes is computed as the total sum of the absolute value of the differences between the two landscapes in each layer separately, where the distance within a layer is the difference of heights of every point where either of the layers is defined. If one of the landscapes has more layers than the other, the missing layers are considered to be zero-height layers. The $L^1$ distance between landscapes $f$ and $g$ is thus defined as:

$$||f-g|| = \sum_{k=1}^{max(K,K')}\int||f_k-g_k|| \tag{A.6}$$

where $K$ and $K'$ are the maximum numbers of layers for landscapes $f$ and $g$ respectively. The norm $||f_k-g_k|| = \int|f_k(x)-g_k(x)|dx$ can be computed as the sum of integrals over the landscape intervals.

Those measures were used to compare the topology of the images from the two sets under consideration.

Persistence homology was computed using a filtration step size of 5-pixel intensity units as the human resolution is restricted to 5 points (Petrov, Y. 2005) [Petrov, 2005]. Additional tests of filtration steps: 0, 2, 5, and 10 showed that only threshold 10 decreased the total landscape area (not shown). PH derived from cubical complexes was computed using the DIPHA library and the persistence landscape toolbox [Bubenik and Dłotko, 2017] rewritten in Julia for this purpose.

#### A.3 Gaze heatmaps from eye-tracking data

Given eye movement data from viewing a single image, fixation sequences lasting over 75 ms were extracted and converted into gaze heatmaps using the Python package *Gaze Point Heat Map* [Roeddiger, 2024]. This outputs a heatmap in the form of a 1434 × 2048, matrix. The heatmap thus obtained is then normalised and converted into a probability distribution. We shall specify each gaze distribution by a matrix $G$, where $\sum_{i,j} G_{i,j} = 1$, where $i$ and $j$ index the rows and columns of a chosen grid size. The grid chosen can be coarse or fine. We shall choose a grid size equal to that of feature maps to obtain distributions of topological descriptors, as described below.

#### A.4 Feature map processing

An image overlaid with cycle representatives of dimension 1 persistent homology classes (visualisation of the representative cycle at its birth) as in Supplementary Fig.-B Supp.B.9, can be covered by a square grid. For convenience, we have chosen a grid with uniform fixed-sized non-overlapping windows.

A feature map can then be specified by a matrix $M$ where, $M_{i,j}$ is the local measure of a chosen topological descriptor in the square in the $i$th row and $j$th the column of the grid, e.g. for cycle density, $M_{i,j}$ is the total number of cycles in the $(i,j)$th grid square (see Table 6 for the other examples used in this paper).

Composite feature maps can be constructed by combining feature map matrices from both filtrations, BW and WB, e.g. for maximal persistence we take $M_{i,j} := \max(M^{BW}_{i,j}, M^{BW}_{i,j})$, where $M^{BW}$ and $M^{WB}$ are the maximum persistence feature maps for the BW and WB filtration respectively. Other feature maps used in this study are summarised in Supplementary Table 6.

Given a feature map $M$ of topological descriptors we can define two types of probability distributions supported on the grid with $n_{row}$ rows and $n_{col}$ columns :

$$P(m = M_{i,j}) = U_{i,j} := \frac{1}{g},\ 1 \leq i \leq n_{row},\ 1 \leq j \leq n_{col}, \tag{A.7}$$

where $g = n_{row} n_{col}$ is the total number of windows in the grid. We shall denote the above as the distribution that is intrinsic to the image.

The distribution,

$$P(m = M_{i,j}) = G_{i,j},\ \forall i,j \tag{A.8}$$

is the distribution relative to a person's gaze map, $G$, where $\sum_{i,j} G_{i,j} = 1$. Given a gaze map $G$ we can also define a person's 'not looking' or complement uniform gaze map $\tilde{G}_{i,j}$ to be:

$$\tilde{G}_{i,j} := \frac{\mathbb{1}(G_{i,j} = 0)}{\sum_{l,m} \mathbb{1}(G_{l,m} = 0)},\ \forall i,j, \tag{A.9}$$

where $\mathbb{1}$ is the indicator function and the denominator gives the total number of grid squares in which the gaze map $G$ is zero. Hence, $P(m = M_{i,j}) = \tilde{G}_{i,j},\ \forall i,j$, is the distribution of where the person was not looking.

In our analysis, we compare the cumulative distribution functions of the above three types of probability distributions. As these CDFs are derived from data, we shall refer to them as Empirical Cumulative Distribution Functions (ECDFs).

Note that for some $(i,j) \neq (k,l)$, it may be that $M_{i,j} = M_{k,l}$ in the above definitions. The intrinsic distribution can be considered the distribution relative to the uniform gaze $\sum_{i,j} U_{i,j} = 1$ that looks equally at all parts of the image, like a scanner rather than a human viewer of art.

The window size used for feature maps in this study was 51 × 51, $H \times W$. Other sizes were also tested, yielding similar results - as shown in Supplementary Fig. Supp.B.13.

#### A.5 Operations on ECDFs

Given a (composite) topological descriptor $M$ and a gaze map $G$, we denote the probability distribution defined in Equation A.8, $\mathcal{D}(M,G)$, and the distribution intrinsic to the image defined in Equation A.7 as $\mathcal{D}(M,U)$. The corresponding ECDFs will be denoted by $EDCF(M,G)$ and $ECDF(M,U)$.

Given an image $k$ with a feature map $M_k$, a subject $s$ with gaze distribution $G_s$, we can compare the distributions using their ECDFs by computing mean error (ME) and mean squared error (MSE) in the usual way as follows:

| Feature map matrix | Definition of matrix element |
|---|---|
| $M^F_{density}$ | number of distinct cycles per grid square |
| $M^F_{persistence}$ | maximum persistence of cycles that occur within a grid square |
| $M^F_{perimeter}$ | maximum perimeter length of cycles that occur within a grid square |
| Combined Density $M_{density}$ | $M_{density} := M^{BW}_{density} + M^{WB}_{density}$ |
| Maximum Persistence $M_{persistence}$ | $M_{persistence} := \max(M^{BW}_{persistence}, M^{WB}_{persistence})$ |
| Maximum Perimeter $M_{perimeter}$ | $M_{perimeter} := \max(M^{BW}_{perimeter}, M^{WB}_{perimeter})$ |

Table 6: Feature maps, where $F \in \{BW, WB\}$ is the filtration.

$$ME_{k,s} = \frac{1}{X} \sum_{x=x_{min}}^{x=x_{max}} ECDF(M_k, U)(x) - ECDF(M_k, G_s)(x), \tag{A.10}$$

where $x \in [x_{min}, x_{max}]$, $X = x_{max} - x_{min}$ is the range of values of the topological descriptor for which $0 \leq ECDF(M_k, U)\,(x) < 1$, in other words, the width of the curve, $ECDF(M_k, U)$.

Analogously the Mean Squared Error is given by:

$$MSE_{k,s} = \frac{1}{X} \sum_{x=x_{min}}^{x=x_{max}} (ECDF(M_k, U)(x) - ECDF(M_k, G_s)(x))^2. \tag{A.11}$$

The full set of ECDFs, for each image $k$, feature map $M$ and participant $s$, $ECDF(M_k, U)$ ('intrinsic'), $ECDF(M_k, G_s)$ ('looking'), $ECDF(M_k, \tilde{G_s})$ ('not looking') can be found in Supplementary section B.7.

### B Supplementary results

#### B.1 Exhibition images

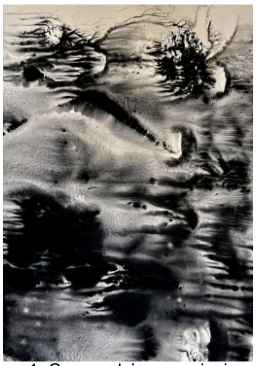

1. Czarne dziury pamięci
(eng. “Black holes of blackness”)

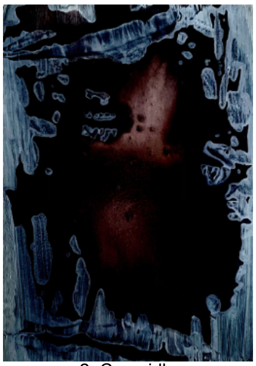

2. Czernidło
(eng. “Black wash”)

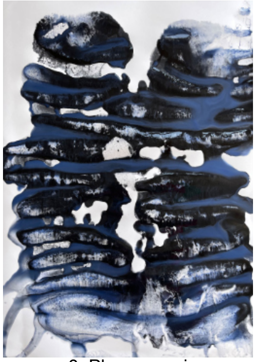

3. Płuca czerni
(eng. “Lungs of blackness”)

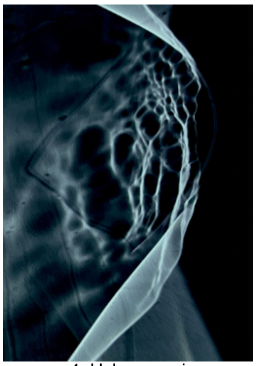

4. Ucho czerni
(eng. “Ear of blackness”)

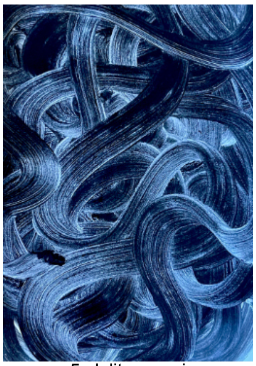

5. Jelita czerni
(eng. “Guts of blackness”)

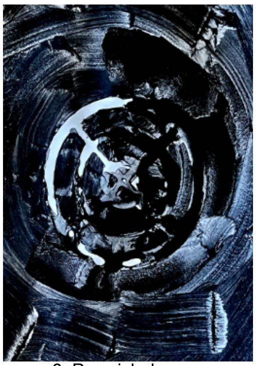

6. Przycisk do serc
(eng. “Hearts button”)

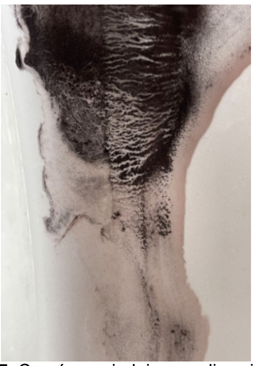

7. Czerń na miednicy emaliowej
(eng. “Blackness on enamel washing basin”)

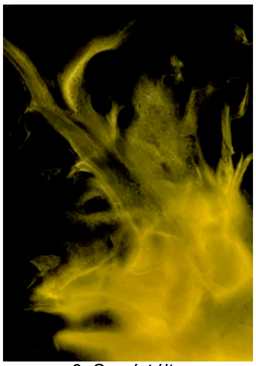

8. Czerń żółta
(eng. “Yellow blackness”)

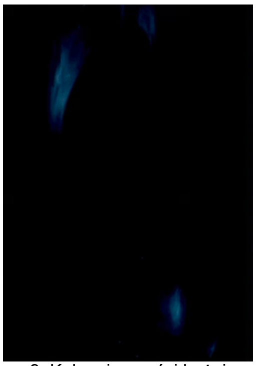

9. Kolor ciemności bożej
(eng. “The colour of holy blackness”)

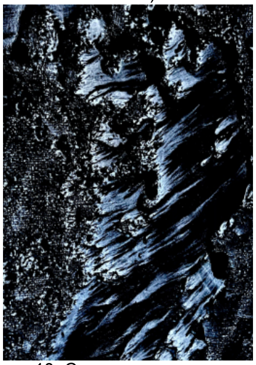

10. Czarne na czarnym
(eng. “Black on black”)

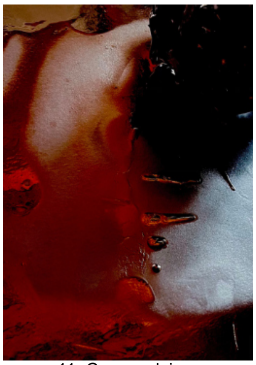

11. Czarna dziura
(eng. “Black hole”)

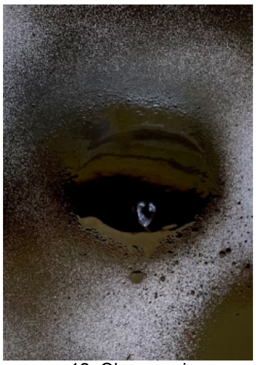

12. Oko czerni
(eng. “Eye of blackness”)

Figure Supp.B.1: All artistic images [Kot, 2022]. Original tiles are given under each image

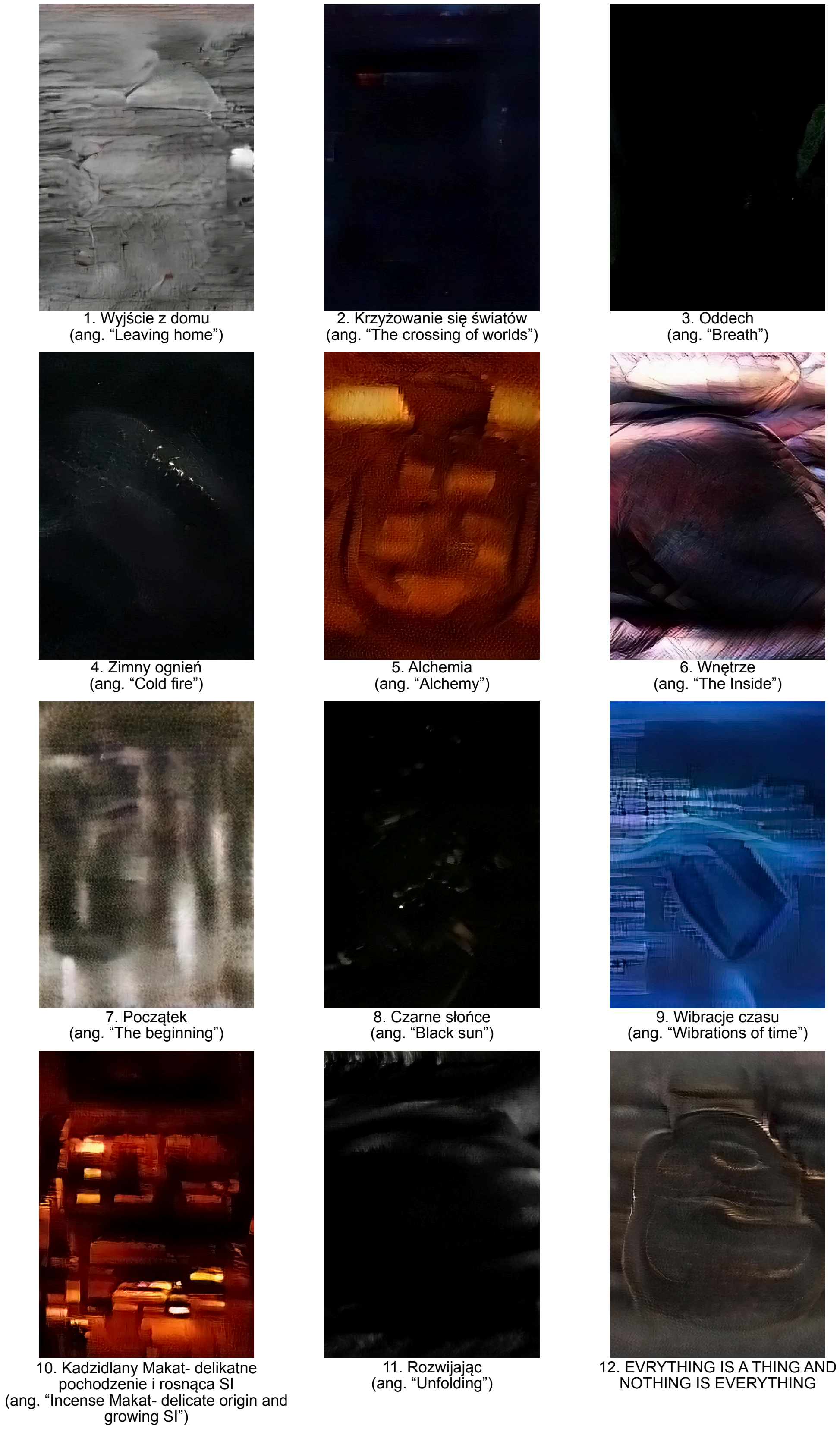


Figure Supp.B.2: All pseudo-artistic images. Original tiles are given under each image.

#### B.2 Visualisation of topological analyses for all investigated images

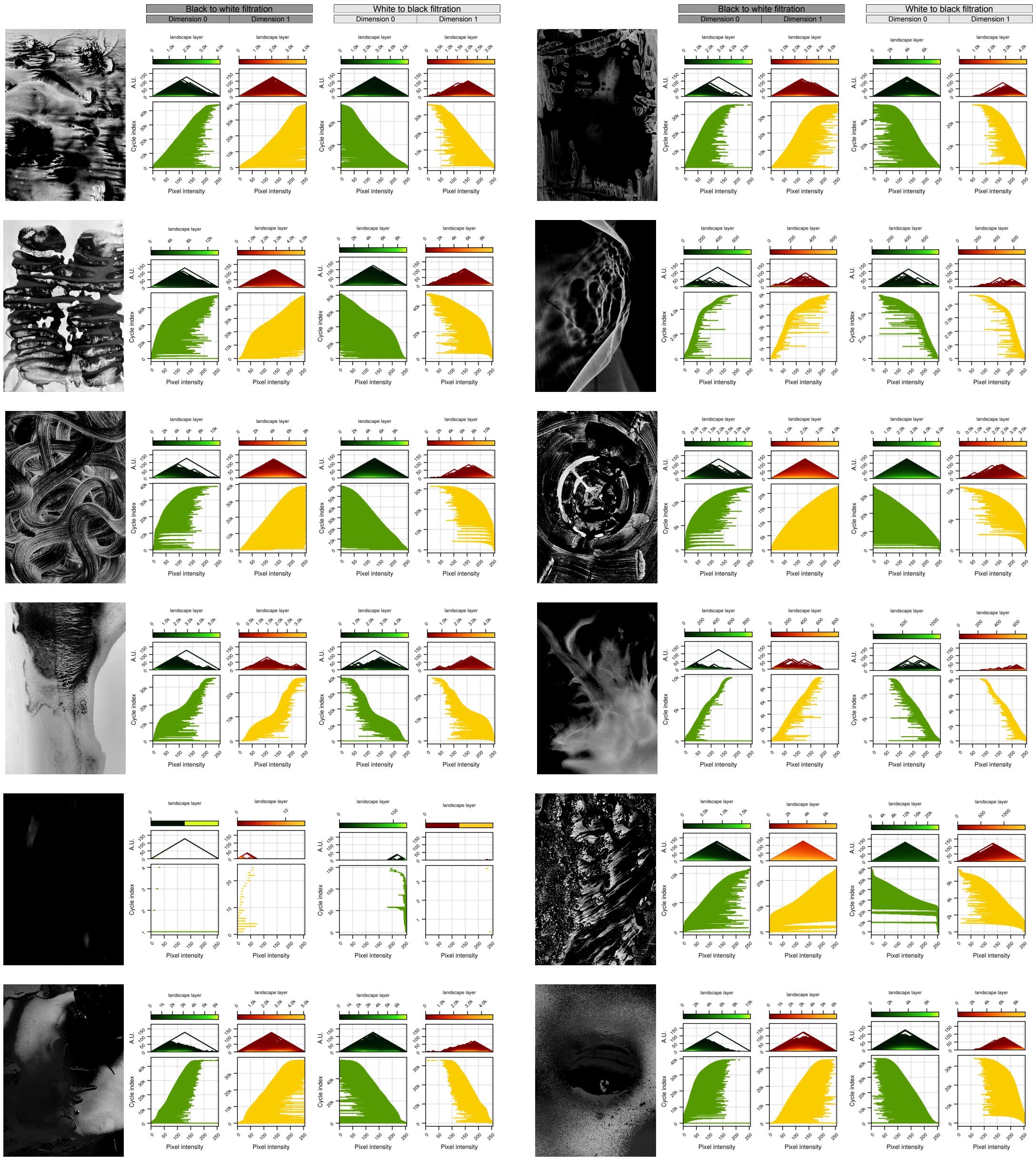


Figure Supp.B.3: Persistence barcodes and landscapes for the artistic [Kot, 2022] images (BW filtration).

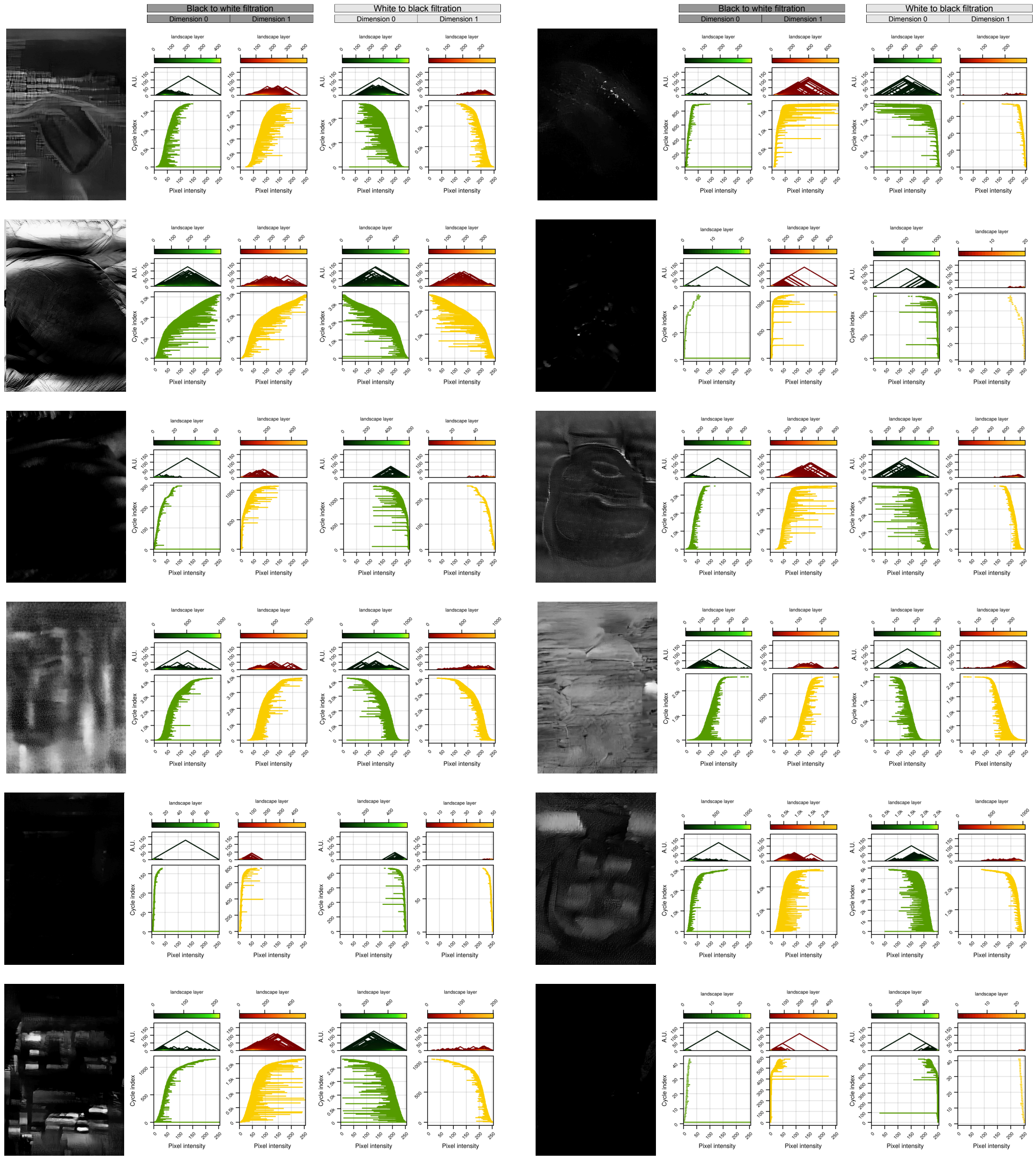


Figure Supp.B.4: Persistence barcodes and landscapes for the Pseudo-artistic images (BW filtration).

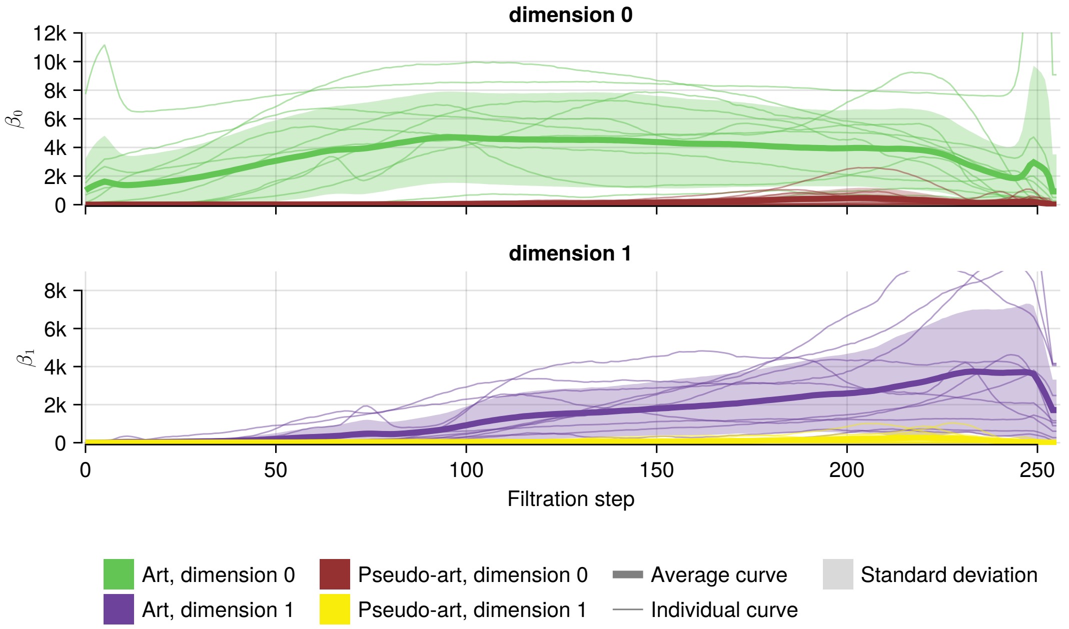


Figure Supp.B.5: Average Betti curves with their standard deviation, together with individual Betti curves for each image for the WB filtration. The average Betti curve was computed for each group: artistic and pseudo-artistic images, with results for dimensions 0 shown in the top row and dimension 1 in the bottom row.

#### B.3 Filtration duality for investigated images

##### B.3.1 Toy example of duality

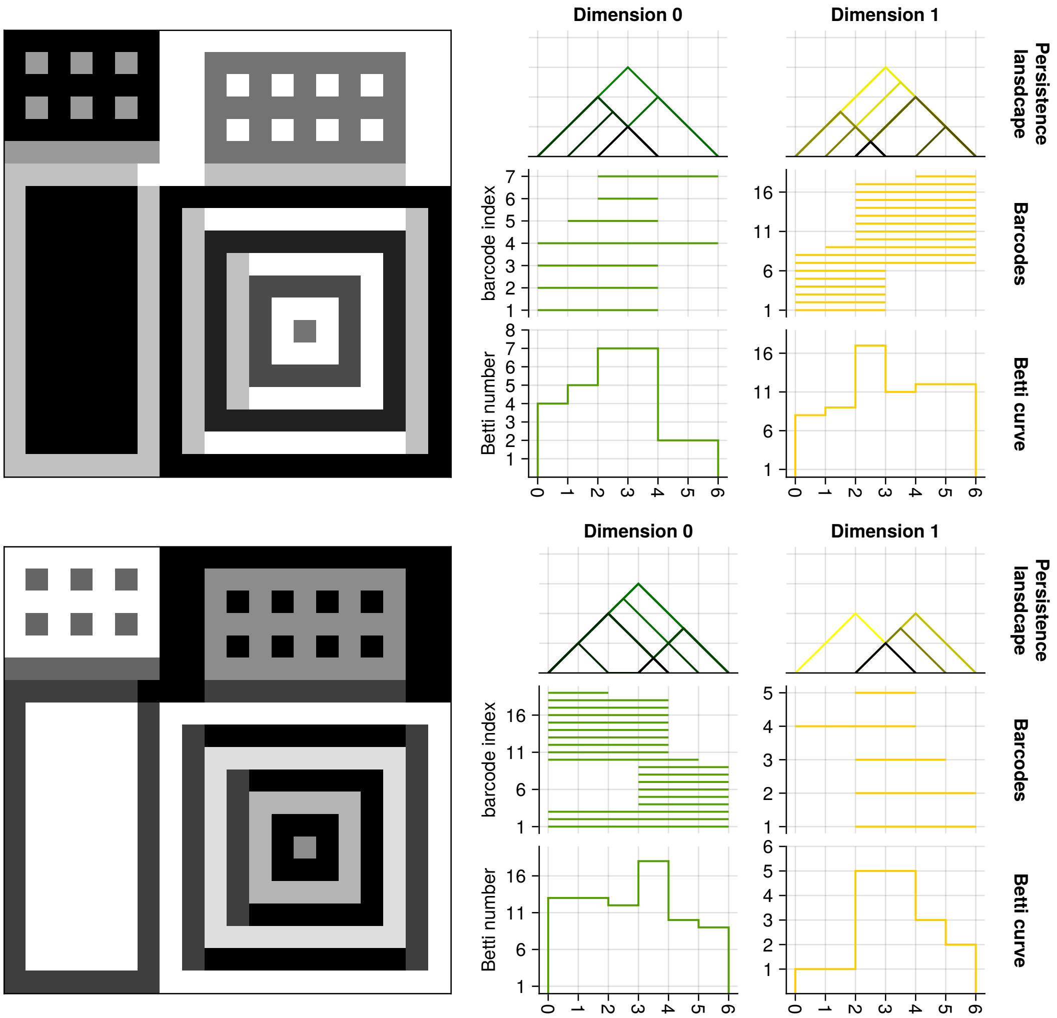


Figure Supp.B.6: An example of duality how duality affects cycles and their lifetime. The image used for black-to-white filtration is presented in the first column, top row; the dual filtration is done with the inverse of the original image, with the filtration again going from black to white (this was done so that it is easier to notice starting points, while the results of the filtration are the same after inverting the horizontal axis on the plots). For both images, results are presented in dimensions 0 and 1 in the form of persistence landscapes, barcodes and Betti curves, as indicated with the labels. The results for dimension 1, top row and dimension 0, bottom row, are the same with respect to inverting the horizontal axis. The results for dimension 0, top row, have 2 more cycles, one is the infinite cycle (which contributes to the outermost layer in the landscape), second missing cycle is related to one of the barcodes starting at step 0 and finishing at step 4 (dimension 0, top filtration).

##### B.3.2 L1 distance comparison for investigated images

To justify the sufficiency of only selecting dimension 1 cycles for our analysis, we computed $L^1$ pairwise distance between all 24 persistence landscapes within each dimension and for both filtrations. The duality between the topological invariants derived from the two filtration predicts (up to boundary cycles) that these $L^1$ distance matrices will be almost identical. Fig. Supp.B.7 verifies the expected similarity between the following pairs of matrices: ((1) BW filtration in dimension 0 and WB filtration in dimension 1; (2) BW filtration in dimension 1 and WB filtration in dimension 0). This is a powerful demonstration of the duality.

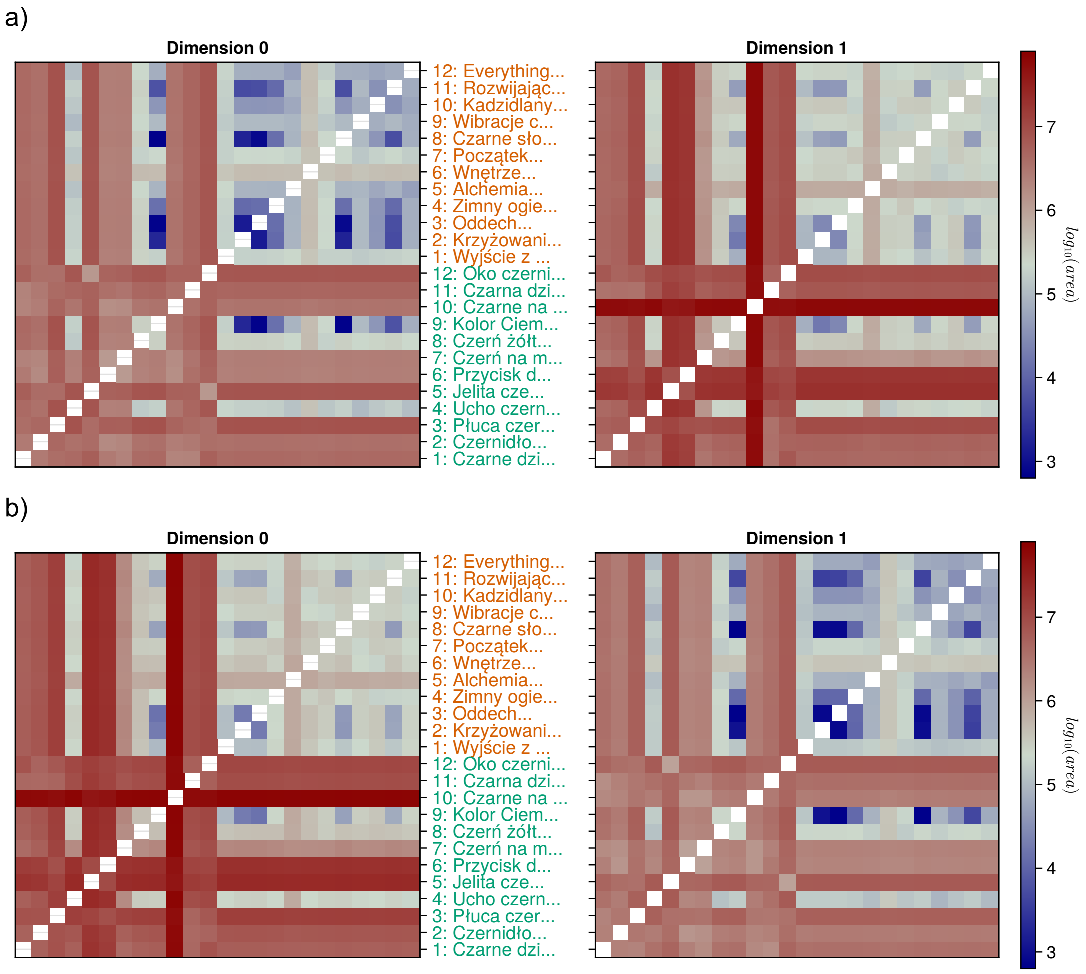


Figure Supp.B.7: Comparison of $L^1$ distance matrices between filtration and demonstration of duality: a) filtration from black to white; b) filtration from white to black. Each heatmap shows $L^1$, pairwise differences between the persistence landscapes of all images in dimensions 0 (left) and 1 (right). Every row (or column) in the heatmap corresponds to an image annotated between the matrices. The top 12 labels (coloured orange) are from Pseudo-artistic images, and the next 12 labels (coloured green) are from Artist images. Please refer to Supplementary Fig. Supp.B.1 and Supp.B.2 for full names of images. While not all cycles are reflected in the duality, the overall shape differences captured with $L^1$ distance between landscapes are mirrored when the filtration of the images is inverted.

#### B.4 The effect of image resizing.

All images used in the study were 1434 pixels wide and 2048 pixels tall. Since the choice of image size was arbitrary, we tested how much topological properties changed with the change in size of the images.

The effect of changing image size on the area under the persistence landscapes for the 24 images (as shown in the caption) is shown in the area-under-landscape space, where the $x$ and $y$ coordinates are areas under the landscape in dimensions 0 and 1, respectively.

As shown in Fig. Supp.B.8, all of the images from both exhibitions were upscaled or downscaled while preserving the image's aspect ratio (image sizes are shown in the upper part of each legend). The upscaling of the images did not change the topological properties significantly- for both groups, the markers occupy the same space. For downscaling, however, the area under the persistence landscape for both data sets is decreasing. It is important to note that at every level of resizing, the relative location of both datasets in the area area-under-landscape space was preserved- the artistic images have higher area-under-landscape than the pseudo-artistic images (except for 3 cases- image number 9 being significantly lower than any other image, and images 4 and 8 being very close to the pseudo-artistic images).

It should be noted that for most of the images, downsizing by a factor of 4 (resulting in image size 512 × 359) did not change the area-under-landscape by less than one order of magnitude.

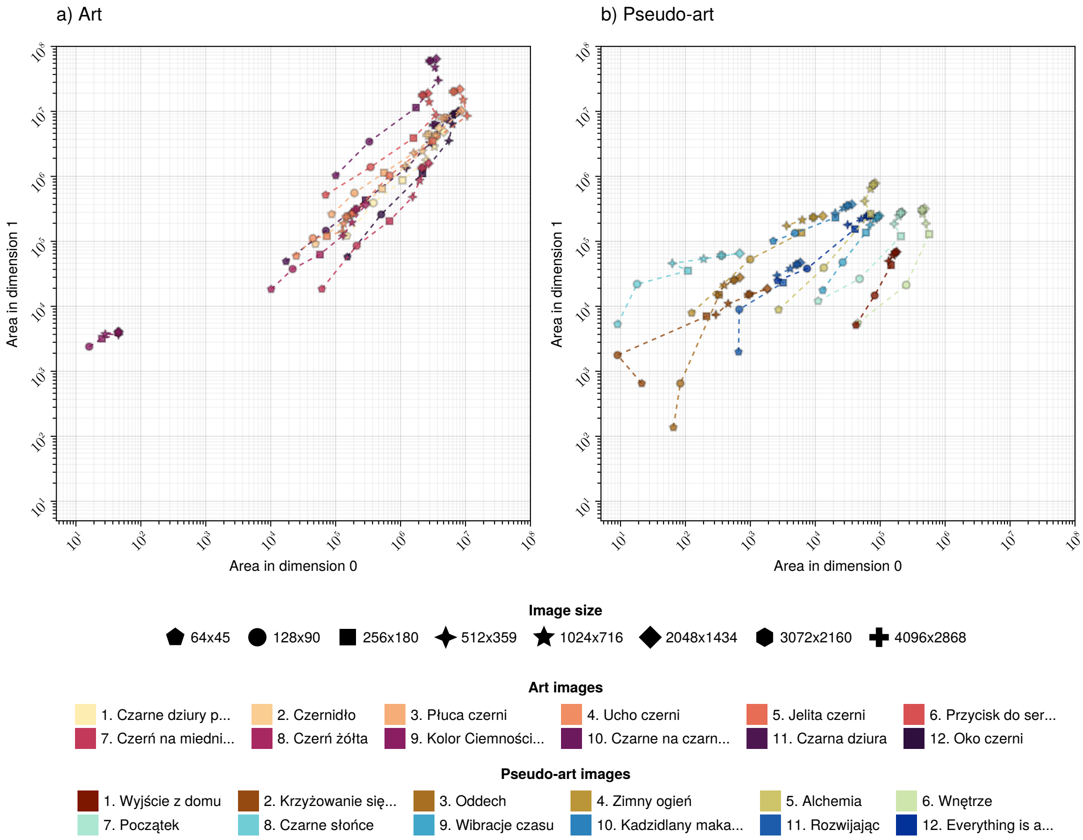


Figure Supp.B.8: Effect of rescaling the images on the persistence features, shown as the trajectory of changes in areas under the landscapes in dimensions 0 and 1, a) artistic images, b) pseudo-artistic images. Persistence properties were computed for each image (marked with different colours) after resizing (size indicated by markers), creating "landscape trajectories".

#### B.5 Cycle visualisation

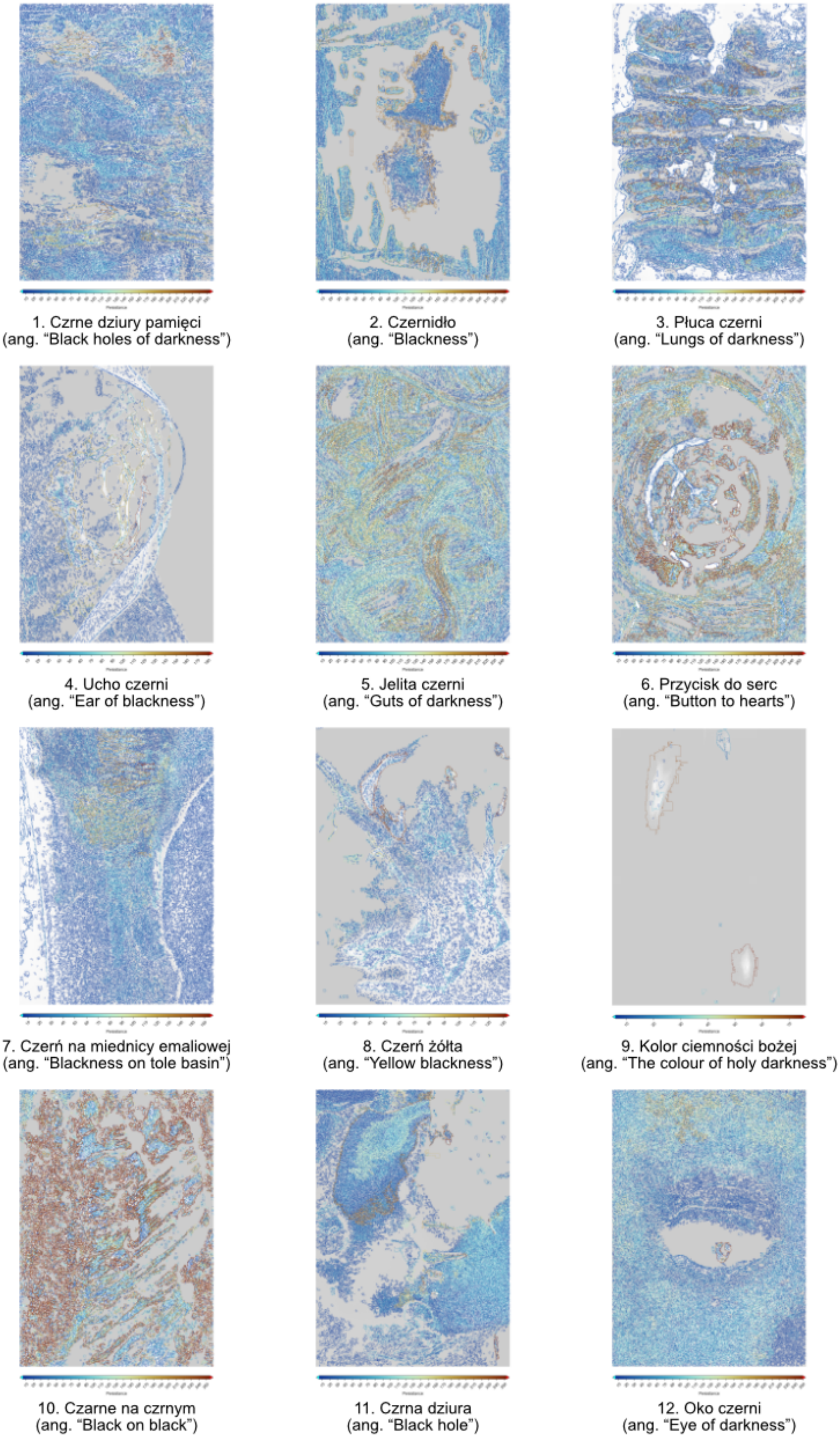


Figure Supp.B.9: Visualisation of all cycles in dimension 1, artistic images [Kot, 2022].

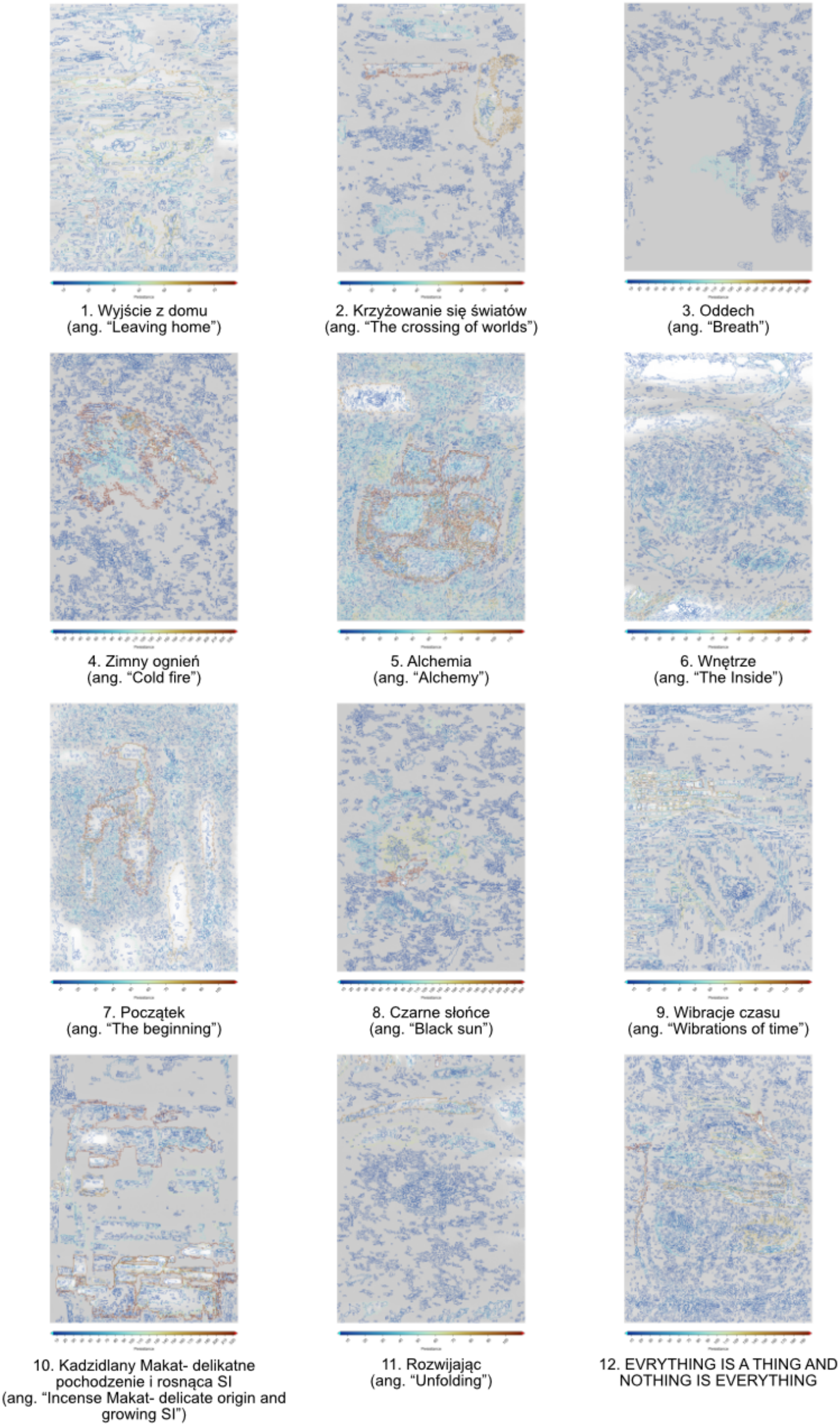


Figure Supp.B.10: Visualisation of all cycles in dimension 1, pseudo-artistic images.

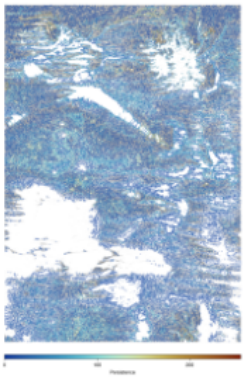

1. Czrne dziury pamięci
(ang. “Black holes of darkness”)

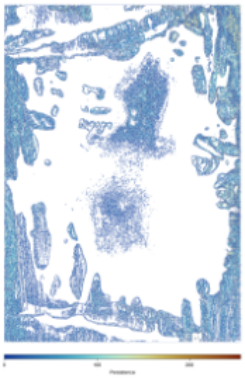

2. Czernidło
(ang. “Blackness”)

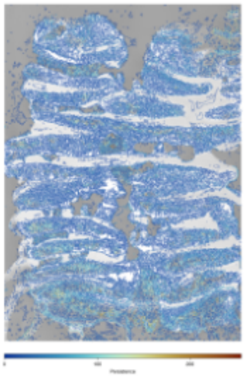

3. Płuca czerni
(ang. “Lungs of darkness”)

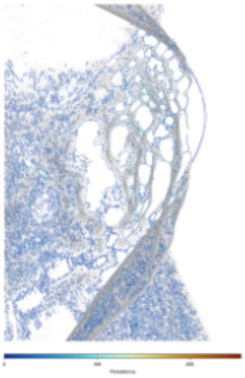

4. Ucho czerni
(ang. “Ear of blackness”)

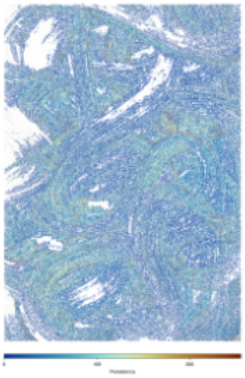

5. Jelita czerni
(ang. “Guts of darkness”)

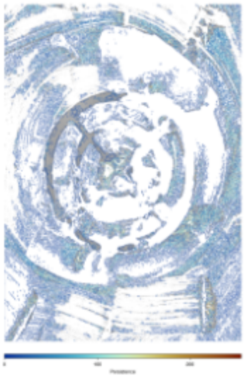

6. Przycisk do serc
(ang. “Button to hearts”)

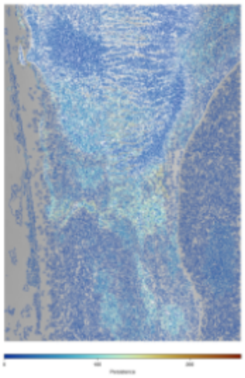

7. Czerń na miednicy emaliowej
(ang. “Blackness on tole basin”)

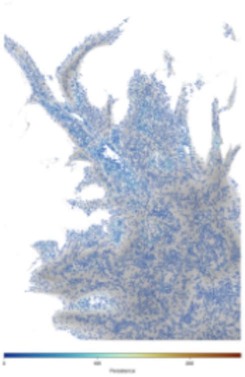

8. Czerń żółta
(ang. “Yellow blackness”)

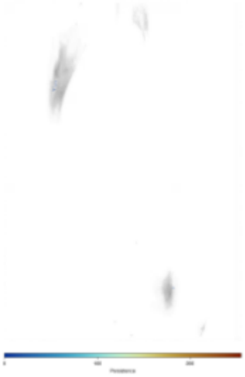

9. Kolor ciemności bożej
(ang. “The colour of holy darkness”)

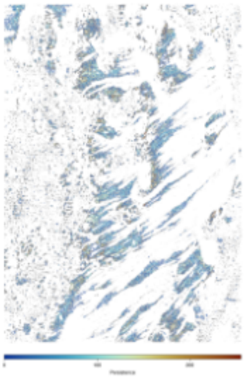

10. Czarne na czrnym
(ang. “Black on black”)

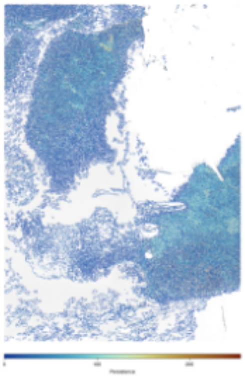

11. Czrna dziura
(ang. “Black hole”)

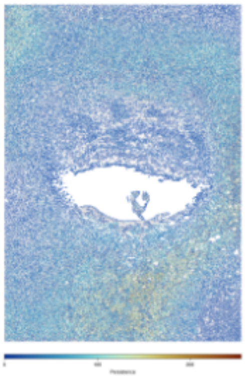

12. Oko czerni
(ang. “Eye of darkness”)

Figure Supp.B.11: Visualisation of all cycles in dimension 1, artistic images [Kot, 2022] with filtration from white to black colours.

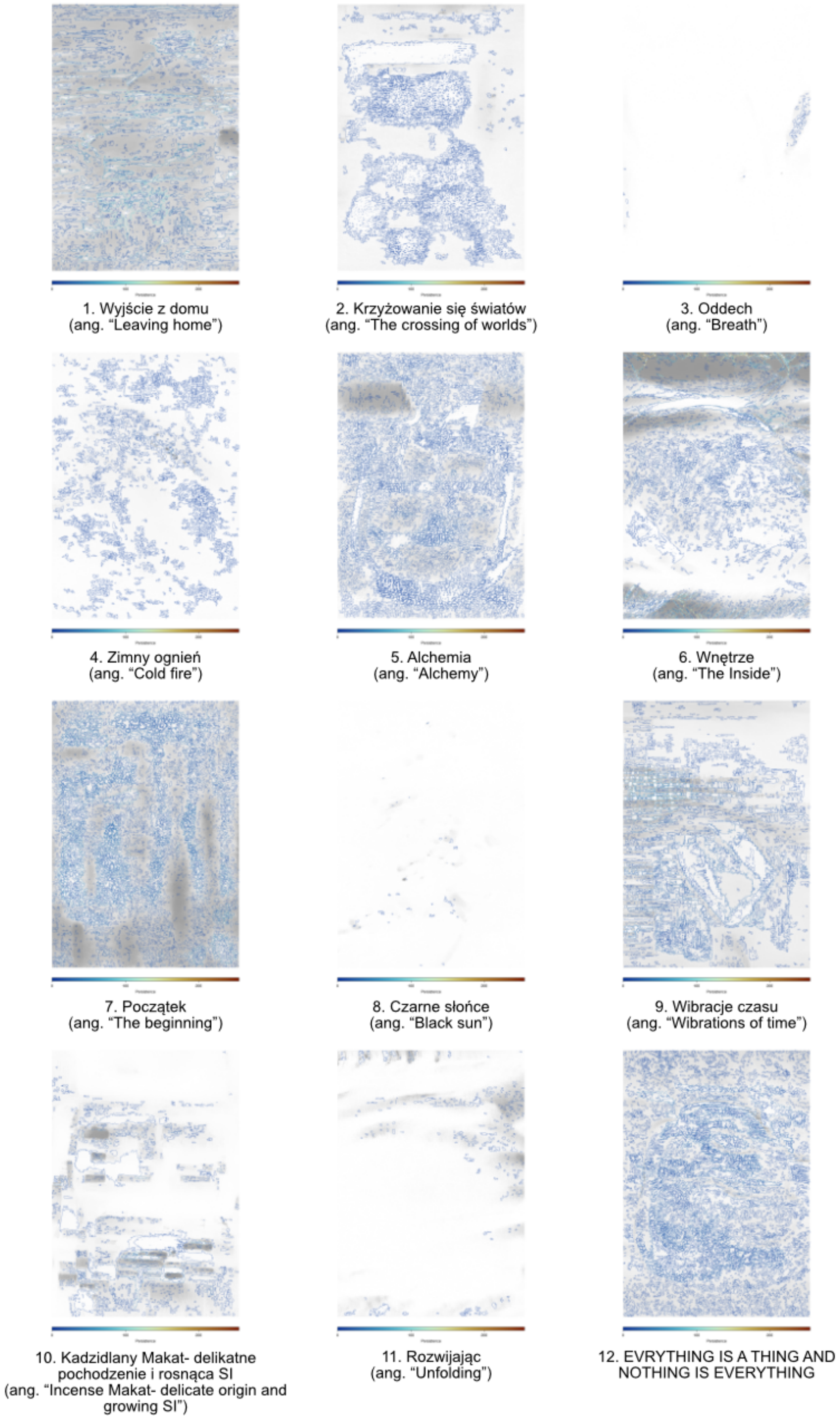


Figure Supp.B.12: Visualisation of all cycles in dimension 1, pseudo-artistic images with filtration from white to black colours.

#### B.6 Grid window size comparison for ECDFs

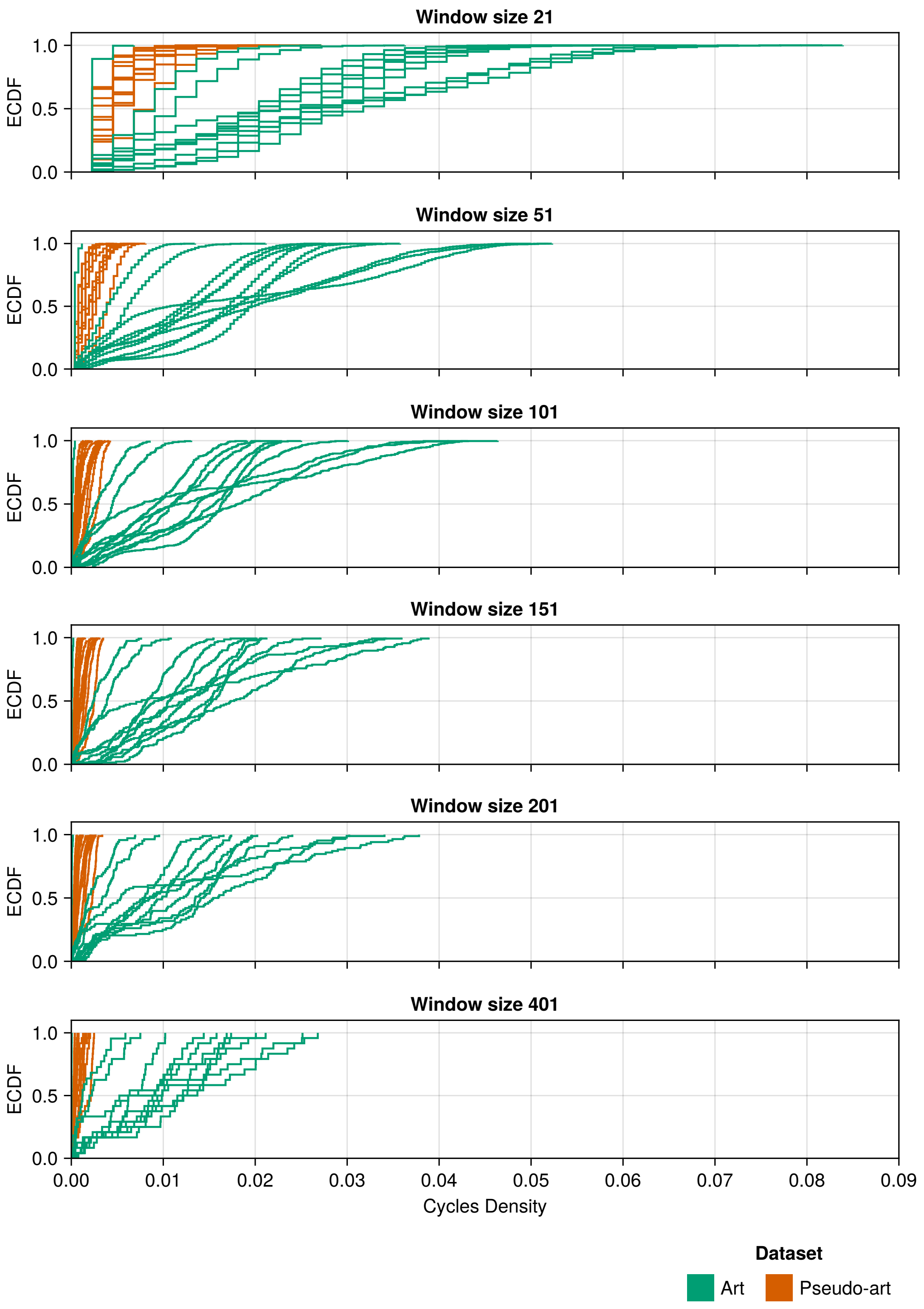


Figure Supp.B.13: Cycle density intrinsic ECDF curves, $ECDF(M_{density}, U)$, for the artistic images (green) and the pseudo-artistic images (orange) for different window sizes (plots from top to bottom): 21, 51, 101, 151, 201, 401.

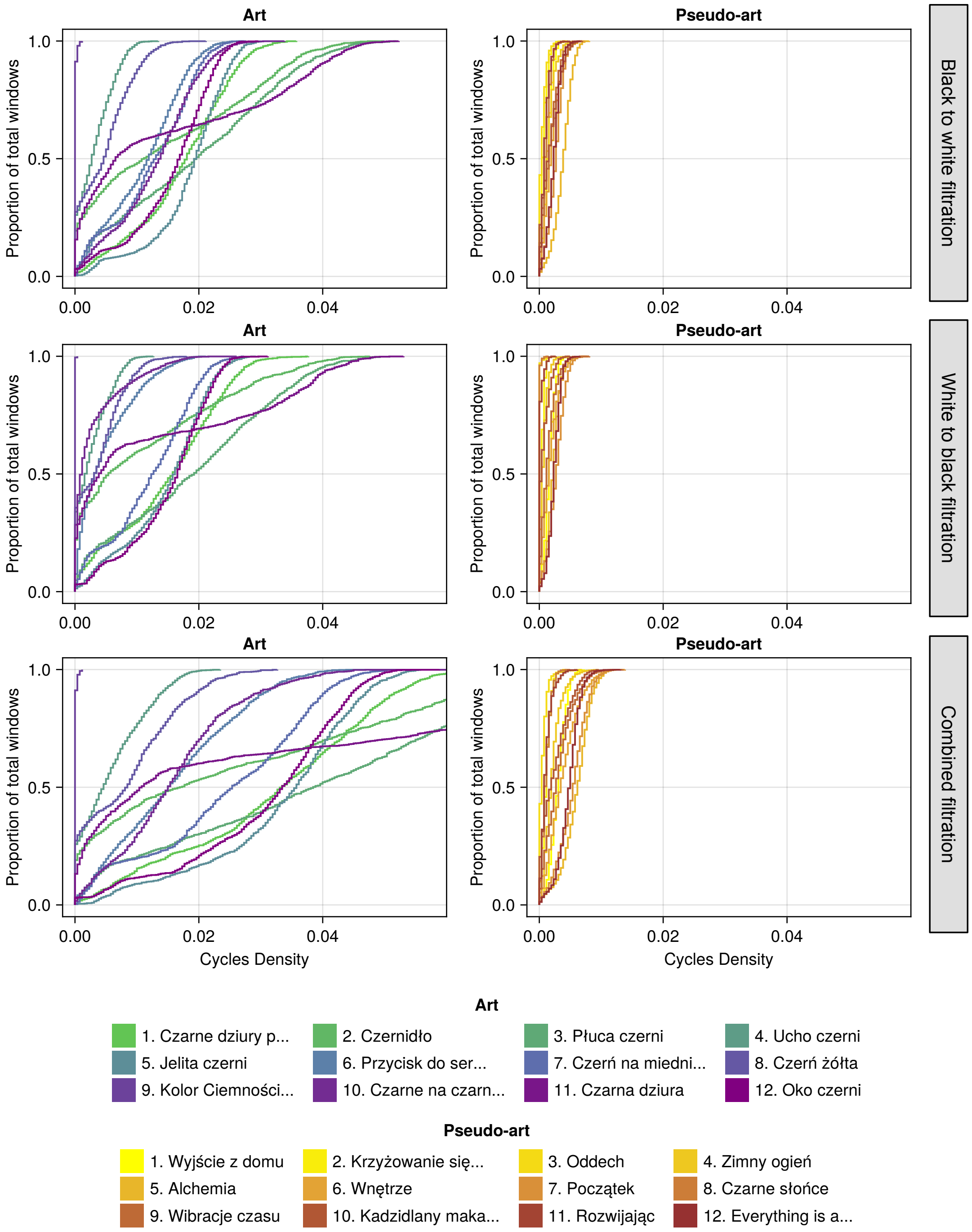


Figure Supp.B.14: Intrinsic ECDF curves for cycle density feature map, $ECDF(M_{density}, U)$, for: BW filtration (top row), WB filtration (middle row), combined BW and WB filtration (bottom row).

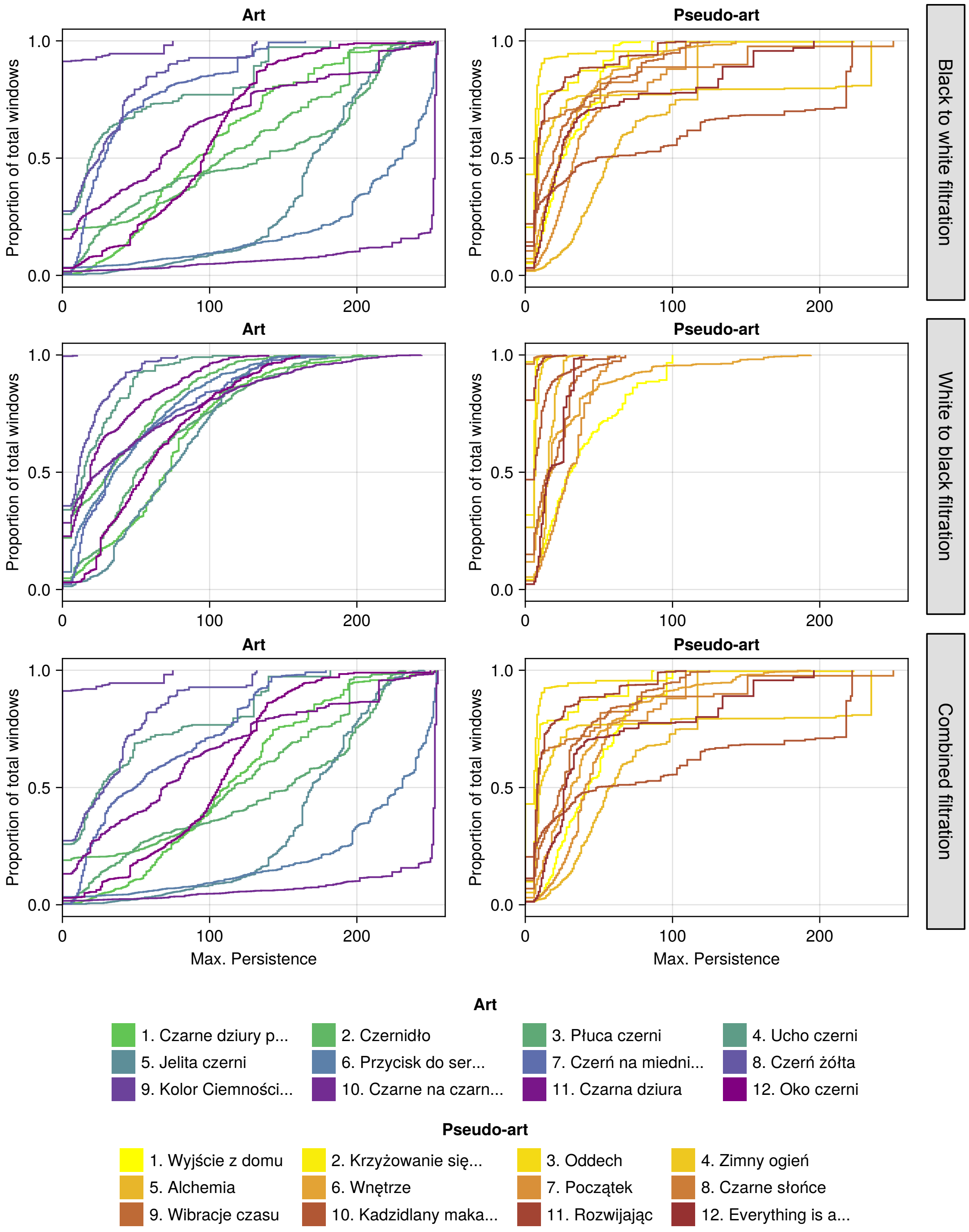


Figure Supp.B.15: ECDF curves for maximal persistence feature map, $ECDF(M_{persistence}, U)$, for: BW filtration (top row), WB filtration (middle row), combined BW and WB filtration (bottom row).

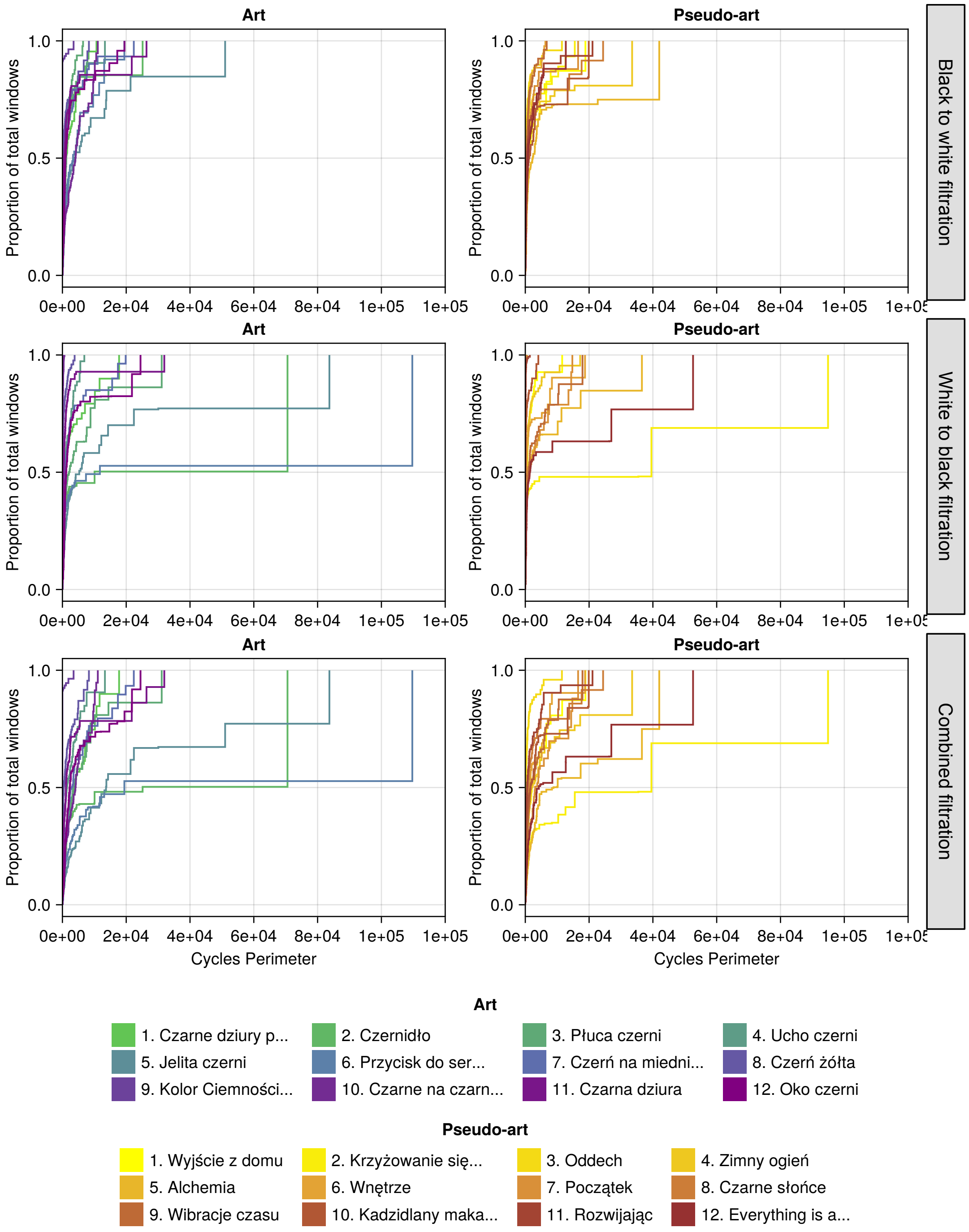


Figure Supp.B.16: Intrinsic ECDF curves for cycle perimeter feature map, $ECDF(M_{perimeter}, U)$, for: BW filtration (top row), WB filtration (middle row), combined BW and WB filtration (bottom row).

#### B.7 Complete set of ECDFs for 'looking' and 'not looking'

Results for the full set of ECDFs. For each image $k$, feature map $M$ and participant $s$, we have $ECDF(M_k, U)$ ('intrinsic'), $ECDF(M_k, G_s)$ (where the participant was 'looking'), $ECDF(M_k, \tilde{G_s})$ (where the participant was 'not looking').

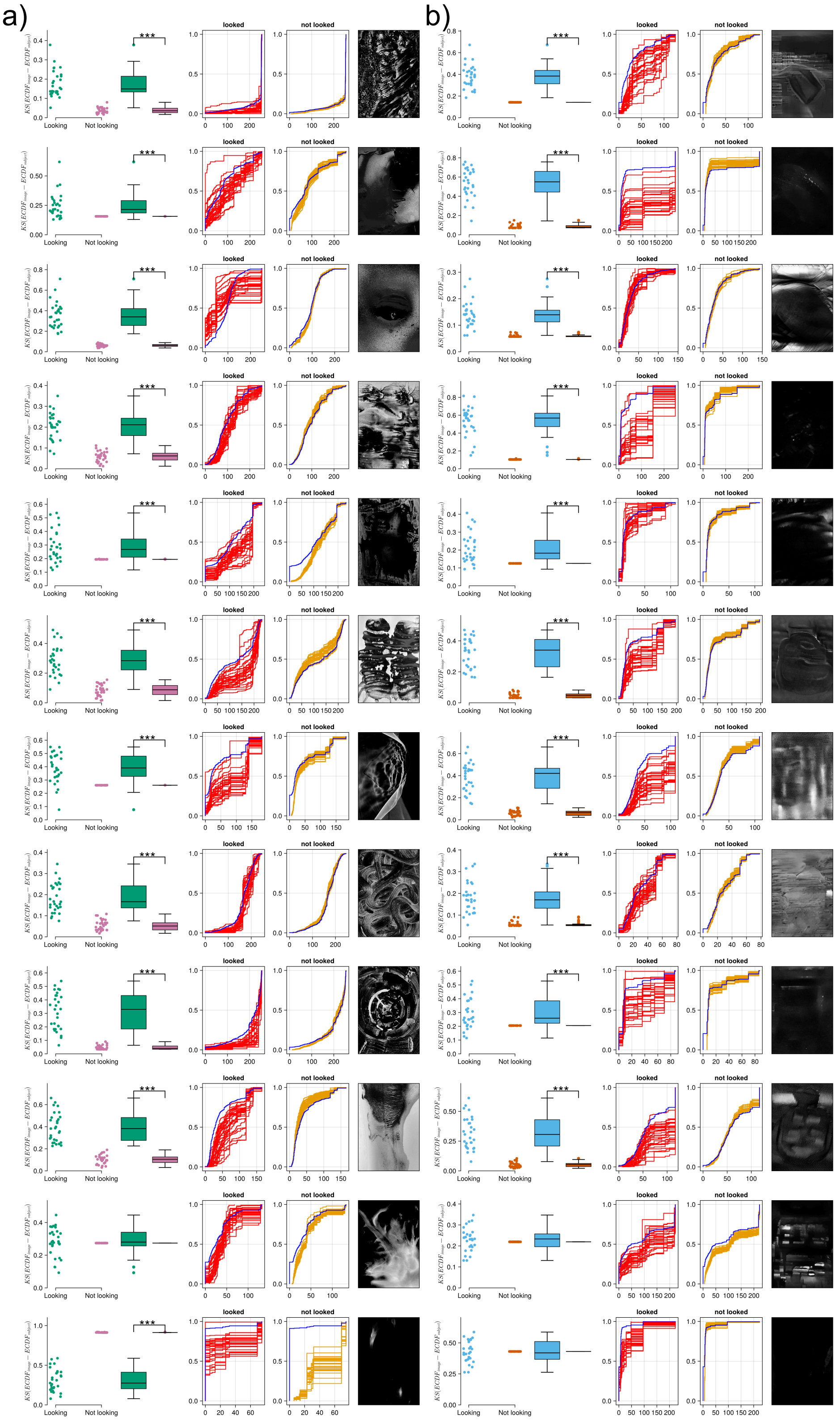

Figure Supp.B.17: ECDFs for maximal persistence for each image and each participant for both (a) art [Kot, 2022] and (b) pseudo art, for both sessions. Within a column the plots are (from left to right): scatterplot of the Kolmogorov-Smirnov statistic between the image's ECDF vs 'looking' ECDF and images's ECDF vs 'not looking' ECDF for each person; bar plot of the same; the 'Looking ECDF' for each person; 'Not looking ECDF' for each person; the image itself. The blue ECDF is intrinsic to the image itself and can be thought to be arising from a 'gaze' that is a uniform scan of the entire image.

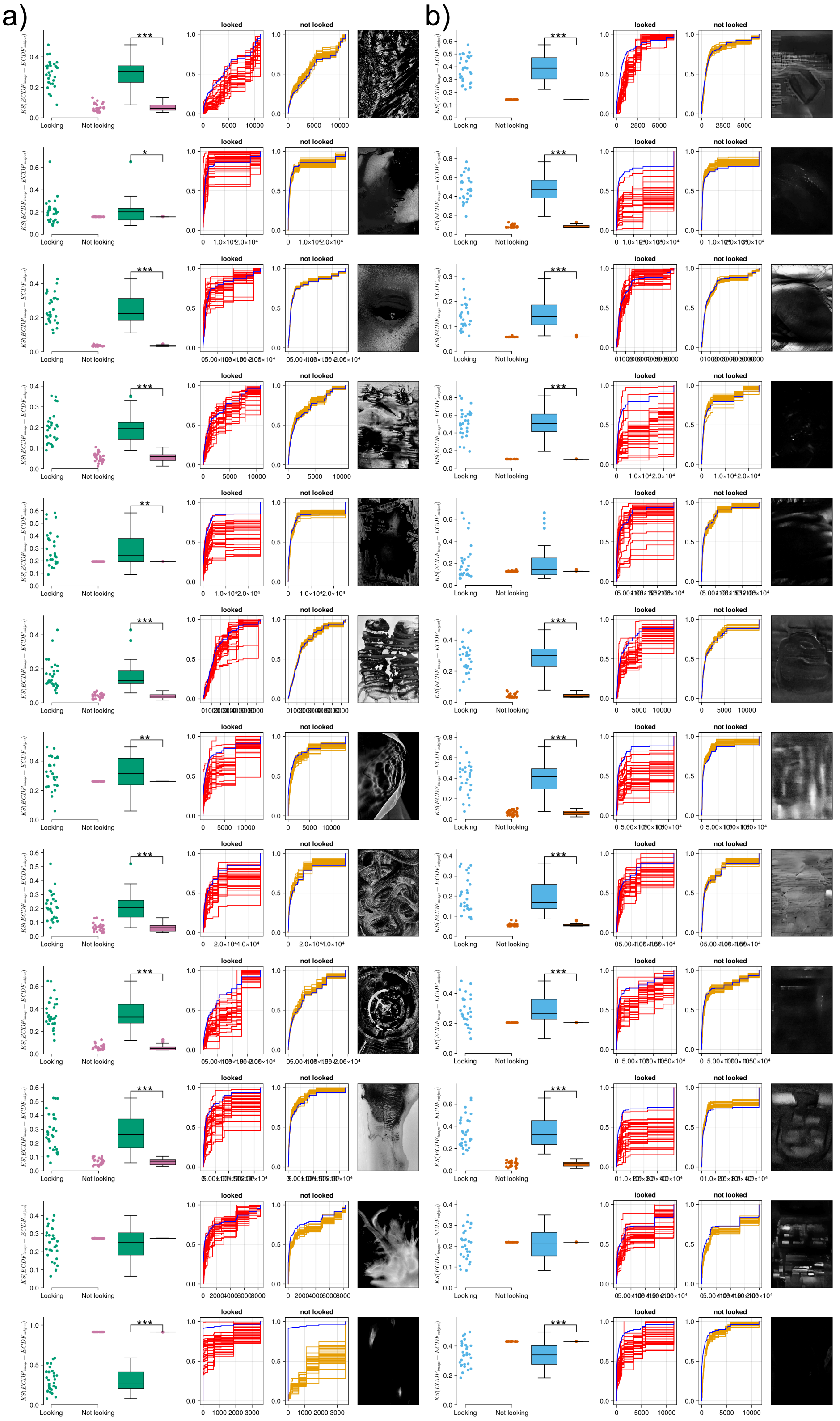


Figure Supp.B.18: ECDFs for cycle perimeter for each image and each participant for both (a) art [Kot, 2022] and (b) pseudo art, for both sessions. Within a column the plots are (from left to right): scatterplot of the Kolmogorov-Smirnov statistic between the image's ECDF vs 'looking' ECDF and images's ECDF vs 'not looking' ECDF for each person; bar plot of the same; the 'Looking ECDF' for each person; 'Not looking ECDF' for each person; the image itself. The blue ECDF is intrinsic to the image itself and can be thought to be arising from a 'gaze' that is a uniform scan of the entire image.

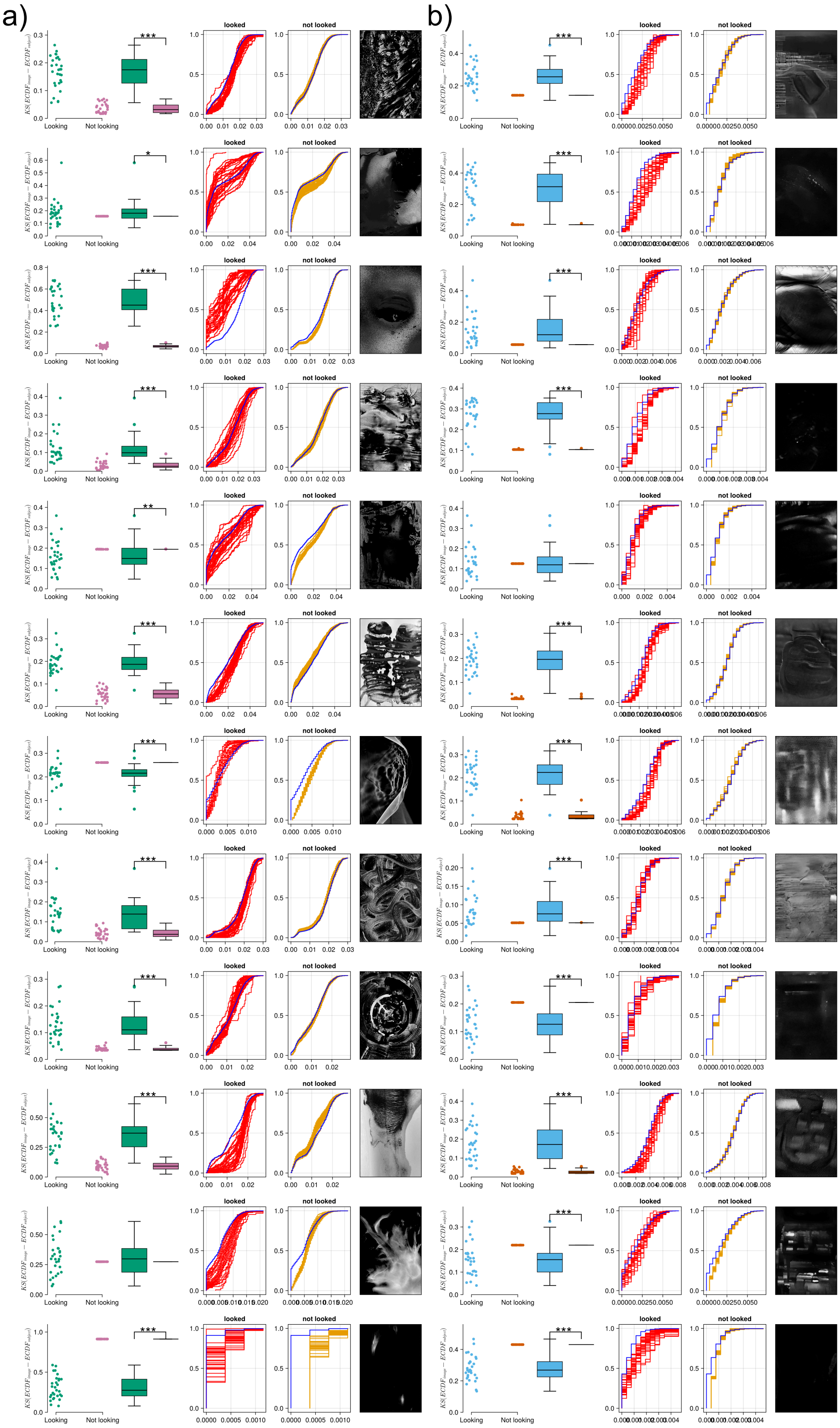

Figure Supp.B.19: ECDFs for cycle density for each image and each participant for both (a) art [Kot, 2022] and (b) pseudo art, for both sessions. Within a column the plots are (from left to right): scatterplot of the Kolmogorov-Smirnov statistic between the image's ECDF vs 'looking' ECDF and images's ECDF vs 'not looking' ECDF for each person; bar plot of the same; the 'Looking ECDF' for each person; 'Not looking ECDF' for each person; the image itself. The blue ECDF isintrinsic to the image itself and can be thought to be arising from a 'gaze' that is a uniform scan of the entire image.
